# Characterization of tomato *canal-1* mutant using a multi-omics approach

**DOI:** 10.1101/2023.11.26.567847

**Authors:** Micha Wijesingha Ahchige, Josef Fisher, Ewelina Sokolowska, Rafe Lyall, Nicola Illing, Aleksandra Skirycz, Dani Zamir, Saleh Alseekh, Alisdair R. Fernie

## Abstract

The recently described *canal-1* tomato mutant, which has a variegated leaf phenotype, has been shown to affect canalization of yield. The corresponding protein is orthologous to AtSCO2 - SNOWY COTYLEDON2, which has suggested roles in thylakoid biogenesis. Here we characterize the *canal-1* mutant through a multi-omics approach, by comparing mutant to wild-type tissues. While white *canal-1* leaves are devoid of chlorophyll, green leaves of the mutant appear wild-type-like, despite an impaired protein function. Transcriptomic data suggest that green mutant leaves compensate for this impaired protein function by upregulation of transcription of photosystem assembly and photosystem component genes, thereby allowing adequate photosystem establishment, which is reflected in their wild-type-like proteome. White *canal-1* leaves, however, likely fail to reach a certain threshold enabling this overcompensation, and plastids get trapped in an undeveloped state, while additionally suffering from high light stress, indicated by the overexpression of ELIP homolog genes. The metabolic profile of white and to a lesser degree also green tissues revealed upregulation of amino acid levels, that was at least partially mediated by transcriptional and proteomic upregulation. These combined changes are indicative of a stress response and suggest that white tissues behave as carbon sinks. In summary, our work demonstrates the relevance of the SCO2 protein in both photosystem assembly and as a consequence in the canalization of yield.

**Significance statement:** The variegated *canalized-1* tomato mutant was recently described and the underlying gene *SCO2* suggested to be a yield canalization gene. Through a multi-omics approach we show that mutants require a transcriptional upregulation of photosystem components and assembly components, likely as overcompensation for partially impaired SCO2 function, to produce a wild type-like proteome and functional photosynthetic tissue Our data, furthermore, suggest that variation of green to white leaf area from plant to plant leads to the yield variation.

## Introduction

Crop yield remains one of the major focal points in plant breeding to feed a growing population. Additionally, in the face of increasing environmental variability, it seems necessary to utilize plant breeding to decrease the variability of crop performance. Many studies have investigated mean yield levels in different crop species and found genetic regions or individual genes that affect crop yield. That said the stability of yield has also been long understood to be desirable with this being the focus of studies in barley by Finlay and Wilkinson in 1963 (Finlay and Wilkinson, 1963) Such studies were extended to include oats and maize (Becker and Léon, 1988) and more recently a wide variety of further species including tomato, eggplant, pepper, melon, watermelon, and sunflower (Fisher *et al*., 2017). While tomato is not one of the main crops responsible for providing the majority of calories for human consumption, it does play an important role in many diets. It is, furthermore, regarded as a model plant for other fruit-bearing crops (Kimura and Sinha, 2008), as well as being an excellent genetic model system for which a wide range of genomic and post-genomic resources are available (Becker and Léon, 1988, Klee and Giovannoni, 2011, Sato *et al*., 2012, Zhu *et al*., 2018, Gao *et al*., 2019). By utilization of different mapping populations, many quantitative trait loci have also been identified in tomato that are responsible for fruit yield (Eshed and Zamir, 1995, Fridman *et al*., 2004, Gur and Zamir, 2015, Ofner *et al*., 2016, Tieman *et al*., 2017, Zemach *et al*., under revision). However, much less is known about the genetic factors influencing the variation of yield from plant to plant, be it in tomato or other crops. A concept to describe the lack of this sort of variation is called canalization, which was coined by Waddington in 1942 (Waddington, 1942). This has mainly been used in a developmental context. Whilst early studies were generally confined to a limited number of genotypes this has not been the case recently. For example, studies of the developmental stability of Arabidopsis recombinant inbred line (RIL) populations and a population of wide geographic origin under different photoperiods resulted in the suggestion that ERECTA contributes to the canalization of rosette leaf number (Hall *et al*., 2007). A similar study looked at the genetic canalization of flowering time on *Boechera stricta* (Lee *et al*., 2014), while treatment of Arabidopsis ecotypes and RILs with pharmacological inhibitors of Hsp70 has been taken to suggest that it acts as a capacitor of phenotypic variation in plants (Queitsch *et al*., 2002). In addition to these developmental studies, the concept has recently been applied to describe metabolic quantitative traits (Alseekh *et al*., 2017, Wijesingha Ahchige *et al*., 2023).

The recent study mentioned above, using archived mapping data, found a bimodal distribution of stable and variable traits (Fisher *et al*., 2017). This study further confirmed this tendency of more and less variable traits, in a common crop garden experiment with many different varieties of crop plants. Further studies in tomatoes have found both loci with positive as well as negative effects on yield stability (Fisher and Zamir, 2021). They also identified a yield canalization mutant with variegated leaves, named *canal-1*. Field-grown plants can vary largely in plant size and accordingly show a strong variation in fruit yield. Fine mapping pinpointed the gene to Solyc01g108200, which is an orthologous gene to SNOWY COTYLEDONS 2 (SCO2/AT3G19220) in *Arabidopsis thaliana*. As the name suggests, *A. thaliana* mutants of this gene have snowy white cotyledons but develop regular green leaves under normal conditions (Shimada *et al*., 2007, Albrecht *et al*., 2008). Indeed, the *snowy cotyledon* mutants came out of a genetic screen that was aimed at identifying genes required for proper chloroplast development, particularly in cotyledons. The screen specifically selected mutants with pale cotyledons but normal green true leaves. As yet *sco1*, *sco2,* and *sco3* have been identified and characterized. *Sco1* was identified in a gene encoding the plastid elongation factor G (Albrecht *et al*., 2006). *Sco2* was identified as a chloroplastic DNA-J-like zinc finger protein that was essential for early seedling survival in Arabidopsis (Albrecht *et al*., 2008), as well as playing a role in leaf variegation in both Arabidopsis (Albrecht *et al*., 2008) and *Lotus japonicus* (Zagari *et al*., 2017). By contrast, although *Sco3* could be partially complemented by phytochrome B overexpression the function of the protein remains unknown and intriguingly some mutant alleles also affect the color of the adult leaves (Ganguly *et al*., 2015). The lack of functional annotation of the protein aside, it is clearly important in normal chloroplast and embryonic development as well as in flowering time and rosette formation (Ganguly *et al*., 2015). By contrast to the unknown function of SCO3, SCO2 is a protein disulfide isomerase and has been shown to play a role in chloroplast biogenesis (Tanz *et al*., 2012). More specifically it is involved in thylakoid development and interacts with other photosystem complex proteins (Zagari *et al*., 2017). The protein has a conserved zinc finger domain, containing four cysteines that have been shown to be relevant for the catalytic activity of the isomerase activity (Muranaka *et al*., 2012).

As mentioned above in *A. thaliana* mutants of the gene primarily show a cotyledon phenotype but under short-day conditions, true leaves also appear slightly paler (Zagari *et al*., 2017). Mutants in *Lotus japonicus* show a variegation phenotype, that resembles the one in tomato (Zagari *et al*., 2017). This interesting phenotype led us to characterize the *canal-1* tomato mutant at the molecular level. In the current study, we employ a multi-omics approach encompassing transcriptomics, proteomics, and metabolomics as well as measuring a range of photosynthetic parameters to shed light on the mechanism of yield canalization.

## Results

### Genetic, phenotypic and physiological characterization of the canal-1 mutant

The *canal-1* mutant has a missense mutation from T to G, 923bp downstream of the start codon of Solyc01g108200, leading to a conversion of Tryptophan (W) 147 to Glycine (G) (Figure 1 A + B)(Fisher and Zamir, 2021). We obtained a 3-D model of the protein in tomato from alphafold and visualized the spatial orientation of different key residues. The altered amino acid residue is in close proximity to four conserved cysteines (C151, C154, C174, C177), which are putatively responsible for the catalytic activity (Figure 1 C) (Shimada *et al*., 2007, Muranaka *et al*., 2012). The variegation phenotype of *canal-1* mutant leaves ranges from almost completely green over differently patterned variegated to completely pale white-beige (Figure 1 D-F). Additionally, the distribution of green and white leaf area varies from plant to plant (Figure 1 G-J). We estimated the content of different pigments in wild-type green leaves as well as in mutant leaves, which were predominantly green or predominantly white, through spectroscopic measurements (Figure 1 K). As expected, white *canal-1* leaves were almost completely devoid of both chlorophyll a and b and also total carotenoids. Interestingly, there were no statistically significant differences in any of these pigments, when comparing green mutant leaves to wild-type leaves. Stems from *canal-1* plants appeared as intermediate, having a strongly reduced albeit detectable level of chlorophyll (Supplemental Figure S 1). We also measured the fruit yield at maturity of plants grown in the greenhouse under both control and drought conditions (Figure 1 L). Unsurprisingly *canal-1* plants produced lower fruit yield than wild-type plants under control and drought conditions. We also wanted to estimate the effect of the mutation on yield stability as seen in the field. However, given that our experiments did not have the same layout as the so-called canalization replication (CANAREPs) trials performed in the field (Fisher and Zamir, 2021), instead of using the CV, we estimated variation by the scaled absolute deviation from the median (Figure 1 M). Indeed, *canal-1* mutants showed a much higher relative variation of fruit yield when compared to wild-type plants. This effect was however only significant under control conditions, potentially being limited by the small number of plants we subjected to drought stress.

**Figure 1:**
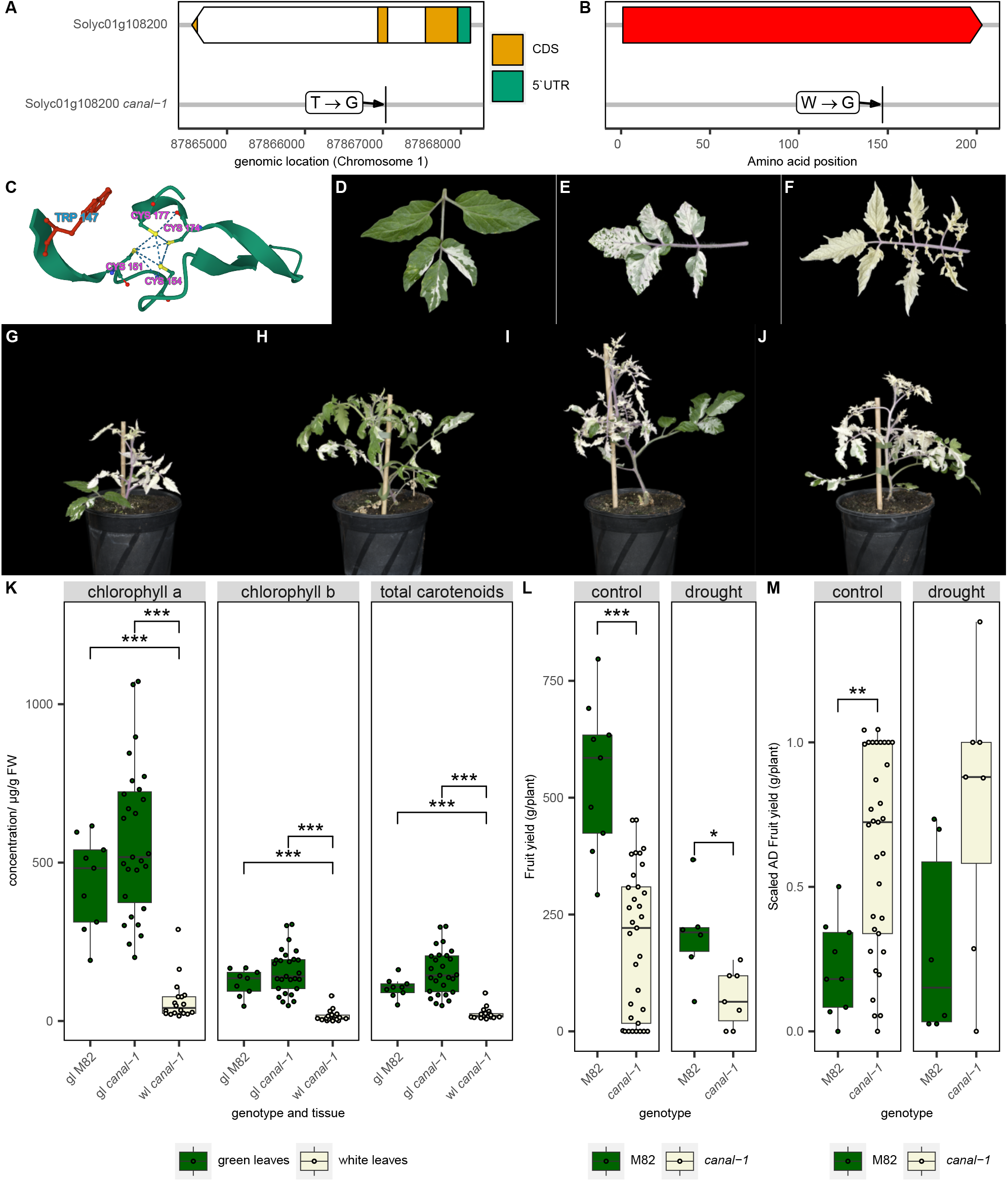
Genotype and Phenotype of canal-1 mutant. A, Schematic representation of gene model of Solyc01g108200, with coding sequence (CDS; orange) and 5’ untranslated region (5’UTR, green). Arrowhead of gene model box indicates gene direction. Label shows nucleotide changes in one letter code and arrow points to location of mutation. B, Schematic representation of protein sequence (top red box). Label shows amino acid changes in one letter code and arrow points to location of mutation. C, 3D-model of conserved zinc finger domain of wild type CANAL protein. Conserved cysteine residues (pink text and yellow side chain sulfur atoms) as well as the tryptophan (blue text and red side chain), that is altered in the *canal-1* mutant are highlighted. D-E, Phenotype of leaves from *canal-1* mutant with varying degrees of variegation. G-J, Habitus of different *canal-1* plants. K, Pigments measured in green and white leaves of mutant and wild type plants. Subpanels show different pigments. L, Fruit yield of M82 and *canal-1* plants. Subpanels show control and drought conditions. M, Scaled absolute deviation (AD) from the median fruit yield calculated from H. Subpanels show control and drought conditions. In G-J, images have been edited to remove parts of the bamboo stick beyond the size of the plant. All boxplots show the interquartile range (IQR) between the first and the third quartile as the box, with the median indicated by a black center line. Whiskers extend from quartiles to most extreme points with a maximum of 1.5 x IQR. Points beyond that range are considered outliers but overlaid by individual data points, shown as quasirandom. Boxes and points are filled according to tissue (green leaves: green, white leaves: beige) in K and according to genotype (M82: green; *canal-1*: beige) in L-M. Statistical significance was estimated by pairwise-wilcox test for K and kruskal-wallis test for L-M and significant differences are indicated by a bracket and asterisks (*: p ≤ 0.05; **: p ≤ 0.005; ***: p ≤ 0.0005).

### Transcriptomics reveal upregulation of photosystem assembly factors in green *canal-1* leaves

Due to the pigmentation phenotype, we considered that photosynthetic tissues would be the most interesting to characterize more thoroughly. We, therefore, selected three samples from wild-type leaves and stems as well as green and white leaves and stems from *canal-1* plants for RNA sequencing (see Materials and Methods). We mapped transcript reads with LSTrAP and analyzed differentially expressed genes (DEGs) via deseq2. The PCA already showed a clear separation of tissues and also highlighted white mutant leaves as being more distinct from green leaves of both mutant and wild type plants (Supplemental Figure S 2). For the analysis, we compared mutant white leaves to both mutant green leaves and wild-type green leaves, as well as green leaves and stems between wild-type and mutant (see Venn diagrams in Supplemental Figure S 3 A + B). Perhaps as expected, we saw the most DEGs when comparing white *canal-1* leaves to green M82 leaves (4873 DEGs), followed by white versus green leaves of the mutant (2390 DEGs, Figure 2 A). With 1404 DEGs green leaves of *canal-1* compared to M82 green leaves still showed some substantial differences, albeit to a minor degree, which is also characterized by smaller log2-fold change and –log10(p)-values in the volcano plot (Figure 2 A). The smallest differences were however detected between stems of mutant and wild type (Supplemental Figure S 4).

**Figure 2:**
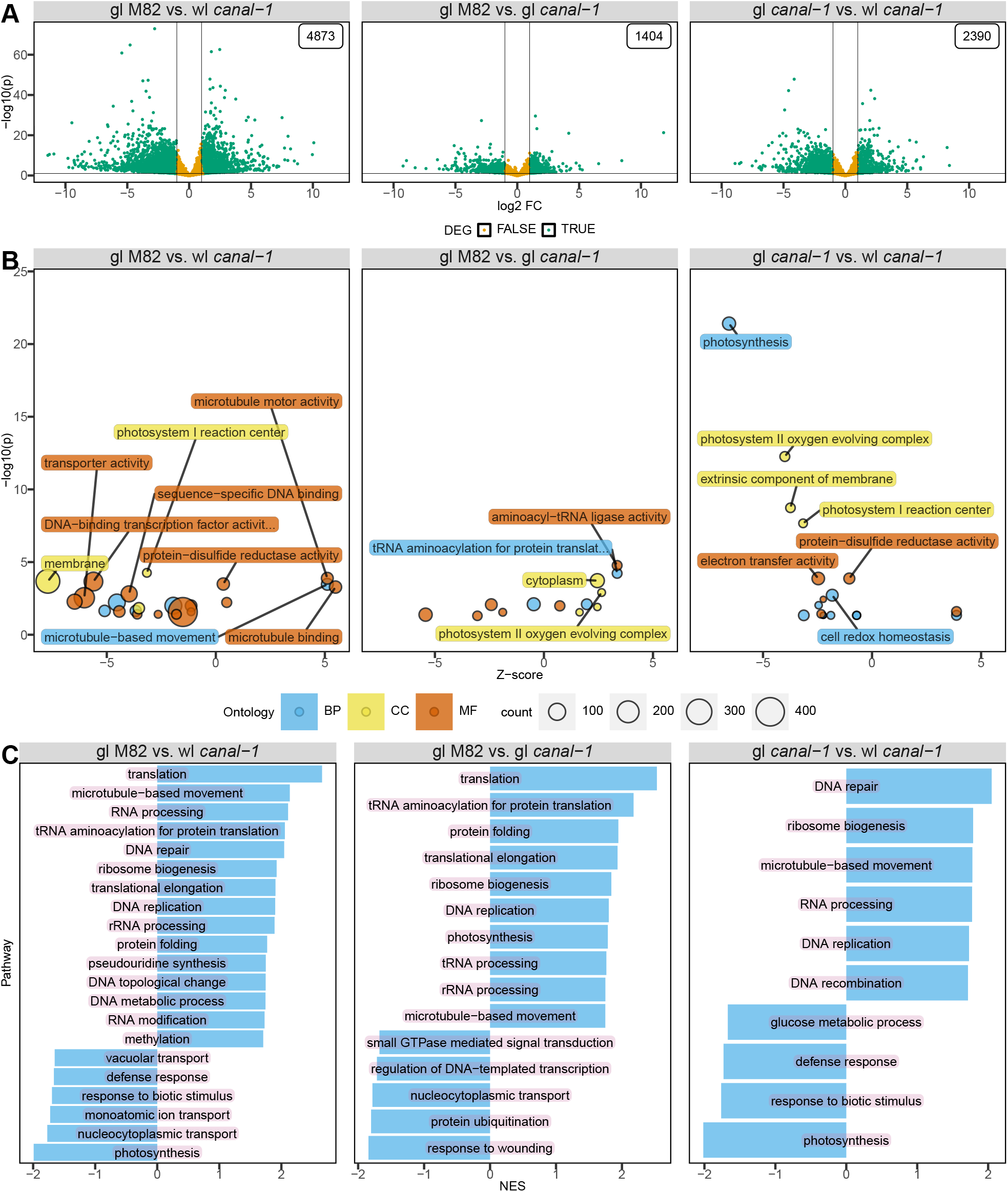
Transcriptome of canal-1 leaves compared to M82 leaves. A, Volcano plot of differentially expressed genes in different contrasts. Differentially expressed genes are shown in green and non-DE genes are shown in yellow. Label in top right-hand corner shows the number of DE genes of the respective contrast. B, Bubble plot with enriched gene ontologies according to topGO. Bubbles and labels are colored according to ontology and sized based on the number of genes in the respective term. C, Barplot of enriched gene sets according to fgsea. gl: green leaves; wl: white leaves; BP: biological process, cc: cellular component, MF: molecular function; NES: normalized enrichment score. (DEG: p ≤ 0.05 & |log2FC| ≥ 2)

We next investigated which kinds of genes were changed in the different comparisons by conducting gene ontology (GO) enrichment analysis (Figure 2B). Since GO enrichment does not consider up-or downregulation of genes within an enriched term, we calculated a z-score by subtracting the number of downregulated genes from the number of upregulated genes and divided this by the square root of the total count of genes in each term (see Formula 1). Accordingly, a positive z-score indicates more upregulated, than downregulated genes and a negative z-score indicates more downregulated than upregulated genes. When comparing white *canal-1* leaves to M82 leaves we found terms such as “photosystem I reaction center” and “protein disulfide oxidoreductase activity” enriched (Figure 2B left and Supplemental Table S 1). Although photosynthesis as a GO term for a biological process was also enriched here the enrichment was much clearer when comparing white leaves of *canal-1* to green leaves (Figure 2B right and Supplemental Table S 1 and Supplemental Table S 2). The comparison of green mutant and wild-type leaves yielded similar terms as the other comparisons (Supplemental Table S 3). Interestingly, the term “photosystem II oxygen evolving complex” is slightly upregulated in green leaves of *canal-1* plants compared to M82 leaves (Figure 2 B, Supplemental Table S 1 and Supplemental Table S 3). Furthermore, terms related to aminoacylation were also enriched (Figure 2B middle and Supplemental Table S 3). Enriched terms in the comparison of stem samples were rather related to cell wall processes (Supplemental Table S 4).

Similar to GO enrichment we also utilized gene set enrichment analysis (GSEA), which not only considers DEGs but takes all genes into consideration and ranks them according to their fold-change and *p-value* (Figure 2 C). GSEA highlighted translation as the most strongly enriched term in both comparisons of white and green leaves of mutant plants compared to wild type plants, but not when comparing the two different leaf types of the mutant (Supplemental Table S 6 - Supplemental Table S 8). Photosynthesis was the strongest downregulated term in the comparisons of white mutant leaves either to wild type leaves or to green mutant leaves (Figure 2 C). In agreement with the GO analysis, photosynthesis is upregulated in the green leaves of mutant plants, when compared to wild type plants. Irrespective of their color mutant leaves showed an upregulation of different DNA-, RNA, and protein-related terms, when compared to M82 leaves. The stem comparison only revealed a small number of enriched terms, related to lignin or protein (Supplemental Figure S 5, Supplemental Table S 9).

The discovery of the enriched protein disulfide oxidoreductase GO term (GO:0015035) and the fact, that SCO2 is a protein disulfide isomerase prompted us to investigate genes related to protein disulfide activity. Among genes belonging to the GO:0015035 one group of genes stood out (Figure 3). All of the genes showed, relatively consistently, low or no expression in wild type green leaves, an increased level in *canal-1* green leaves, and even higher expression in white mutant leaves. We noticed that these genes carry consecutive gene IDs and when checking the results from our orthology search, we realized, that they all belong to the same orthogroup, also including the CC-type glutaredoxin (ROXYs), *ROXY16* and *ROXY17* (Berardini *et al*., 2015, Jung *et al*., 2018)(Supplemental Figure S 6). As class III glutaredoxins, ROXYs comprise one of three classes of glutaredoxins, which are historically characterized by their glutathione-dependent reductase activity (Gutsche *et al*., 2015). ROXYs have been shown to be involved in various processes from flower development to stress responses (Gutsche *et al*., 2015). Several ROXYs (including ROXY16 and ROXY17) have been shown to be differentially expressed under nitrate starvation conditions and are believed to be involved in ROS-mediated nitrate signaling in *Arabidopsis thaliana* (Jung *et al*., 2018). It has also been shown, that CC-type glutaredoxins can act as transcriptional repressors for target promoters regulated by TGA-transcription factors (Uhrig *et al*., 2017).We also investigated the terms photosystem I reaction center and photosystem II oxygen evolving complex (Supplemental Figure S 7 & Supplemental Figure S 8). Genes from the photosystem I reaction center term showed a regular pattern of differential expression between white mutant leaves to both mutant and wild-type green leaves, with a lower abundance in white mutant leaves (Supplemental Figure S 7). For the term photosystem II oxygen evolving center, expression was also mostly significantly lower in white mutant leaves than in green mutant leaves and/or green wild type leaves but for some genes, the expression was significantly higher in green *canal-1* leaves in comparison to green M82 leaves (Supplemental Figure S 8).

**Figure 3:**
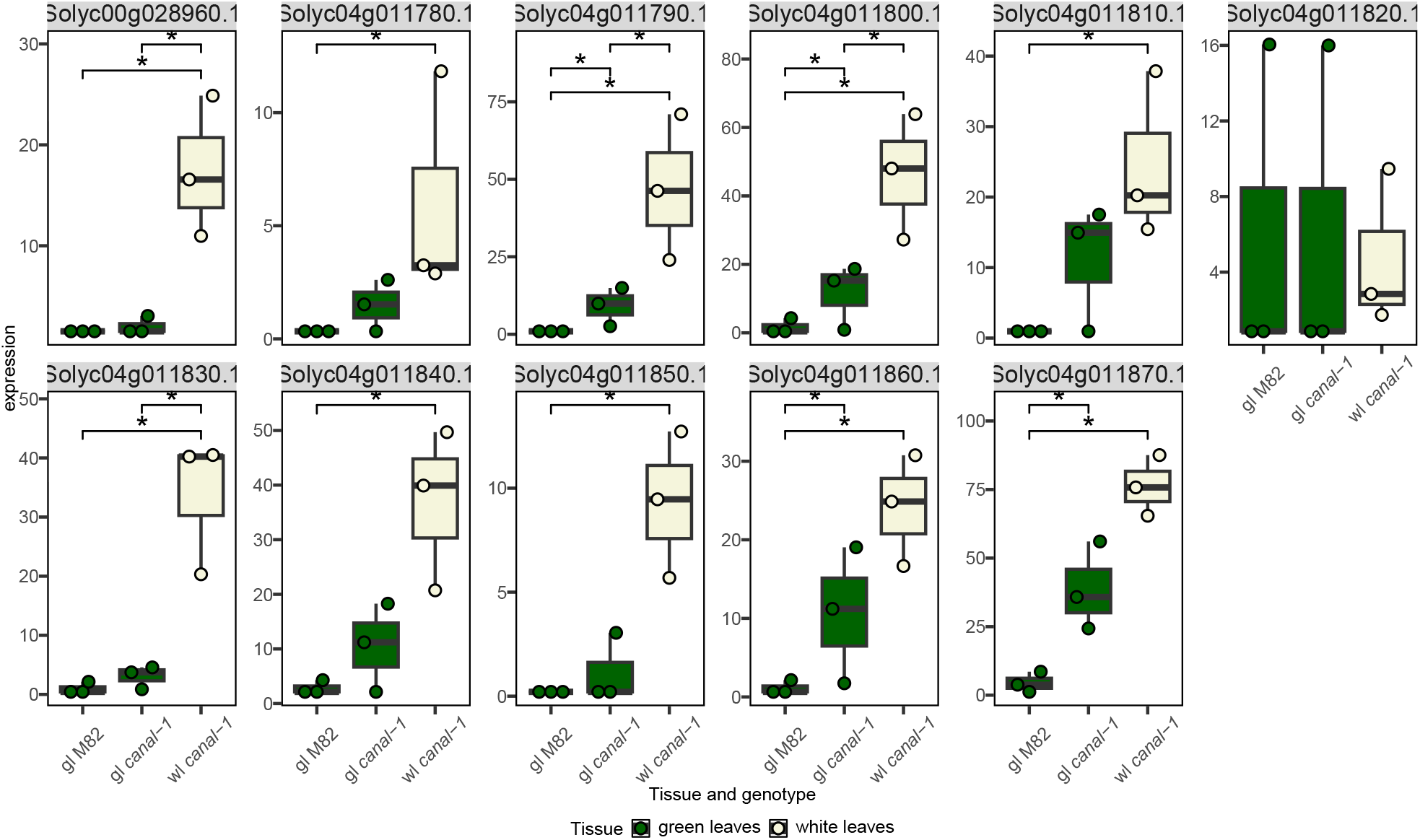
Expression of group of glutaredoxin genes. Boxes and points are filled according to tissue (green leaves: green, white leaves: beige). Statistical significance was extracted from differential gene expression analysis. Significant differences are indicated by a bracket and asterisks (*: p ≤ 0.05 & |log2FC| ≥ 2).

Due to the likely role of the SCO2 protein and the photosynthesis-related terms appearing in the enrichment analysis, we investigated the expression of different genes involved in photosystem assembly more deeply (Figure 4). First, as the protein under investigation is itself putatively involved in thylakoid assembly, we checked its expression in mutant and wild-type leaves (Figure 4 A). The gene however did not appear to be differentially expressed in any of the comparisons. To expand our scope to all potential photosystem assembly genes, we curated a list of known photosystem components and photosystem assembly genes in *A. thaliana* from the literature (Hankamer *et al*., 1997, Jensen *et al*., 2007, Shi *et al*., 2012, Berardini *et al*., 2015, Yang *et al*., 2015, Lu, 2016b) and matched them to the tomato genes via orthology searches (Supplemental Table S 5). When considering all these genes, we could see that green leaves of *canal-1* plants showed a significant upregulation of the expression in comparison to both white *canal-1* leaves and wild-type leaves (Figure 4 B). White *canal-1* leaves, however, showed a wild type level of expression of photosystem assembly genes, with the exception of a few outliers that displayed a strong upregulation. We visualized the level-scaled expression of these genes individually in a heatmap and could confirm that the vast majority shows the highest expression in green mutant leaves, with the exception of a small cluster of genes that showed the highest expression in white mutant leaves and another cluster of genes with the highest expression in wild type leaves (Figure 4 C). The assembly genes, which are highly expressed in white *canal-1* leaves are orthologs of ELIP1 (Solyc09g082690/700) and DEG7 (Solyc03g043660) as well as CPRabA5e (Solyc11g012460) and Ycf3 (Solyc00g500145), some of which are differentially expressed between white mutant leaves and/or green mutant and wild type leaves (Figure 4 C + D). Early light inducible proteins (ELIPs) have been shown to be involved in early chloroplast development (Casazza *et al*., 2005). We were interested to see how these differences affect the gene expression of the proteins which finally comprise the photosystems. We, therefore, filtered the transcriptome dataset for genes orthologous to known photosystem components in *A. thaliana* (Figure 4 E). In contrast to the assembly factors, the photosystem components are significantly reduced in the white leaves of the mutant compared to mutant and wild-type green leaves. Additionally, expression of these genes is also significantly increased in green *canal-1* leaves, relative to M82 leaves.

**Figure 4:**
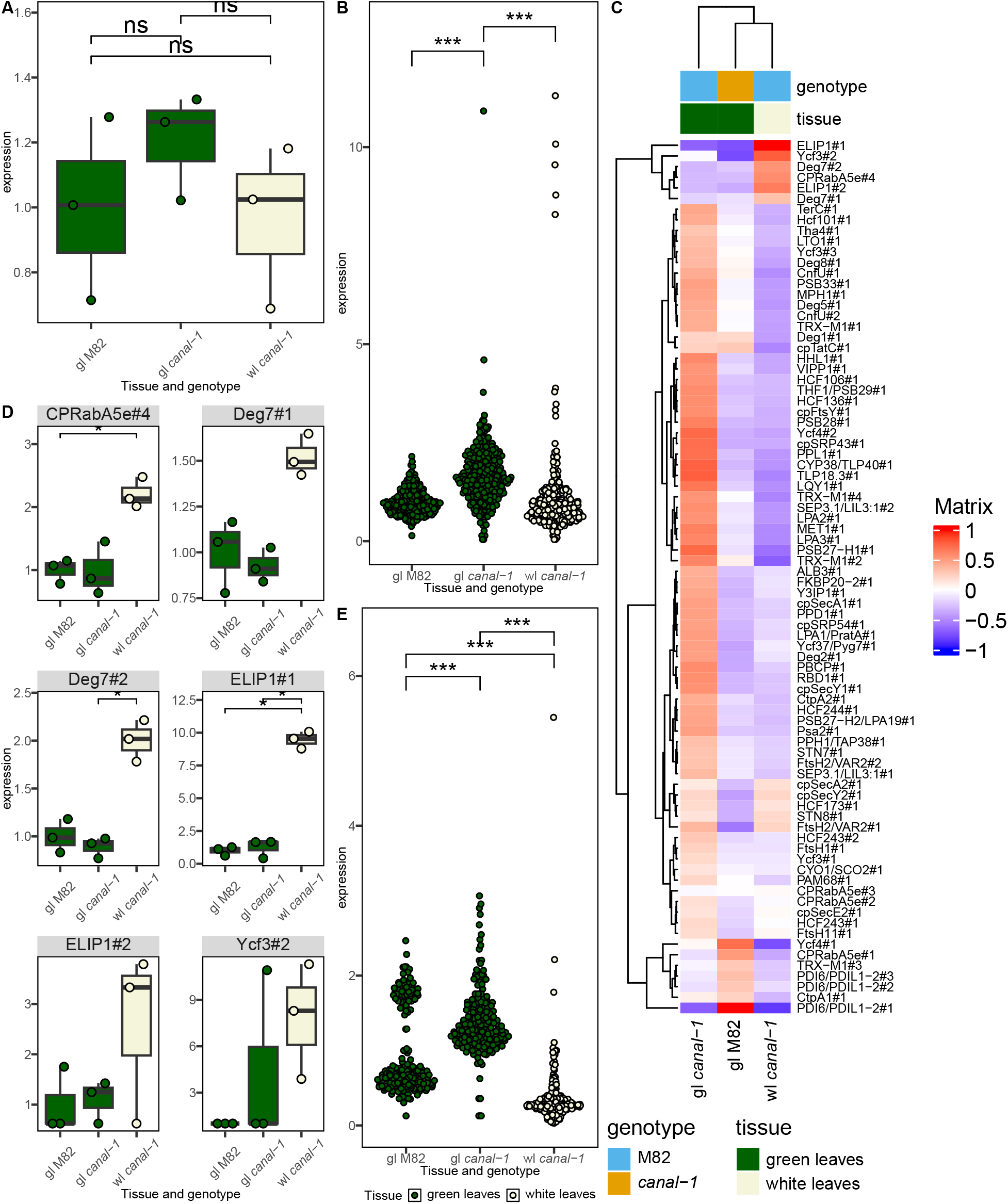
Transcriptome of different photosystem components in *canal-1* and M82 leaves. A, Expression of Solyc01g108200. B, Expression of photosystem assembly components. C, Heatmap of level-scaled expression levels for individual photosystem assembly components with color scale ranging from -1 to 1. D, Expression of orthologs to Deg7, ELIP1, CPRabA5e and Ycf3. E, Expression of protein complex genes of photosystems. Boxes and points are filled according to tissue (green leaves: green, white leaves: beige). To facilitate scaling to wild type samples, zero values were imputed by the half-minimum per gene. Statistical significance was estimated by pairwise-wilcox test for B and D and extracted from differential gene expression analysis for A + C. Significant differences are indicated by a bracket and asterisks for A + C (*: p ≤ 0.05 & |log2FC| ≥ 2) and for B and D (*: p ≤ 0.05; **: p ≤ 0.005; ***: p ≤ 0.0005).

### Proteomics show wild type like proteome of green *canal-1* leaves

Having assessed transcriptional changes we next investigated which of these translated to changes at the proteome level, for which we again selected samples from white and green *canal-1* leaves and M82 leaves, as well as stems from mutant and wild-type. When performing a PCA, we could once again see the separation of several groups of samples (Supplemental Figure S 9). Interestingly here, the white mutant leaves clustered even further away from green mutant and wild type leaves as compared to the PCA of RNAseq data (Supplemental Figure S 2).

Similarly, as for the transcriptome data, we determined differentially abundant proteins (DAPs) (Figure 5). For this purpose, we used an established shotgun proteomics platform and mapped the peptides to the tomato proteome UP000004994. We were able to identify a total of 6315 protein groups in this manner which could be further split, yielding 7729 different proteins. Interestingly, *canal-1* green leaves appeared to be wild-type like on a proteomic level (Figure 5). There was only a single protein A0A3Q7EIF5, which showed differential expression when comparing *canal-1* green leaves to M82 leaves (Figure 5 A). This protein was annotated as a component of the oxygen evolving center (OEC) of photosystem II (Figure 5B). This is surprising, considering that on a transcriptomic level the term “photosystem II oxygen evolving complex” showed enrichment in the comparison of green mutant and wild type leaves and expression was generally higher in green mutant leaves (Figure 2, Supplemental Figure S 8). When comparing the *canal-1* white leaves to *canal-1* and M82 green leaves, we found 1105 and 1443 DAPs, with 869 DAPs overlapping between them (Figure 5A). We used the GO terms to perform another enrichment analysis (Figure 5C). As would be expected, due to the single DAP, the comparison of green mutant and wild-type leaves did not yield any enriched terms. For the comparison of white mutant leaves and green wild type leaves the most significantly enriched term was photosynthesis and for the comparison of white and green mutant leaves it was chlorophyll binding (Figure 5C). Besides other terms related to photosynthesis, we found also terms related to RNA binding and ATP-dependent protein folding chaperone (Figure 5C, Supplemental Table S 10 and Supplemental Table S 11).

**Figure 5:**
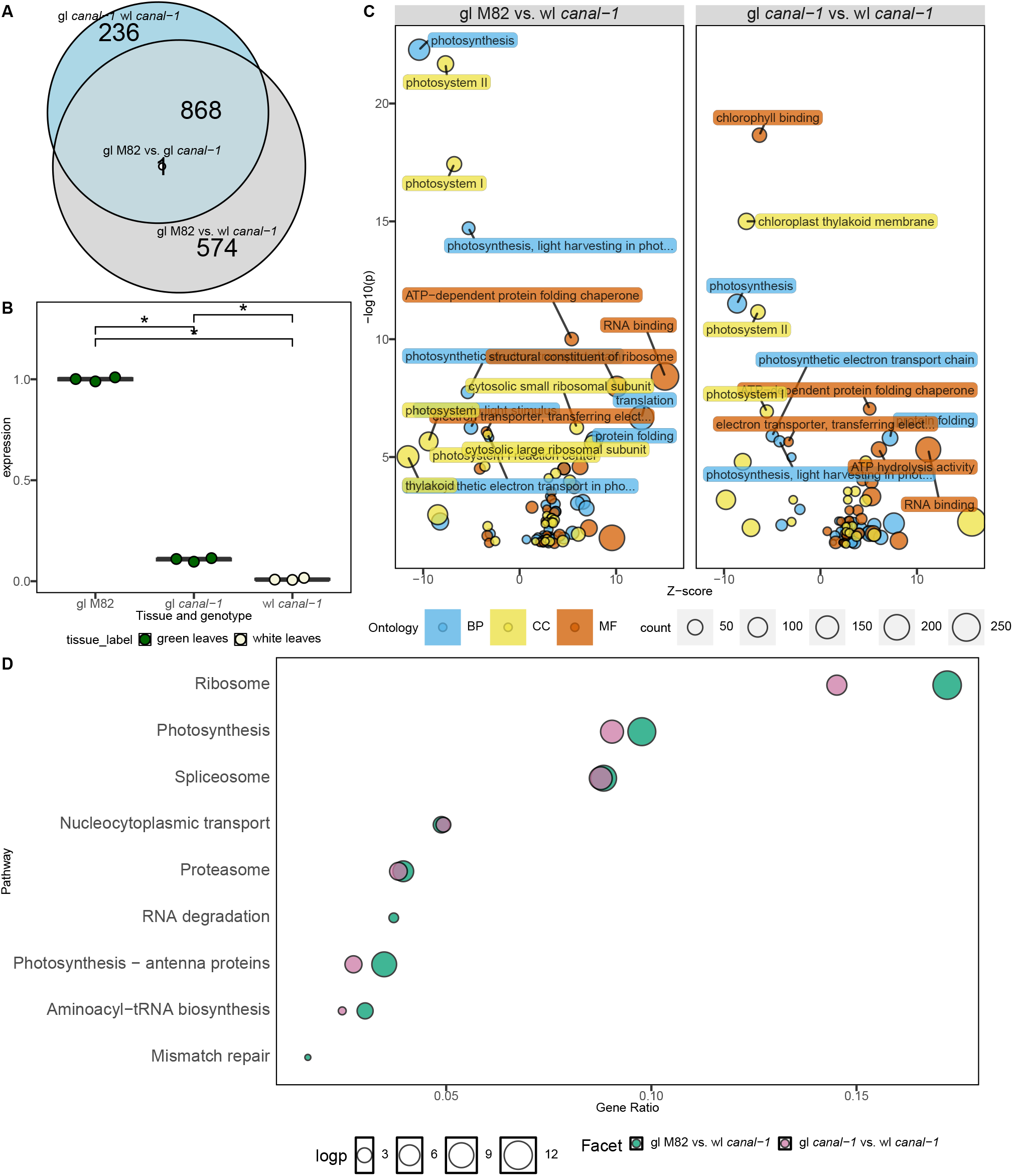
Differentially abundant proteins across different contrast. A, Euler plot of overlapping differentially abundant proteins between different contrasts. B, Relative abundance of oxygen evolving complex in mutant and wild type leaves. C, Bubble plots of enriched GO terms according to topGO. Bubbles and labels are colored according to ontology and sized based on the number of genes in the respective term. D, Enriched KEGG terms according to clusterProfiler. gl: green leaves; wl: white leaves

Having the opportunity to directly translate UniprotIDs into EntrezIDs, we also performed an enrichment analysis using KEGG terms (Figure 5D). Again we found photosynthesis terms enriched, but also the terms “Spliceosome”, “Ribosome” and “Proteasome”. (Supplemental Table S 12 + Supplemental Table S 13). In general, the pattern of enriched terms seems relatively congruent between both comparisons, highlighting the common differences between white *canal-1* leaves to both green *canal-1* and M82 leaves (Figure 5C and D).

Similar to the transcriptome data, we also took a closer look at protein levels of components and assembly factors of photosystems (Figure 6). Unfortunately we were not able to detect the protein of interest SCO2 itself among the 7729 protein and thus could not make any assumptions about its abundance. When considering other assembly factors, we could see a slightly different pattern to the transcriptome. The upregulation of assembly factors in green mutant leaves in comparison to green wild type leaves, that was visible on a transcriptomic level did not translate to the proteome level and there were no significant differences between the two groups (Figure 6A). Despite showing mostly lower values than green leaves, white mutant leaves did also not show any significant changes in the cumulative abundance of assembly factors (Figure 6A). We could again see however, a few outliers with a relatively high abundance in white leaves (Figure 6A). When looking at the level-scaled mean abundance of individual proteins, we could again see that orthologs of ELIP1 and DEG7 but also orthologs of FtsH11, HCF106, PDI6/PDIL1-2#2 built a cluster of proteins with higher abundance in white leaves of *canal-1* in comparison to green leaves of the mutant and wild type plants (Figure 6B). When looking at the individual proteins, we could see that while orthologs of ELIP1, DEG7 and FtsH11, showed some relatively high individual values only the orthologs of HCF106 and PDI6/PDIL1-2 displayed differential abundance between white leaves of the mutant and green leaves of wild type and mutant plants (Figure 6 C). We also investigated the abundance of the proteins that finally make up the photosystem components (Figure 6 D). Again the levels for white mutant leaves were extremely low and significantly lower than the protein levels in green leaves of mutant wild type plants. In contrast to the transcriptome however, green *canal-1* leaves showed a slightly but significantly reduced amount of photosystem components in comparison to green M82 wild type leaves (Figure 6C).

**Figure 6:**
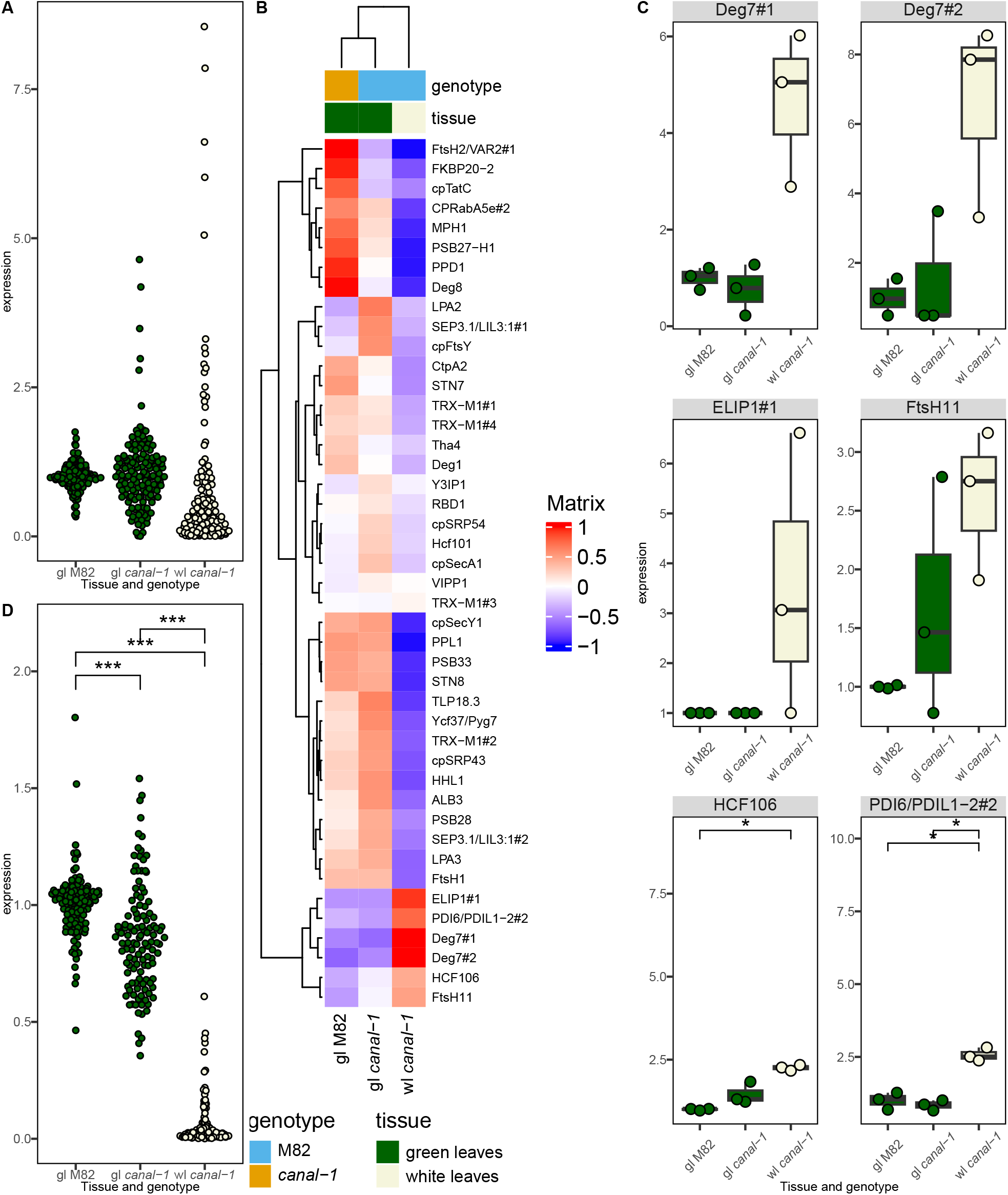
Protein levels of photosystem components and assembly factors in *canal-1* and M82 leaves. A, Assembly factors of photosystem components. B, Heatmap of level-scaled protein levels for individual photosystem assembly components with color scale ranging from -1 to 1. C, Individual proteins with high abundance in white mutant leaves. D, Abundance of photosystem components proteins in leaves. Statistical significance was estimated by pairwise-wilcox test for A and D and extracted from differential gene expression analysis for C. Significant differences are indicated by a bracket and asterisks for C (*: p ≤ 0.05 & |log2FC| ≥ 2) and for A and D (*: p ≤ 0.05; **: p ≤ 0.005; ***: p ≤ 0.0005).

### Integrating transcriptomic and proteomic data highlights stronger downregulation of photosystem components in white *canal-1* leaves

Having both transcriptomic and proteomic data available, we set out to visualize both regarding the photosystem components. We took the log10-fold changes of the contrasts comparing mutant green leaves to wild-type leaves as well as comparing mutant white leaves to wild-type leaves (Figure 7). On a transcript level, many of the photosystem components show a moderate upregulation in green leaves of *canal-1* leaves in comparison to wild-type leaves. White *canal-1* leaves, however, already show a downregulation, when compared to green M82 leaves. When considering the level of the proteome, we could see that the previously detected upregulation in green leaves of the mutant in comparison to those of the wild-type, translates to normal wild-type-like protein levels. White *canal-1* leaves conversely, show a reduction of protein abundance of photosystem components, in comparison to wild-type leaves.

**Figure 7:**
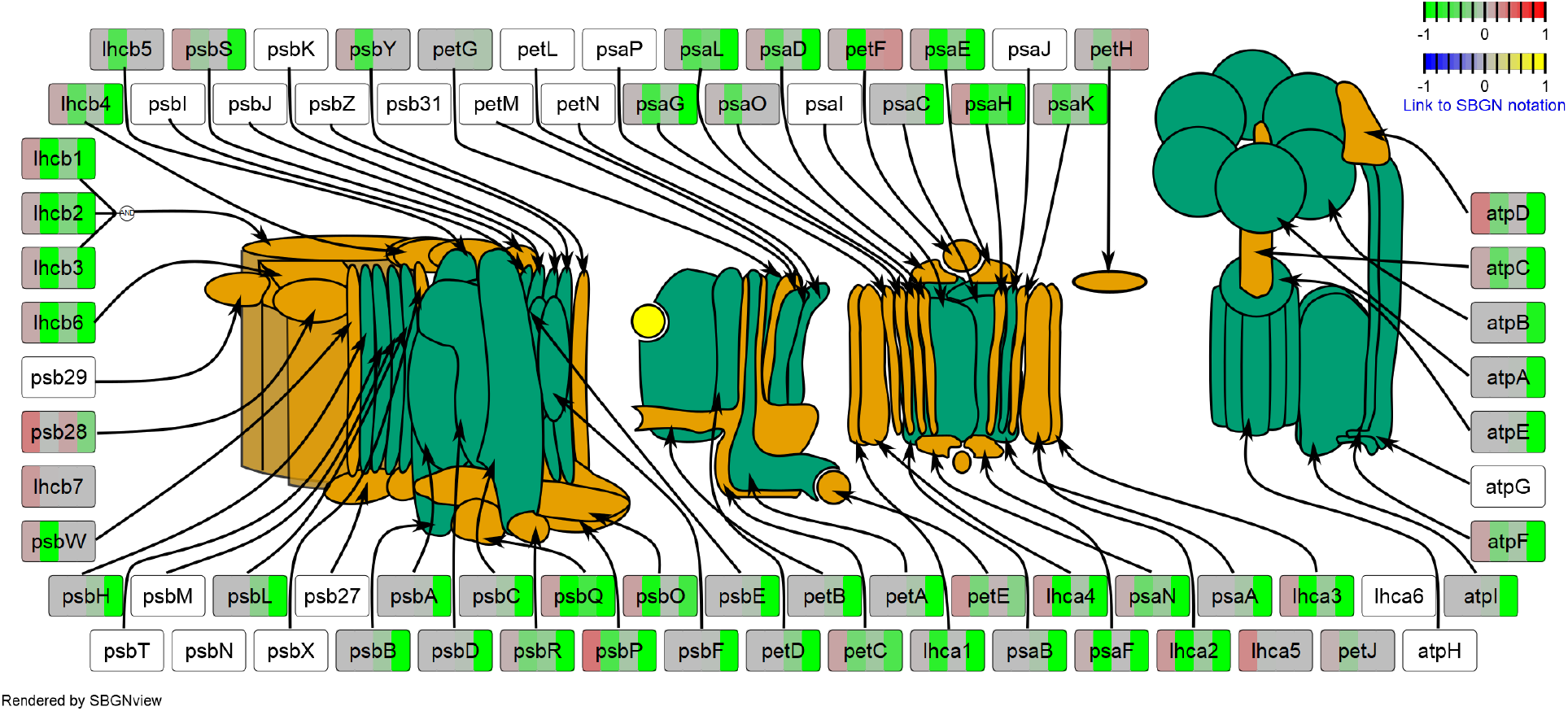
RNA and protein levels of photosystem components. Each rectangular glyph representing a photosystem component is split into 4 panels. The first two panels represent transcript levels, while the second two represent protein levels. In both the transcript and the protein section, the left panel shows the contrast of green *canal-1* leaves to wild type leaves, while right panel shows the contrast of white *canal-1* leaves to wild type leaves. Color mapping corresponds to log10-fold changes relative to wild type leaves as displayed by the color key in the top right corner. Image is drawn according to (Allen *et al*., 2011).

### Metabolomics reveal amino acid accumulation in white *canal-1* leaves

We next measured metabolites, extracted from different tissues of mutant and wild-type plants (Figure 8). For this purpose, we used established protocols for the GC-MS-based profiling of primary metabolites (Lisec et al., 2006) and the LC-MS measurement of polar secondary metabolites (Giavalisco *et al*., 2009) and lipophilic metabolites (Hummel *et al*., 2011), allowing the quantification of 126, 116 and 210 compounds, respectively (Supplemental Data S 1). We see the strongest changes in the white leaves of *canal-1* plants (Figure 8 A). At the very top of the heatmap we can see a cluster, mainly containing amino acids and derivatives thereof, that display a strong upregulation in white leaves of *canal-1* plants under control and also drought conditions, in comparison to the respective wild-type leaves. Compounds of another cluster below that, comprised mainly of triacylglycerides (TAGs) and galactolipids, show a moderate to strong downregulation in white *canal-1* leaves. Green leaves and stems of *canal-1* plants show similar patterns for these clusters, albeit to a much lesser extent (Figure 8 A).

**Figure 8:**
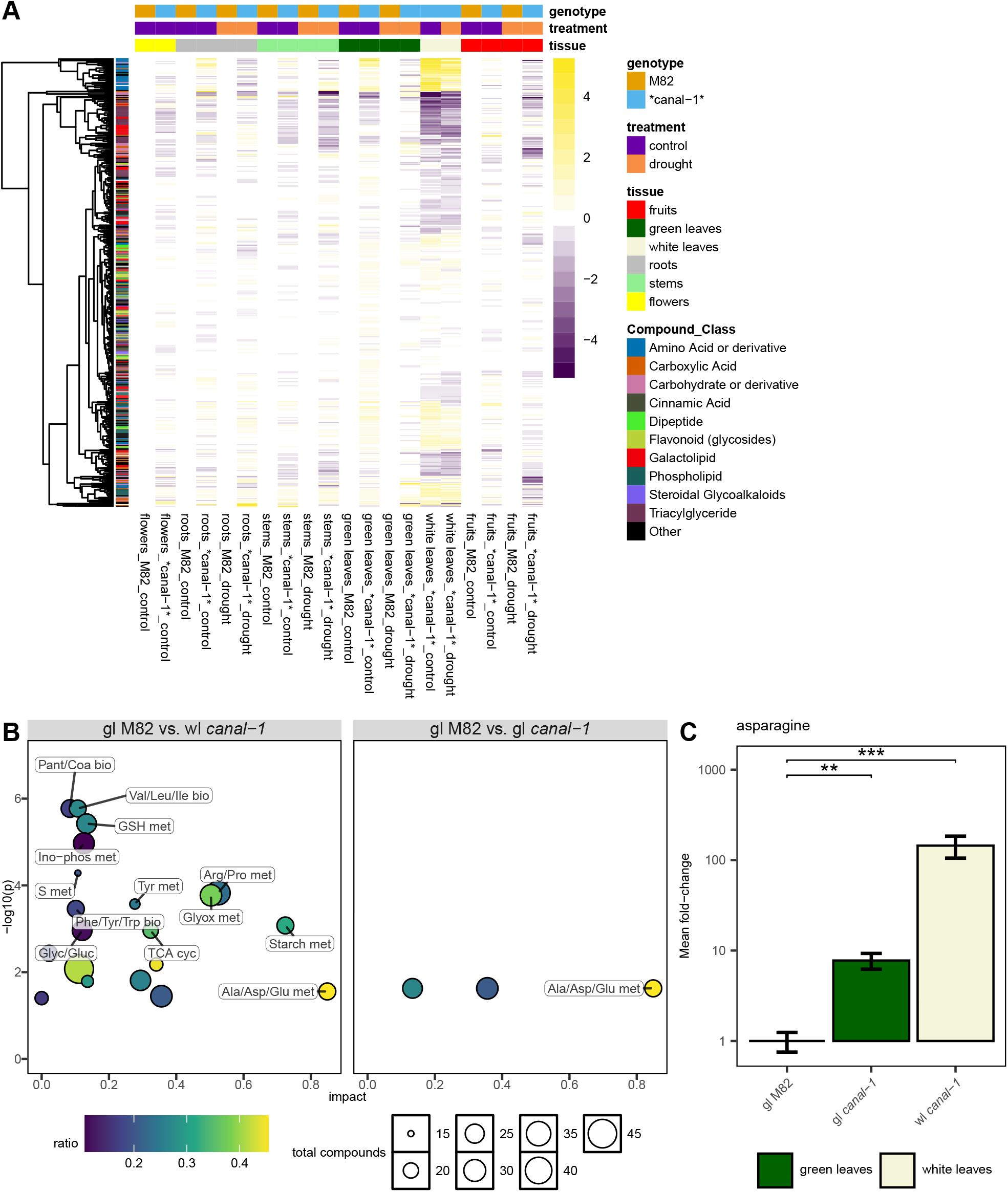
Metabolome of *canal-1* and wild type plants. A, Heatmap of all measured metabolites across different tissues genotypes and conditions. B, Pathway enrichment according to MetaboAnalyst. Bubbles are colored according to gene ratio and sized by their amount of total compounds in the respective pathway C, Asparagine level in *canal-1* and wild type leaves. Logarithmic scale was used for the y-axis. Statistical significance was estimated by pairwise-wilcox test for C. Significant differences are indicated by a bracket and asterisks for C (*: p ≤ 0.05; **: p ≤ 0.005; ***: p ≤ 0.0005). Full pathway names are shown in Supplemental Table S 12 and Supplemental Table S 13.

We wanted to gain a better idea of the altered primary pathways and used Metaboanalyst to perform a KEGG pathway enrichment (Figure 8 B). The pathway with the highest impact was “Alanine, Aspartate and Glutamate metabolism” and it could be detected in both comparisons (Supplemental Table S 14 and Supplemental Table S 15). We found many other pathways related to the biosynthesis and metabolism of various amino acids but also some related to carbohydrates (Figure 8 B, Supplemental Table S 14). The compound which showed the strongest changes was asparagine. In comparison to wild-type leaves it shows roughly a ten-fold increase in green mutant leaves and even a 100-fold increase in white mutant leaves (Figure 8 C).

### Multi-omics integration reveal shift of white mutant leaves from source to sink tissues

Having transcriptomic, proteomic and metabolomics data at our disposal, we took the opportunity to visualize them together and investigate more deeply the pathways, revealed by KEGG pathway enrichment. Starting at the reductive part of the pentose phosphate pathway (Calvin-Cycle), we could see that green *canal-1* plants showed some upregulation of genes on a transcriptomic level, which related to a minor upregulation on the proteomic level for example for phosphoribulokinase, chloroplastic (PRK) and RuBisCO (Figure 9). White *canal-1* leaves however showed a downregulation of transcripts and most proteins. Again this can be seen well in PRK and RuBisCO, which show downregulation on a transcriptomic and proteomic level. Several enzymes of the (oxidative) pentose phosphate pathway, some of which are shared with the Calvin-Cycle however showed an upregulation and additionally the compound 6-phospho-D-gluconate (6PG) was reduced in white *canal-1* leaves (Figure 9). The compounds sucrose, glucose, fructose and maltose are strongly reduced in white *canal-1* leaves, whereas in green mutant leaves sucrose shows wild-type levels and maltose even an increase in comparison to wild type leaves. Several enzymes like beta-glucosidase (BGLU), endoglucanase (NGLU), 6-phosphofructokinase (PFK) fructose-1-6-bisphosphate bisphosphate aldolase (FBPA) and fructose-1,6-bisphosphatase (FBP) show different or even opposite responses in white and green mutant leaves, suggesting that carbohydrates are catabolyzed in white leaves and synthesized in green leaves of the mutant (Figure 9). Also in the TCA cycle and the glyoxylate and dicarboxylate metabolism, several catabolic enzymes are upregulated on a proteomic level in white *canal-1* leaves and downregulated in green *canal-1* leaves. The amino acids valine, leucine, isoleucine, were upregulated in green leaves but even more strongly in white leaves of the mutant. We could also see that enzymes involved in this biosynthetic pathway, namely (ALS), (ILVC), (ILVD) and (ILVE) displayed an upregulation, which was stronger in white leaves than in green leaves of *canal-1* plants. Similarly, glutamate, glutamine, alanine, aspartate and asparagine were upregulated in mutant leaves. For glutamate and glutamine, we could see the upregulation of glutamate synthase (ferredoxin) (GLUS), glutamate dehydrogenase (NAD(P)+) (GLUD) and glutamate synthase (NADH) (GOGAT) which could explain the elevated amino acid levels. In the aspartate and asparagine biosynthesis we found asparagine synthase (glutamine-hydrolysing) (ASNS) and in white leaves also aspartate aminotransferase, cytoplasmic (AAT) upregulated, which likely lead to a higher flux from oxaloacetate to these two amino acids (Figure 9). These observations supported the results from the KEGG pathway enrichment.

**Figure 9:**
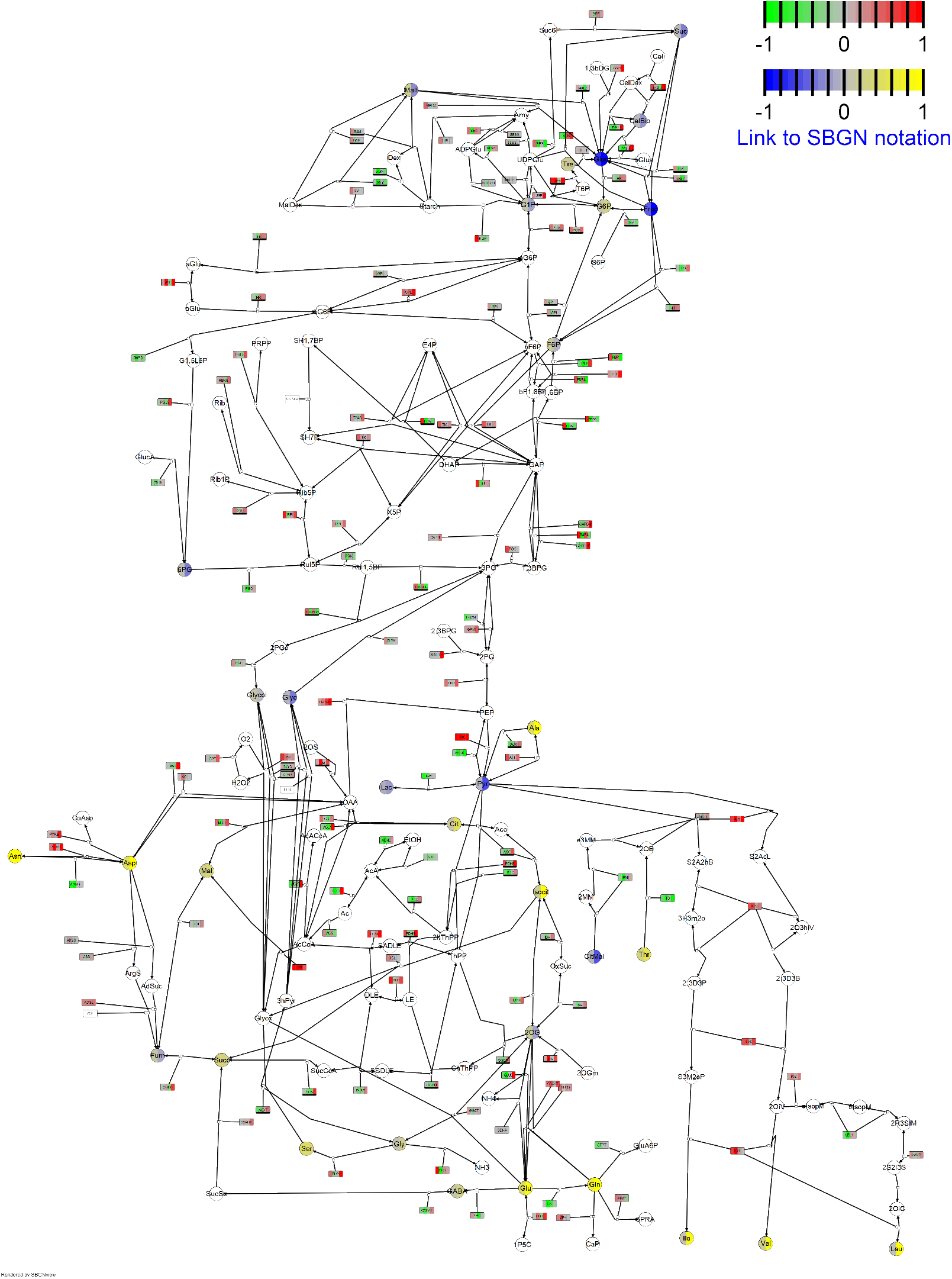
Pathway map derived from KEGG pathways designed in SBGN format showing relative transcript, protein and metabolite levels of green and white *canal-1* leaves in comparison to wild-type leaves. Rounded rectangular glyphs are split into four sections, where the first two depict transcript levels and the last two depict protein levels. For both transcript and protein levels, the first panel displays the relative level of green mutant leaves and the second panel the relative level of white mutant leaves in comparison to wild-type leaves. Circular glyphs depict metabolites, which are split in the same way as the transcript and protein panels. Colors correspond to log10-fold changes limited from -1 to 1 as depicted by legend in upper right corner (green to red for genes and proteins, blue to yellow for metabolites). Nodes without mapping data are uncolored. Entities that had to be duplicated to optimize the layout carry a black bar as a clone marker on the lower end of the glyph. Full names of genes/enzymes and compounds are found in Supplemental Data S 10.

### Correlation network construction supports a hub function of SCO2

The availability of this multi-omics dataset for the 15 samples from mutant and wild type stems and leaves also gave us the opportunity to explore how transcript-, protein- and metabolite levels correlate to one another. We therefore calculated the Spearman-correlation value for all pairwise-combinations, resulting in over 900 million correlation values. In order to reduce the amount of correlations to a manageable amount we directly filtered out correlation values below 0.75, which still left us with more than 29 million correlations. For an overview, we applied an even more stringent threshold of 0.95, created a correlation network graph using absolute correlation values as edge-weights and only retained the largest connected graph for a visualization (Figure 10). This resulted in a graph with 12478 vertices and 326539 edges. We could visually discern different groups of nodes. The whole network forms a ring like structure that is more strongly connected at the top and only sparsely on the base. Towards the right-top-hand side of the network graph we found a tightly connected set of transcript nodes together with also many protein nodes (Figure 10). Towards the right there was a patch of secondary metabolites connected to protein and transcript nodes and below that was a patch of lipid nodes where not that many protein nodes could be found anymore. Towards the left we could see a cluster of nodes which comprises mostly protein vertices, together with some transcript and lipid nodes. We were also interested to find out, where our gene of interest lies within the correlation network. As mentioned earlier, we were unable to detect the protein sequence in the proteomics dataset but we could see that the transcript of SCO2 was located very centrally within the large set of transcript and protein nodes (Figure 10). Due to its function in canalization of yield, we wondered, whether SCO2 shows typical properties of a network hub and therefore compared different network metrics of SCO2 to those of the overall network (Figure 11). When comparing the degree, closeness and betweenness of SCO2 to the rest of the filtered network, we realized that the values of SCO2 were indeed significantly higher than the (pseudo-)median of the overall network (Figure 11 A, Supplemental Table S 16). While the degree of 29 of SCO2 is only somewhat higher than the median degree of the network, the metrics closeness and betweenness both exceed the third quantile of the overall network (Figure 11 A). Focusing further on our gene of interest we queried the direct neighbors that SCO2 has at the 0.95 threshold (Figure 11 B). As already observed by the degree of SCO2, we could now see the 29 neighbors which strongly correlated to SCO2 and also among themselves (Figure 11 B, Table 1). The first observation we could make is that most of the SCO2-neighbors are transcript nodes, with only 3 protein nodes. Out of the correlations to SCO2 only 6 were negative, while 23 were positive. The strongest negative correlation was to the transcript Solyc06g053710.3, which is annotated as ethylene receptor homolog (ETR4) (Figure 11 B, Table 1). The strongest positive correlation was to the transcript Solyc09g010110.3 which is annotated as Chaperone protein dnaJ-related protein. Besides this gene we also found two more transcripts carrying an annotation relating to DnaJ proteins, namely Solyc12g009440.2 and Solyc01g105340.4. Solyc09g010110.3 is orthologous to the ORANGE gene in Arabidopsis (AT5G06130.2), which has been shown to be involved in carotenoid biosynthesis by binding to phytoene synthase (Zhou *et al*., 2015). Other interesting correlation partners were Solyc12g096100.2 and Solyc00g500066.1, which carry the annotations Protein PAM68, chloroplastic and Photosystem I assembly protein Ycf4, respectively. These are also orthologous to the Arabidopsis assembly factors PAM68 (AT4G19100) and Ycf4 (ATCG00520), respectively, which we found in our orthology search before (Supplemental Table S 5). This observation made us wonder, whether SCO2 transcript abundance also correlates to the abundance of other photosystem assembly factor transcripts or proteins, albeit with a lower correlation value and how strong the correlation is to the components that the photosystem finally consists of. We therefore reinvestigated the whole dataset with the more relaxed threshold of 0.75 and looked at correlations between photosystem components and assembly factor transcripts and proteins. This resulted in a dense network of transcript and protein nodes connected mostly through positive correlations (Figure 11 C). Both transcripts and protein nodes connected more strongly to members of their own class than to members of other classes. We found that SCO2 clustered together mostly with transcripts of other assembly factors like Ycf4 and PAM68 towards the left of the network. Towards the middle we found cluster with mostly transcript nodes of assembly factors and towards the right half of the network we found more protein nodes, with a densely connected cluster of protein nodes of photosystem components (Figure 11 C). We can make a similar observation, by using a hierarchical clustering algorithm of all raw unfiltered correlation values of the previously discussed transcripts and proteins and plotting them on a heatmap (Supplemental Figure S 10).

**Figure 10:**
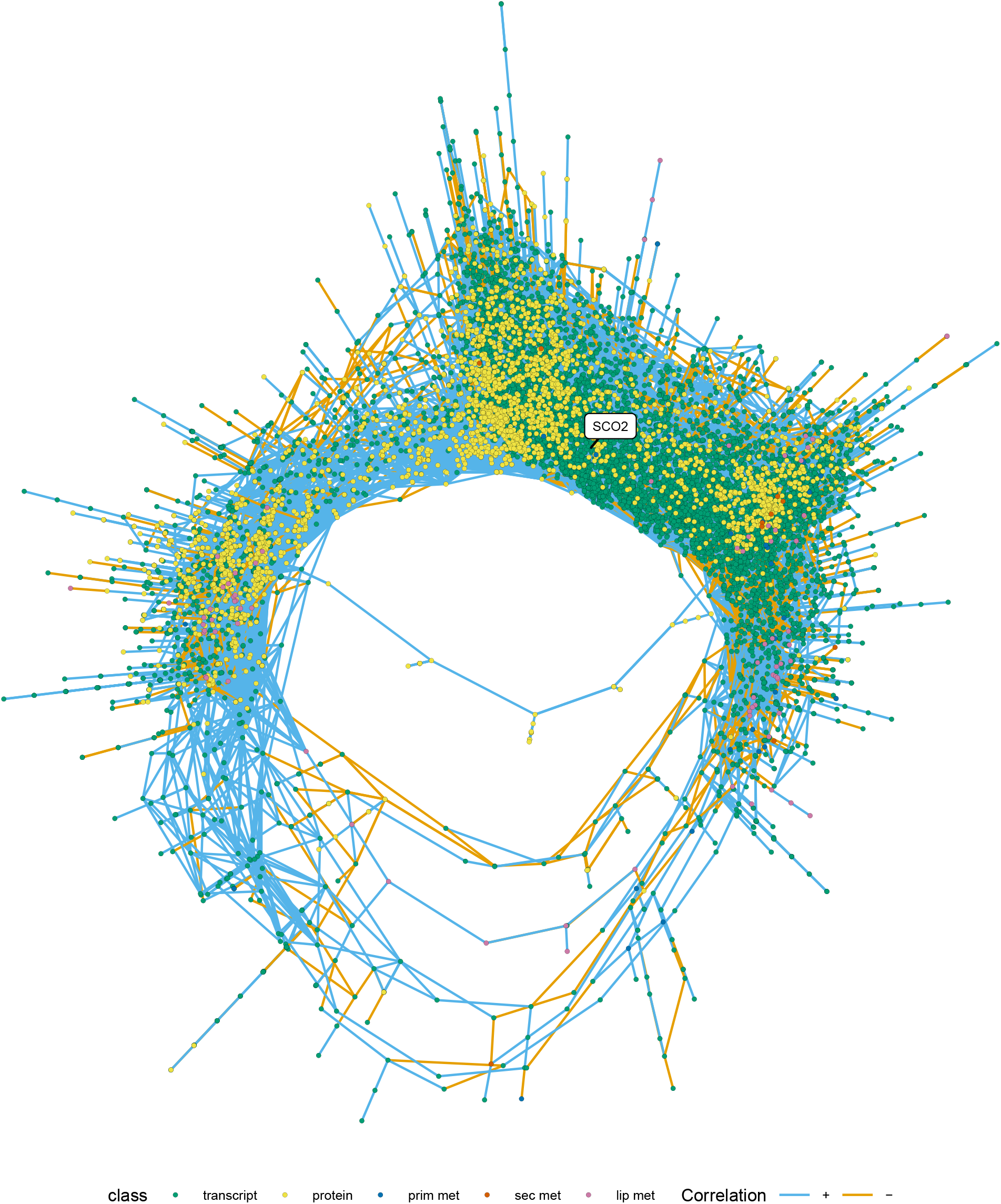
Correlation network graph. Spearman’s correlation value was calculated between levels of transcripts, proteins, primary metabolites, secondary metabolites and lipophilic compounds across leaf and stem wild-type and mutant samples and correlation values were used to construct a single connected graph. Vertices correspond to individual transcripts, proteins or compounds and are displayed by filled points, colored according to their vertex class. Vertices are connected through edges colored by the sign of the correlation. Graph layout was generated by a progressive multidimensional-scaling algorithm, using the inverse of the edge-weights. Correlation-value threshold = 0.95, n_vertex_ = 12478; n_edge_ = 326539.

**Figure 11:**
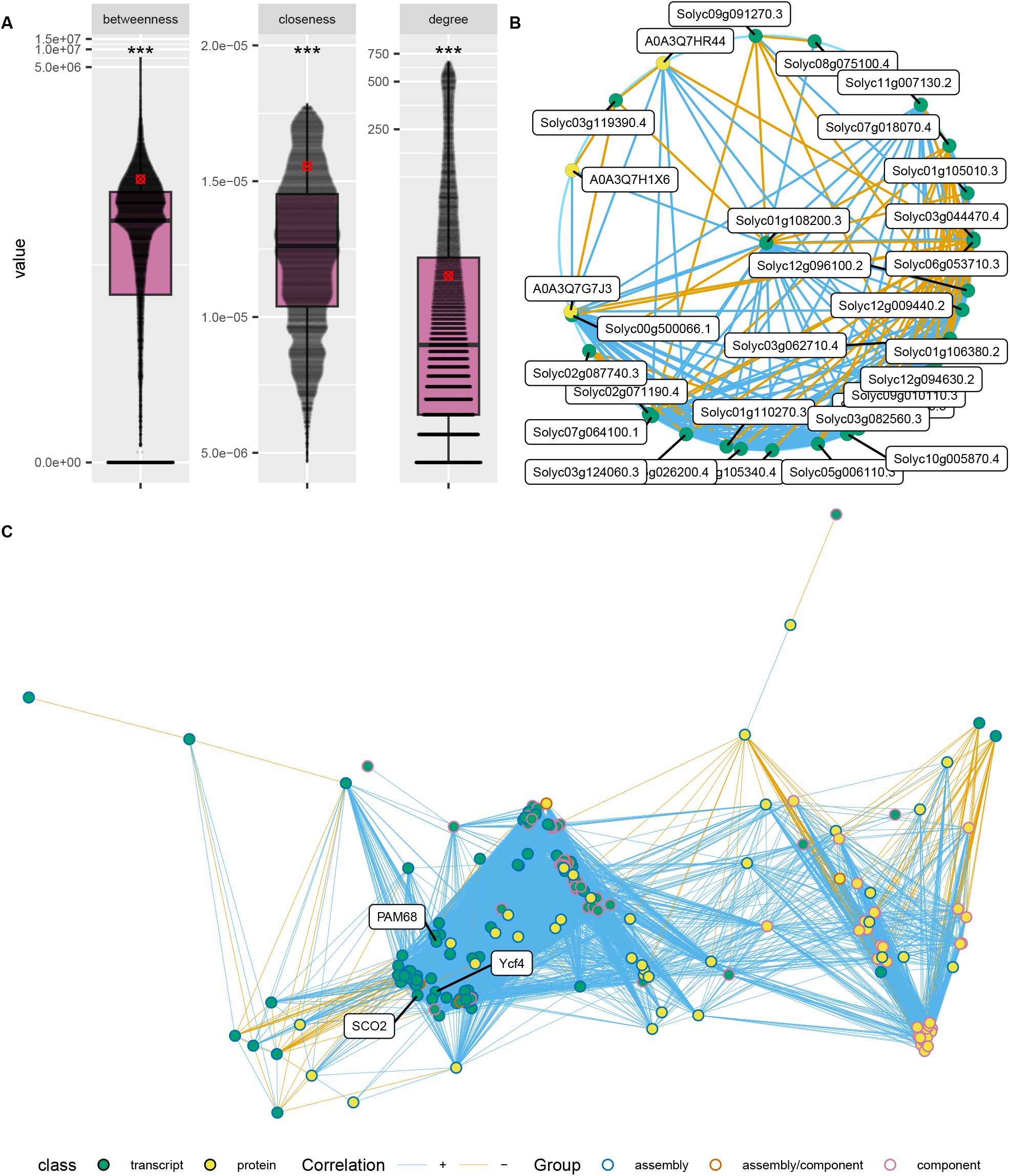
Subnetworks and network metrics. A, Betweenness, closeness and degree of SCO2 transcript in relation to the overall network. All boxplots show the interquartile range (IQR) between the first and the third quartile as the box, with the median indicated by a black center line. Whiskers extend from quartiles to most extreme points with a maximum of 1.5 x IQR. Points beyond that range are considered outliers but overlaid by individual data points, shown as quasirandom. The respective SCO2 value is indicated by a red cross-circle. Y-axis is transformed by log(1+x) to accommodate 0 values as well as large outliers. B. Correlation network graph of direct SCO2 neighbors at correlation value threshold 0.95. C, Correlation network graph of photosystem assembly factor genes and proteins at correlation value threshold of 0.75. In B + C, vertices correspond to individual transcripts or proteins and are displayed by filled points, colored according to their vertex class. Vertices are connected through edges colored by the sign of the correlation. In C the outline of the points (stroke) is colored according to the functional grouping. Graph layout was generated by a circular layout algorithm for B and a progressive multidimensional-scaling algorithm for C, using the inverse of the edge-weights. Statistical significance was estimated by single-wilcox test for A. Significant differences are indicated by asterisks for A (*: p ≤ 0.05; **: p ≤ 0.005; ***: p ≤ 0.0005).

**Table 1:** Direct neighbors of SCO2 transcript with absolute correlation-value above 0.95.

| Gene/Protein | ITAG4.0 annotation | Correlation value |
| --- | --- | --- |
| Solyc06g053710.3 | ethylene receptor homolog (ETR4) | -0.971428571 |
| Solyc09g091270.3 | cotton fiber protein | -0.964285714 |
| Solyc01g105010.3 | Non-specific lipid-transfer protein-like protein | -0.960714286 |
| Solyc02g087740.3 | Plant cysteine oxidase 2 | -0.957142857 |
| Solyc07g018070.4 | Double Clp-N motif-containing P-loop nucleoside triphosphate hydrolases superfamily protein | -0.957142857 |
| Solyc03g119390.4 | bHLH transcription factor 026 | -0.95 |
| Solyc01g106380.2 | hypothetical protein | 0.95 |
| Solyc01g110270.3 | RNA-binding CRS1 / YhbY (CRM) domain protein | 0.95 |
| Solyc03g082560.3 | Aldo-keto reductase/ oxidoreductase | 0.95 |
| Solyc12g094630.2 | Low-density receptor-like protein | 0.95 |
| Solyc01g105340.4 | Chaperone protein DnaJ | 0.953571429 |
| Solyc02g071190.4 | Carboxyl-terminal-processing protease-like protein | 0.953571429 |
| Solyc03g026200.4 | protein COFACTOR ASSEMBLY OF COMPLEX C SUBUNIT B CCB2, chloroplastic | 0.953571429 |
| Solyc03g044470.4 | red chlorophyll catabolite reductase | 0.953571429 |
| Solyc03g062710.4 | protein NDH-DEPENDENT CYCLIC ELECTRON FLOW 5 | 0.953571429 |
| Solyc12g096100.2 | Protein PAM68, chloroplastic | 0.953571429 |
| A0A3Q7HR44 | NA | 0.953571429 |
| Solyc00g500066.1 | Photosystem I assembly protein Ycf4 | 0.957142857 |
| Solyc03g063240.3 | Pyridoxamine 5'-phosphate oxidase-related FMN-binding protein | 0.957142857 |
| Solyc03g124060.3 | Polyketide cyclase/dehydrase and lipid transport superfamily protein | 0.957142857 |
| Solyc08g075100.4 | Initiation factor 4F subunit (DUF1350) | 0.957142857 |
| Solyc11g007130.2 | Major facilitator superfamily | 0.957142857 |
| A0A3Q7H1X6 | NA | 0.967432647 |
| Solyc07g064100.1 | Chlororespiratory reduction 41 | 0.967857143 |
| Solyc12g009440.2 | DnaJ/Hsp40 cysteine-rich domain superfamily protein | 0.967857143 |
| A0A3Q7G7J3 | NA | 0.967857143 |

### Photosynthesis in green leaves of *canal-1* is increased to compensate impaired photosynthesis in white leaves

We were additionally interested to see, whether the changes observed via the –omics analysis were also reflected in the photosynthetic activity and therefore estimated several photosynthetic parameters using a LiCOR (Figure 12). The assimilation rate of green leaves of the *canal-1* plant was significantly increased in comparison to wild-type leaves, whereas white mutant leaves showed a negative assimilation rate, which means, that they were actually carrying out net respiration (Figure 12 A). Additionally, green leaves of the *canal-1* mutant had a significantly increased stomatal conductance and transpiration rate compared to wild-type leaves, whereas white mutant leaves showed wild-type levels (Figure 12 B and C). Regarding the quantum yield of photosystem II, green leaves of mutant and wild-type showed similar levels, whereas white *canal-1* showed a statistically significant lower level (Figure 12 D).

**Figure 12:**
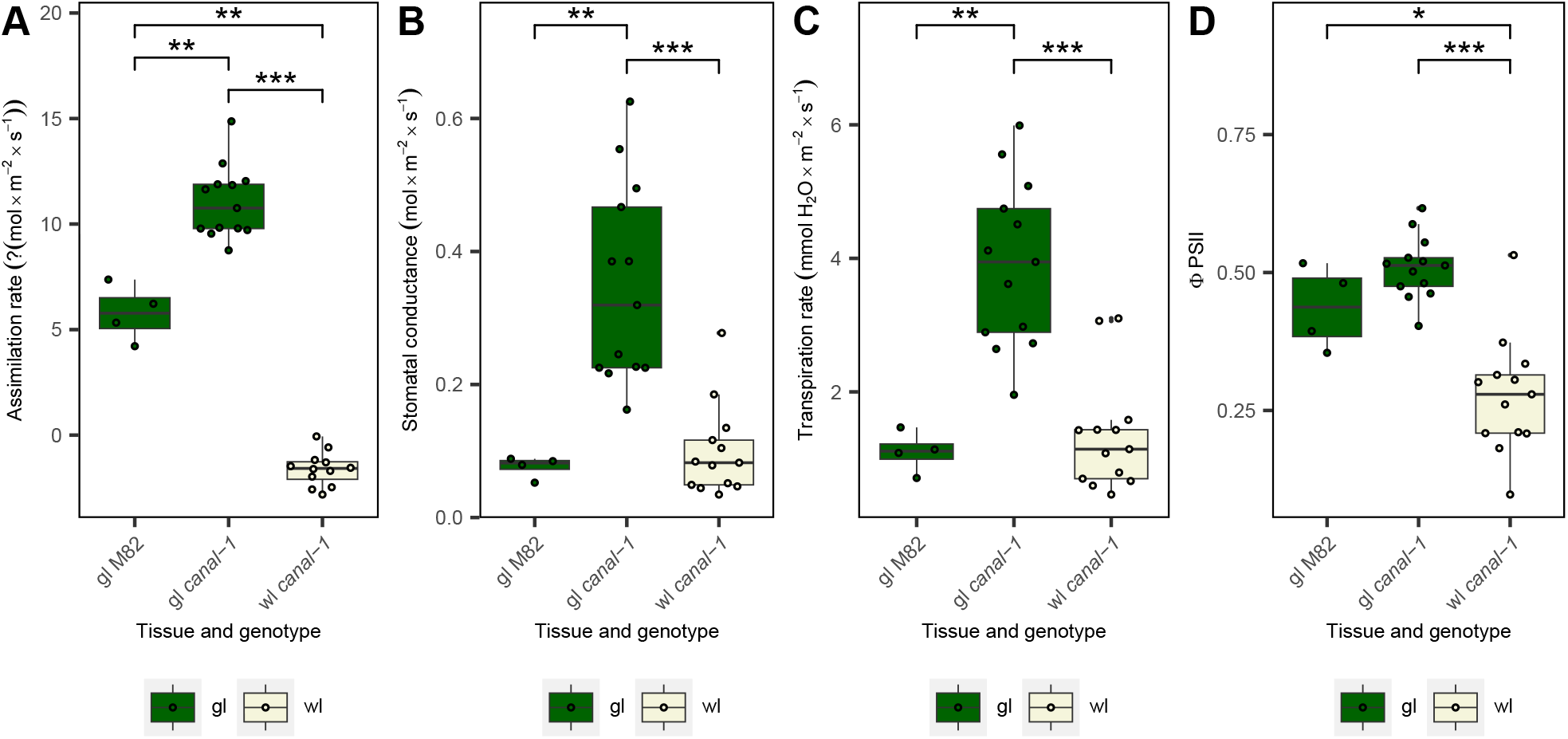
Photosynthetic measurements as conducted by LiCOR in the light. A, Photosynthetic assimilation rate. B, Stomatal conductance. C, Transpiration rate. D, Quantum yield of photosystem II (DPSII). Boxes and points are filled according to tissue (gl = green leaves: green, wl = white leaves: beige). Statistical significance was estimated by pairwise-wilcox test for A-D. Significant differences are indicated by a bracket and asterisks (*: p ≤ 0.05; **: p ≤ 0.005; ***: p ≤ 0.0005).

In a final experiment, we grew the *canal-1* mutant, the allelic *ghost-2* mutant, and three lines over-expressing the protein of interest, together with their respective wild-type cultivars, M82, Sioux (SX) and MoneyMaker (MM) in a greenhouse experiment (Figure 13). As observed before, the *canal-1* mutant showed a significantly decreased yield, as did the *ghost-2* mutant (Figure 13 A). In fact, the effect of the *ghost-2* mutant, was so strong, that neither of the plants was able to produce any fruit (Figure 13 A). Lines overexpressing the protein of interest, did not show significant differences regarding fruit weight in relation to the MoneyMaker wild-type (Figure 13 A). However, Line CANAL-OX31 shows a significantly earlier flowering time and an increased number of fruits per plant (Supplemental Figure S 11).

**Figure 13:**
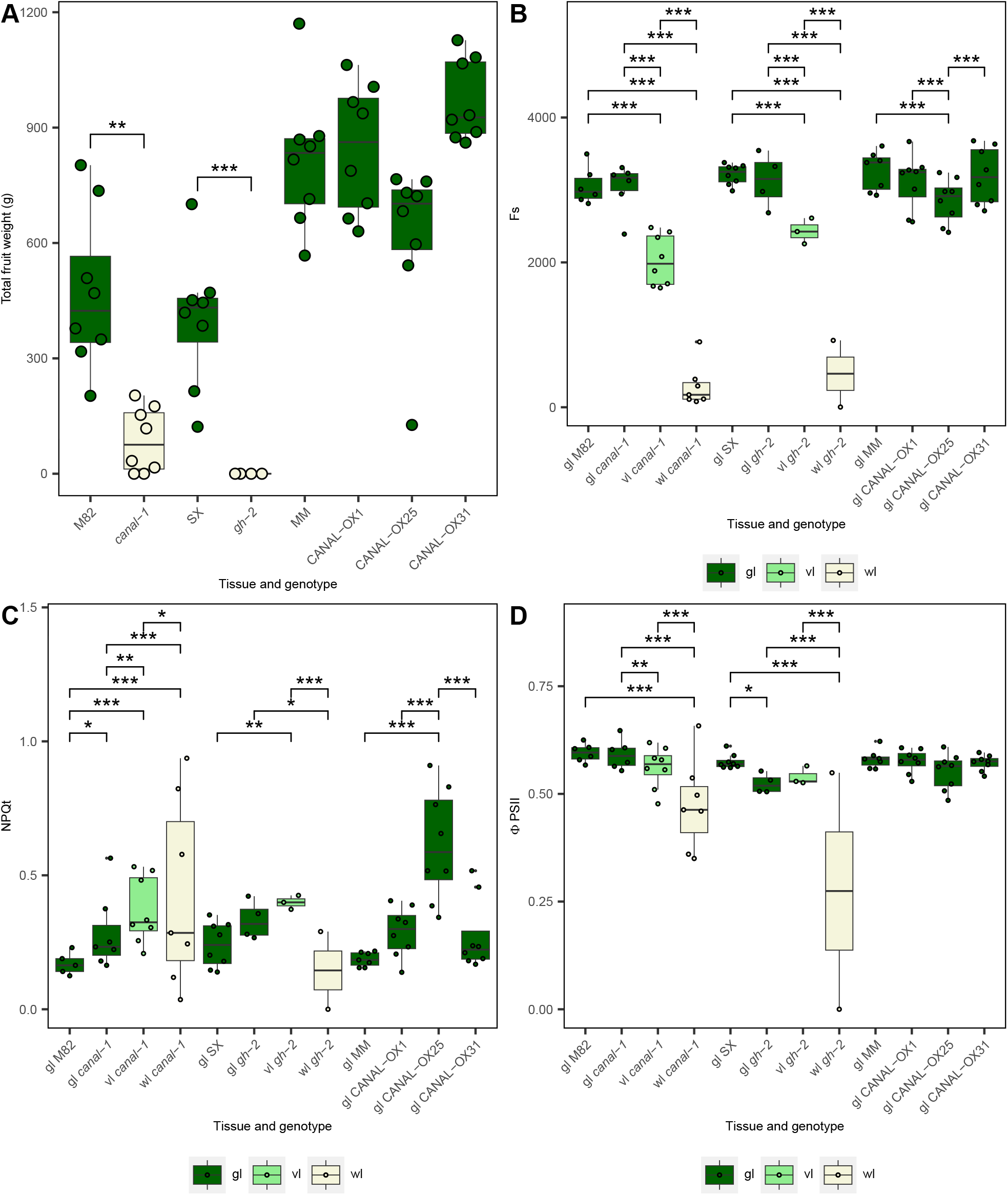
Yield and photosynthetic parameters on wild type, mutants and overexpression lines. A, Total fruit weight (g) of different lines. B, Baseline Fluorescence (Fs). C, Non-photochemical quenching (NPQt). D, Quantum yield of photosystem II (DPSII). Boxes and points are filled according to tissue (gl = green leaves: green, vl = variegated leaves: bright green, white leaves: beige). Statistical significance was estimated by pairwise-t-test for A-D. Significant differences are indicated by a bracket and asterisks (*: p ≤ 0.05; **: p ≤ 0.005; ***: p ≤ 0.0005).

Due to the increased sample size, we did not use the LiCor for photosynthetic measurements in this experiment but rather used a small multispeq handheld device. On the other hand, as an intermediate between green leaves and white leaves, we now also included variegated leaves, i.e. leaves that had both white as well as green sectors, in the measurement. The steady-state fluorescence measurements revealed significantly reduced levels for variegated mutant leaves and an even greater reduction for white leaves, when compared to the green leaves of the corresponding wild-type (Figure 13 B). Green mutant leaves on the other hand showed wild-type fluorescence levels. Interestingly, green leaves from the CANAL-OX25 line also showed a significant reduction in Fs, when compared to MoneyMaker leaves. More or less the opposite effect could be observed for non-photochemical quenching (NPQt) (Figure 13 C). Here, variegated mutant leaves showed an increase in NPQt. White leaves of *canal-1* also showed an NPQt increase, whereas white leaves of *ghost-2* showed a reduction. Finally, the quantum yield of photosystem II was similar to that observed with the LiCor (Figure 13 D), while overexpressor lines were invariant from wild-type plants.

### SCO2 in the context of previously described canalization loci

Although *canal-1* was discovered as an EMS mutant with unstable yield, we were interested, whether we could find any evidence for the yield canalization role in existing data. Field trials of an introgression line population in different seasons, has yielded both metabolomics and yield-related data, which has extensively been used both for mapping of the level as well as the canalization of metabolite levels and also network correlations (Schauer *et al*., 2006, Schauer *et al*., 2008, Alseekh *et al*., 2017, Kuhalskaya *et al*., 2020). The locus Solyc01g108200 to which *canal-1* is associated resides in the introgression line segment IL 1-4. This locus has previously shown a genotype-by-environment interaction on canalization of leucine (Alseekh *et al*., 2017), however yield-related data has not been studied with regards to canalization. We therefore used yield-related data previously gathered for a network-correlation approach for the mapping of canalized yield QTL, with the same procedure described previously (Alseekh *et al*., 2017, Kuhalskaya *et al*., 2020). Interestingly, among 750 tested associations, we could only find two associations between genotype and metabolite canalization with a significant genotype-by-environment effect (Supplemental Data S 3). We found this association between ILs 2-5 and 8-3 and the GxE effect on canalization of harvest index (Supplemental Figure S 12). However, we found no association for IL 1-4, even when not accounting for the environment (Supplemental Data S 4). If we compare the SCO2 gene of the domesticated tomato to its counterpart in the wild tomato species *Solanum pennellii* we can see that there are only six single nucleotide polymorphisms in the coding region, resulting in four amino acid changes, none of which are close to the conserved zinc finger domain (Supplemental Figure S 13 and Supplemental Figure S 14). However, it is important to note that even if these displayed a functional effect it may be masked by the influence of genetic variance in the many other genes carried by the large chromosomal segment substitution. These facts suggest that in future study of this gene across wider natural variance such as that apparent in genome wide association panels may prove more revealing.

## Discussion

In the current study, we presented the molecular characterization of the variegated *canal-1* mutant in tomato. Leaves showed different patterns and degrees of variegation, from almost completely green over variegated to completely white, with white leaves being devoid of photosynthetic pigments, whereas green mutant leaves contained wild-type levels. Transcriptomic analysis revealed the enrichment of photosynthesis-related terms and highlighted upregulation of photosystem assembly and photosystem component genes in green mutant leaves, while white mutant leaves showed wild type levels, with the exception of orthologs of ELIP and DEG genes, which showed high outlying values. These changes seen at the transcriptomic level, were paralleled at the proteome level, with the proteome of green *canal-1* leaves being largely invariant from wild-type in, but white *canal-1* leaves displayed strong deficiencies in proteins related to photosynthesis. Here we did not only see changes in the thylakoid membrane proteins, but also in RuBisCO, which was downregulated in white mutant leaves. The metabolome showed a strong upregulation of amino acids and a downregulation of TAGs and galactolipids in white *canal-1* leaves. The results of our study report similar findings to the earlier more focused studies in *Arabidopsis thaliana* and *Lotus japonica*. Indeed, the tomato *canal-1* mutant more similar resembles the sco2 mutants of Lotus in that their true leaves are affected and display a variegated phenotype under long-day conditions and similarly the levels of PsaB, PsaL, PsbC and PsbD are also decreased in white mutant leaves as was shown for Lotus. By contrast to the results described in Lotus, PsbO and PsbQ are not more abundant at the protein level in tomato. In fact, PsbP - which is also part of the oxygen-evolving complex-was the only differentially abundant protein in green *canal-1* leaves. On the other hand, all three of these proteins (psbO, psbP and PsbQ) show transcriptional upregulation in green leaves but downregulation in white leaves (Figure 6). Also similar to previous results is the reduction of lhcb1 in white *canal-1* leaves, which has been shown on a transcriptomic level in Arabidopsis (Tanz *et al*., 2012) and on a proteomic level in Lotus (Zagari *et al*., 2017). A reduction of D1 and LhcB protein had also been shown for Arabidopsis by an earlier study, which however found little changes in gene expression (Albrecht *et al*., 2008).

Given that correlation analysis clustered the SCO2 transcript closer to assembly genes than to other components of the photosystems, we doubt that SCO2 directly affects transcription or translation of photosystem components and feel that they are more likely to be secondary consequences. Rather, as we discuss in detail below, we feel that SCO2s primary role is in photosystem assembly and that the chloroplast biogenesis may be affected due to the function as assembly component. That said, correlation network analysis showed above median levels of different network metrics (degree, closeness and betweenness), for the SCO2 transcript and high correlation values to a range of other putative photosystem assembly genes. This Furthermore, pathway integration highlighted the increased carbohydrate catabolism in white leaves and confirmed the upregulation of amino acid biosynthetic pathways. Moreover, photosynthetic measurements showed a reduction of activity in variegated and white leaves of *canal-1*, whereas green mutant leaves showed an upregulation of some parameters. Thus, whilst falling short of defining the exact underlying causal mechanisms, we want to highlight two characteristics of our work that distinguish it from the existing literature. First, we have quantitative data for the whole transcriptomeand are able to observe changes in gene expression on a much broader range and with higher precision than this has been done previously (Albrecht *et al*., 2008, Tanz *et al*., 2012) and secondly, we separately analyzed white and green leaves/leaf sectors and could contrast them against each other. This of course was not possible for Arabidopsis but was also not performed for Lotus (Zagari *et al*., 2017). We discuss below the collective results both with respect to the function of SCO2 and with regard to its previous identification as a canalization gene (Fisher and Zamir, 2021).

Due to the changes observed in the multi-omics analysis, we suggest, that an adequate amount of SCO2 activity is indispensable for appropriate thylakoid development. The mutation of the *canal-1* mutant is in close proximity to a conserved zinc finger binding domain that is crucial for catalytic activity (Muranaka *et al*., 2012, Fisher and Zamir, 2021). This is also underlined by the phenotype of other mutants described in literature. Two CRISPR mutants have previously been described (Fisher and Zamir, 2021). The strongest phenotype is displayed by a mutant with an early stop codon before the conserved zinc finger domain, which produces a truncated protein. It has pale cotyledons and fails to build true leaves. Slightly less severe was the phenotype of another CRISPR/Cas9 mutant, with a frame shift that resulted in a change of amino acids 15 to 43. This mutant also developed variegated leaves, but did not survive past the first true leaves. The *ghost-2* mutant, which similarly to *canal-1* is an EMS mutant with a single point mutation, leading to the change of an amino acid within the conserved zinc finger domain. Despite also showing a variegated leaf phenotype, this mutant has been described to not show the unstable variegation phenotype and produce uniformly large plants (Fisher and Zamir, 2021). This is, however, in disagreement with our observation that plants harboring this mutation show a more severe phenotype and are characterized by being unable to produce any fruit. Additionally, despite the increased variation in fruit yield, *canal-1* mutant plants did not vary as strongly in the greenhouse as was observed in the field. Therefore, the discrepancies between our results and the field trials, may be attributed to the effect the different environments had.

The comparison of DNA and amino acid sequences between the cultivated tomato species and the wild tomato species *Solanum pennellii*, revealed few differences, none of which were close to the conserved zinc finger domain. This suggests a strong conservation of the gene which in itself may indicate the important role of SCO2. In addition, no significant association between variation of yield-related traits and the IL segment, harboring the gene could be found. However, as we mention above it will be interesting in future studies to look, in broader panels of genetic variance, at the relationship between individuals harboring various haplotypes of the gene and metabolic and yield associated traits.

The RNAseq experiment did not yield any significant differences in gene expression in the causal gene SCO2 between mutant and wild type leaves suggesting that the protein was expressed and likely transcribed but non-functional. Moreover, despite being able to detect 7729 proteins, which is similar to the number reported in previous experiments or even higher (Kilambi *et al*., 2016, Szymanski *et al*., 2017, Yu *et al*., 2021), we were not able to detect SCO2 among them and can not say whether protein levels were changed. This was most likely just the technical limitation of the proteomics in achieving complete coverage of the whole proteome. More generally however, the upregulation of gene expression, specifically of photosystem assembly components, in green mutant leaves in comparison to wild type leaves, suggests an overcompensation in response to an, at least partially, impaired protein function in SCO2. The correlation network analysis confirmed the high correlation of this expression pattern between the SCO2 transcript and other photosystem assembly components. Our overcompensation hypothesis is further supported by the wild-type-like protein levels in these leaves. Indeed, white *canal-1* leaves showed wild-type transcript levels of most photosystem assembly components. Exceptions to this statement were the upregulation of genes orthologous to ELIP and DEG genes. In de-etiolating seedlings ELIPs were demonstrated to be responsible for the etioplast to chloroplast transition (Casazza *et al*., 2005), however since the true leaves develop in the light, ELIP accumulation may be rather indicative of high photooxidative stress in white leaf sectors (Hutin *et al*., 2003). This could lead white leaf sectors to get trapped in a stage similar to pre-de-etiolation, wherein thylakoid development is arrested and misfolded proteins get degraded by DEG protease. As a consequence of such a scenario proper thylakoids would be unable to develop and protein levels of photosystem components would be significantly reduced. Being unable to photosynthesize, white mutant leaves rather behave as a sink tissues on a metabolic level. This can be seen by the depletion of several mono- and disaccharides like sucrose, maltose, glucose and fructose and the upregulation of enzymes involved in their catabolism in the glycolysis, like PFK and GAPDH. On the other hand, enzymes along the same pathway in the anabolic direction, like GAPA and FBPA are downregulated. By contrast, green mutant leaves seem to be able to compensate for the lack of photosynthates by an upregulation of photosynthesis and an altered carbon allocation. This can also be seen for example in the enzymes GAPA and FBPA showing the opposite effect than white leaves towards an increased sugar anabolism. The upregulation of amino acids is likely a secondary effect, as a general response to stress, which is supported by transcriptomic and proteomic changes in enzymes involved in the biosynthesis of amino acids, like ASNS, GOGAT, ILVC, ILVD and ILVE. Downregulation of TAGs and galactolipids may be related to the missing thylakoid membranes. The increased photosynthesis in green mutant leaves is displayed by a higher assimilation rate. This in turn, however, requires green leaves to open their stomata wider, which consequently comes at the cost of increased transpiration.

The compensation response described above fits to a threshold model described for other variegation mutants (Wu *et al*., 1999, Aluru *et al*., 2006). This model suggests a base-level of either protein function or abundance, to be necessary for the development of functioning photosystems. Furthermore, it is well known that the stoichiometry of proteins is important for thylakoid development (Zagari *et al*., 2017). In this regard, SCO2 has previously been described as a “scale” that balances different photosystem components (Zagari *et al*., 2017). All of this applied to the observations in the *canal-1* mutant, could mean that there is a certain critical level of SCO2 function, which if surpassed enables the formation of thylakoids and consecutively the development of green leaf sectors. If this level is however undershot, thylakoid formation is not achieved and leaf sectors stagnate in an undeveloped state. This may be exacerbated by the fact that at least photosystem II is especially vulnerable to photodamage in early development (Shevela *et al*., 2019). It seems reasonable to suggest that the larger the total area of photosynthetically active tissue, the larger the fruit mass it can sustain. Conversely, if large amounts of the leaf area remain white, only a low fruit yield can be produced a fact that is aggravated by if our supposition that white leaves act as additional sink tissues is correct. Ergo, statistically speaking, the instable yield phenotype in comparison to wild-type plants, is likely founded in an increased variation of photosynthetic active tissue, which in the wild type usually equals close to 100%. This theory could also explain, why our results from greenhouse experiments were different from those obtained in the field. Light intensity is usually much higher in the field in comparison to a greenhouse. For young mutant plants, with an impaired photosystem assembly protein, this may present a stress factor, which could early on lead to the development of white sectors. Additionally, a necessary increase in stomatal conductance in mutant green leaves, likely makes mutant plants more vulnerable to heat and drought stress. This may be an additional fact that explains the large variance of small and large plants in the field.

As we are interested in SCO2 as a canalization gene, we should put it into the context of known canalization genes. A general idea for canalization genes has been suggested, that they are so-called hub genes (Laitinen and Nikoloski, 2019). The most well-described canalization gene, that is such a hub gene, is the chaperone heat shock factor 90 (*HSP90*)(Lachowiec *et al*., 2016). HSP90 interacts with many different client proteins and facilitates their proper folding (Schopf *et al*., 2017). Initially in *Drosophila melanogaster*, it could be shown that impairment of HSP90 leads to an increase in phenotypic variance (Rutherford and Lindquist, 1998). Later it could also be shown that reduction of HSP90 function in *Arabidopsis thaliana* by pharmacological treatment or by RNA interference leads to an increase in frequency of atypical phenotypes, putatively by the release of cryptic genetic variation (Queitsch *et al*., 2002, Sangster *et al*., 2007). A study in *C. elegans* found several DnaJ proteins that affect canalization of seam cell number (Hughes *et al*., 2019). This study found that not all of the DnaJ genes were expressed in seam cells and it was concluded that they are rather part of a larger gene network. Another hub gene, which was found to affect the variation of glucosinolate levels, circadian periodicity as well as flowering time, is the gene *EARLY FLOWERING 3* (*ELF3*), which is known to be generally involved in the circadian pathway (Hicks *et al*., 2001, Jimenez-Gomez *et al*., 2011). Similarly, the *ERECTA* gene, which is a known pleiotropic developmental regulator, was suggested to influence canalization of rosette leaf number (Hall *et al*., 2007, van Zanten *et al*., 2009). Based on these examples, we could say that there are some hub genes, which are more directly involved in specific gene networks as well as those, like HSP90, that are only peripherally connected but affect many processes (Lachowiec *et al*., 2016).

In congruence with these examples, we can see several overlaps to SCO2. Firstly, on the molecular level SCO2 is also a DnaJ-like chaperone that is suggested to interact with other proteins (Shimada *et al*., 2007, Tanz *et al*., 2012). The demonstrated catalytic activity of SCO2 as a disulfide isomerase further suggests it might accelerate the folding of photosystem subunits, which contain cysteine residues (Muranaka *et al*., 2012). Even though the specific function of SCO2 in thylakoid assembly, would qualify it as a gene of a specific gene network, we know that it also has a global effect on yield (Fisher and Zamir, 2021). We would also argue that SCO2 is involved in a larger gene network, which extends beyond its immediate interaction partners. Our correlation analysis has shown, that SCO2 has 29 direct neighbors, which makes its node degree already significantly higher than the median node degree of the correlation network, again highlighting the role of SCO2 as a hub gene. Beyond that SCO2 also displayed a higher betweenness than the median of the network, suggesting its involvement in a larger network. Our results also revealed that SCO2 expression correlates more closely with the expression of other putative assembly factors and only more distantly to photosystem components. SCO2 transcript showed the highest positive correlation to Solyc09g010110, an ortholog to the ORANGE gene in Arabidopsis, which has been shown to play a key role in chromoplast differentiation by posttranscriptionally regulating phytoene synthase (Zhou *et al*., 2015). Similarly, Solyc09g010110 in tomato (also called OR-like) has also been suggested to play an important role in tomato fruit ripening and carotenoid accumulation (D’Andrea *et al*., 2018). Here, it is suggested that OR and OR-like, as plastidial chaperones, could fulfill multiple roles to ensure enzyme availability and prevent carotenoid degradation (D’Andrea *et al*., 2018). Other interesting genes that showed a high correlation to SCO2 transcripts were orthologs of Ycf4 and PAM68, which are known factors in the assembly of photosystem I and II respectively (Yang *et al*., 2015, Lu, 2016a). Ycf4 binds to different PSI subunits, but in tobacco seems to be non-essential for photosystem I assembly (Krech *et al*., 2012). PAM68 has been shown to be involved in early PSII assembly, interacting both with early assembly intermediates as well as other assembly factors (Armbruster *et al*., 2010). Similarly, SCO2 may fulfill analogous roles to ensure the proper folding of photosystem proteins during photosystem assembly or repair. The proper establishment of photosystem components, in which SCO2 likely plays an important role, could be described as a multi-layered regulation mechanism as detailed below.

The proper assembly of photosystem components requires a tight coordination between nucleus and plastid on multiple levels (Pogson and Albrecht, 2011). On a first level, during chloroplast biogenesis, there is a two-phase transcriptional regulation, with anterograde and retrograde signaling between nuclear and plastid genome (Dubreuil *et al*., 2018). Through a second layer of control, the translation of some photosystem subunits is autoregulated by the assembly state, to ensure the correct stoichiometry (Zoschke and Bock, 2018). As mentioned above SCO2 has been assigned a similar role balancing the correct ratio of PSI/PSII complexes and LHC proteins in *Arabidopsis thaliana* and *Lotus japonicus* (Zagari *et al*., 2017) and our results suggest a similar mechanism in tomato. As this step concerns readily translated proteins, this could be regarded as a third layer to ensure appropriate stoichiometry, where SCO2 plays a key role. It therefore seems plausible that reduced SCO2 function could decanalize the development of proper photosynthetic tissue and as a consequence increase variation of plant yield.

Although we are largely satisfied with our explanation of how SCO2 affects canalization of plant yield, further more targeted experiments with higher spatial and temporal resolution would likely be required to gain a more precise understanding of the exact mechanistic role of the SCO2 protein. Nevertheless, our work underlines the importance of the SCO2 protein in thylakoid formation and explains the differences between white and green sectors of the *canal-1* mutant, by different responses on a transcriptome, proteome and metabolome level as well as physiological response. It also largely explains how SCO2 can stabilize both metabolism and morphology. It is our contention that our work alongsided the earlier work on Hsp90 suggest that other chaperone-like proteins may be interesting candidate genes as a means to gain a fuller understanding of how this important and highly agriculturally relevant phenomenon is achieved.

## Experimental procedures

### Cloning and transformation

The gateway cloning system was used to create vectors expressing SCO2 fused C-terminally to eGFP under the 35S-promotor. To achieve this cDNA was generated from tomato leaf tissue with the maxima first strand cDNA synthesis kit (ThermoFisher Scientific, Waltham MA USA) and used as template for a two-step PCR using Phusion DNA Polymerase (ThermoFisher Scientific, Waltham MA USA) to add adapter sequences to the PCR product. This PCR product was then used in a BP reaction with pDONR221 and then in an LR reaction with pK7FWG2 to create the final construct. *Agrobacterium tumefaciens* GV2260 was transformed with the final construct via electroporation. Tomato plants of the cultivar MoneyMaker were generated via agrobacterium-mediated plant transformation as a service of the green team of the MPIMP, similar to published protocols. Here, calli are regenerated from cotyledon segments, which are inoculated with the transformed *A. tumefaciens* and transferred to fresh medium several times, followed by induction of roots and shoots. Shoots of young putatively transformed plants, were cut, transferred to fresh MS cultivation medium (0.68% (w/v) agar, 2% sucrose (w/v)) containing beta-bactyl (125 mg/l; (w/v) and regenerated for 6 weeks. This was repeated at least two times to prevent the release of transgenic *A. tumefaciens* into the wild and to ensure that positive PCR products are no false negatives. Elimination of *A. tumefaciens* was confirmed by PCR amplification of the chromosomal rpoB gene. Successful transformation of putative transformants was achieved by PCR amplification of a portion of the transformed construct, integrated into the plant genome. PCR positive, agrobacterium-free, T0 plants were transferred to the greenhouse for propagation.

### Plant material and growth conditions

The *canal-1* mutant was described previously and seeds were supplied by the Zamir Lab (Fisher and Zamir, 2021). The *ghost-2* mutant, together with the Sioux wild-type plant was ordered from the TGRC. Plants overexpressing the SCO2 protein were generated as described above and used in the generation T2 for experiments. Plants were grown in multiple seasons in different location. The first growth season was 2017 in a greenhouse in Ashkelon Israel on so-called lysimeter assay, which is a high-throughput screening system that monitors water relations for each plant (Halperin *et al*., 2017). Seeds of *canal-1* plants and M82 plants were sown at Histill nursery, Ashkelon in September 3^rd^ 2017 and supplied with organic treatment, devoid of dwarfing hormones. The experiment started October 3^rd^ 2017 and tissue samples were collected on November 6^th^ 2017.

Consecutive experiments were conducted at the MPIMP in Potsdam-Golm, Germany and plants were grown as described previously (Wijesingha Ahchige *et al*., 2023). Plants were grown 2018 and 2019 in a greenhouse. The experiment including plants overexpressing the SCO2 protein was conducted in 2021. Seeds were directly sown to soil, kept in a growth chamber (York International/Johnson Controls; Cork Ireland), in a (16h/8h)-(22°C/18°C)-(70% RH/70% RH)-(day/night)-cycle, with 150 µmol m^-2^ s^-1^ of additional light. Plants were transferred to individual pots after cotyledons were fully expanded. After four weeks, plants were potted into 18 cm diameter pots and transferred to a greenhouse chamber. Here plants were grown in a (16h/8h)-(22°C/20°C)-(50% RH/50% RH)-(day/night)-cycle and automatically irrigated several times a day by a drip irrigation system. A subset of the plants grown in 2019 received a drought treatment, comprising 50% of the optimal irrigation. Sodium discharge lamps supplied additional light in order to compensate for seasonal differences in light intensity. Plants were fertilized once after potting, onset of flowering and onset of fruit ripening.

### Phenotyping

During the growth period, plants were checked three times a week for the first flower and the first red ripe fruit. For any plants that had not produced fruit at the end of the experiment, we assigned the date of the next regular fruit checking as the date for red ripe fruit. Fruits were harvested when around 80% of fruits were ripe. At harvest date, the number of all ripe and unripe fruits per plant was counted and the total fruit weight determined on a scale (Sartorius, Göttingen, Germany). Average fruit weight, was determined by dividing the total fruit weight per plant by the total fruit number per plant.

### Sampling

Mature non-senescing leaves were sampled and directly snap-frozen in liquid nitrogen. For fruit samples, a section of the pericarp was cut out of a whole red ripe tomato fruit. The cuticle of the fruit was removed and the sample was directly snap frozen in liquid nitrogen. All samples were stored at -80°C until further processing.

### RNA extraction

Raw RNA was extracted with a lithium chloride precipitation protocol, similar to a published protocol (Bugos *et al*., 1995).

A Retsch mill (Retsch, Haan, Germany) was used to grind plant tissue to fine powder and 100 mg was weighed into a tube. To frozen plant tissue 280 µl extraction buffer (100 mM TRIS HCl pH 9, 200 mM NaCl, 15 mM EDTA pH8, 0.5% sarcosyl (v/v), 0.8 % β-mercaptoethanol (v/v)) was added and the sample vortexed. After that first 280 µl phenol, then 56 µl chlorofomisoamylalchohol (24:1; v/v) and 19.6 µl 3M NaOAc (PH 5.2) were added, vortexing in between steps. Samples were incubated 15 minutes on ice and centrifuged for 10 min at 10621 rcf. The supernatant was transferred to a new tube and 280 µl phenol-chloroform-isoamylalcohol (25:24:1, v/v/v) was added. After vortexing, the sample was centrifuged for 10 min at 10621 rcf. The upper phase was transferred to a new tube and the previous step was repeated until the white interphase was not visible any more. After adding 280 µl isopropanol, samples were incubated for 30 minutes on ice to precipitate RNA. Samples were then centrifuged at 10621 rcf and 4°C for 10 minutes before discarding the supernatant and washing the pellet with 500 µl 80% ethanol/DEPC. The pellet was then air-dried at room temperature and resuspended in 200 µl sterile H2O/DEPC, before centrifuging the sample for 5 minutes at 10621 rcf and 4°C to remove any particulate. The supernatant was transferred to a new tube, to which 100 µl 8M LiCl was added. The sample was mixed by carefully inverting the tube. The sample was then incubated overnight. After that precipitated RNA was collected by centrifuging the sample 15 minutes at 10621 rcf and 4°C. The pellet was washed once with 250 µl 70% ethanol/DEPC before discarding the supernatant and eluting the pellet in 50 µl H2O/DEPC. A NanoDrop_TM_ One spectrophotometer (ThermoFisher Scientific, Waltham, MA, U.S.A.) was used to estimate RNA concentration. For the digestion of residual DNA, 10 µg of raw RNA was digested with the Invitrogen Turbo DNAse free kit (Invitrogen/Thermo Fisher Scientific, Waltham, MA, U.S.A.). The Bioanalyzer 2100 (Agilent; Santa Clara, CA, U.S.A.) was used to determine RNA quality and quantity.

### Orthology

For orthology search, OrthoFinder v2.5.2 (Emms and Kelly, 2019) was utilized, using the diamond_ultra_sens algorithm with proteomes using only the primary transcript of *Arabidopsis thaliana* Araport11, tomato (*Solanum lycopersicum*) ITAG4.0, potato (*Solanum tuberosum* v6.1), *Lotus japonicus* Lj1.0v1, maize (*Zea mays* RefGen_V4) and rice (*Oryza sativa* v7.0) obtained from Phytozome (Ouyang et al., 2007; Jiao et al., 2017; Li et al., 2020; Hosmani et al., 2019; Cheng et al., 2017; Pham et al., 2020). Orthologs of photosynthetic genes were found by matching genes from *Arabidopsis thaliana* found in literature (Hankamer *et al*., 1997, Jensen *et al*., 2007, Shi *et al*., 2012, Berardini *et al*., 2015, Yang *et al*., 2015, Lu, 2016b) to the discovered orthogroups. A resolved gene tree of the glutaredoxin orthogroup was constructed with ggtree (Yu, 2020).

### Protein visualization

Putative protein structure was obtained from AlphaFold (Jumper et al., 2021; Varadi et al., 2022) and visualized with Mol* Viewer (Sehnal et al., 2021).

### RNAseq analysis

Library preparation and RNA sequencing was performed by the MPIPZ Cologne. Libraries were prepared by a polyA enrichment. Reads were acquired by a HiSeq3000 sequencer (Illumina, CA, U.S.A.) as 2 x 150 bp paired-end reads, with a sequencing depth of 20,000,000 reads. Raw reads were mapped to the nuclear and organellar genome via LSTrAP (Proost *et al*., 2017). Differential gene expression was performed with deseq2. Genes with an FDR-adjusted p-value ≤ 0.1 and a log2-fold-change ≥ 1 were considered differentially expressed based on recommendations in the literature (Conesa *et al*., 2016, Lamarre *et al*., 2018). The expression matrix as produced by LSTrAP and a deseq2-normalized expression matrix together with log2-fold changes and adjusted p-values of the respective contrasts (Supplemental Data S 5 + Supplemental Data S 6).

Solgenomics Gene ID to Entrez Gene ID correspondence was inferred by aligning the ITAG4.0 proteome against the NCBI Refseq SL3.1 proteome with Diamond v.0.9.9. Alignments were filtered for 100% identity and 95% query cover as well as being on the same chromosome in both assemblies. Solgenomics Gene IDs mapping to more than one Entrez Gene ID were ranked based on the absolute distance of their start regions and only the entry with the smallest distance was retained. A correspondence table can be found with the supporting information (7) Gene ontology enrichment analysis and gene set enrichment analysis (GSEA) was performed with the help of the packages topGO and fgsea. Bubble plots were inspired by the GOplot package (Walter *et al*., 2015). Accordingly the z-score was calculated as detailed below (Formula 1).

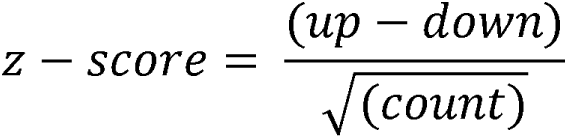

Formula 1: Z-score calculation for bubble plots. up: number of upregulated genes, down: number of downregulated genes, count: total number of genes

### Metabolite MTBE extraction

Plant tissue was ground to fine powder with a mixing mill (Retsch; Haan, Germany). Here, plant tissue was added to a pre-cooled stainless steel grinding jar loaded with a 20 mm steel ball and milled for 1 min at 30 Hz. Aliquots of 50 mg plant tissue powder were weighed with a fine scale and used for combined extraction of metabolites, proteins and pigments according to a previously described protocol (Salem *et al*., 2020). To the ground tissue aliquots 1 ml of pre-cooled MTBE:MeOH (3:1, v/v) containing internal standards, was added vortexed before shaking the samples at 4°C for 30 minutes. Samples were then sonicated in an ice-cooled sonication bath for 15 minutes, before centrifuging them at 4°C and 10000 g for 10 minutes. From the supernatant, 500 µl was transferred to a new tube and 500 µl of H_2_O:MeOH (3:1, v/v) was added and the sample vortexed briefly. After centrifugation at 4°C and 10000 g for 10 minutes 250 µl of the upper phase was collected for lipid analysis. For samples, for which pigments were analyzed a second aliquot of 100 µl was taken. The BVC fluid aspiration system (Vacuubrand Inc: Essex, CT, U.S.A) was used to remove the remainder of the upper phase, before taking aliquots of the lower (apolar) phase. From this phase, aliquots of 200 µl and 250 µl were taken for GC- and LC-MS respectively. Extracts were dried for 3 h or overnight in a Scan Speed 40 centrifugal vacuum concentrator (Labogene; Allerød, Denmark), coupled to a Scanvac CoolSafe cryo unit (Labogene; Allerød, Denmark) at 30°C and 1000g for 3h or overnight. Until further use, dried extracts were stored at -80°C. Samples from 2017 were initially extracted with a modified version of a different protocol also described before (Lisec *et al*., 2006). Here, 730 µl of MeOH containing 30 µl of the internal standard ribitol were added to the frozen tissue sample and the sample was vortexed. The samples were shaken at 70°C for 15 minutes, before centrifuging the samples for at least 20 minutes at 20817 rcf.

From the supernatant 684 µl was transferred to a new tube. Samples for which pigments were analyzed 584 µl was transferred to another new tube, leaving an aliquot of 100 µl for pigment analysis. The addition of further components was adjusted accordingly (see brackets). Now, 375 (300) µl of chloroform and then 750 (600) µl H2O was added before vortexing each sample carefully. Samples were centrifuged for 10 minutes at 20817 rcf and two aliquots were taken from the upper phase for GC- and LC-MS. Aliquots were dried in a speed vacuum concentrator without heating overnight.

### Pigment determination

The aliquot of 100 µL taken from the lipophilic phase of the MTBE extraction was diluted 10-fold with methanol and the extracts were measured under an Epoch2 96-well plate reader (Biotek/Agilent; Santa Clara, CA, U.S.A.). Spectrum was measured at 470 nm, 652 nm and 665 nm. Assuming a pure methanolic extract pigment concentration was calculated with a pathlength correction of 0.51 as described previously (Lichtenthaler and Buschmann, 2001, Warren, 2008).

### Proteomics

Proteins were purified from the pellet of the MTBE extraction according to previously published protocols (Sokolowska *et al*., 2019, Salem *et al*., 2020). Here the protein pellet was dissolved in 50 µL denaturation buffer (50 mM Ammonium bicarbonate pH 8 (Ambic), 2M Thiourea, 6M Urea) and sonicated in a sonication bath for 15 minutes. After determining protein concentration by Bradford Assay (Bio-rad, Hercules, CA, USA), a volume containing 50 µg of protein was added to a new tube and filled to 20 µl with denaturation buffer. After adding 2 µl of reduction buffer (5 mM DTT) samples were incubated for 30 minutes at room temperature. Samples were then incubated for another 30 minutes at room temperature before adding 2 µl alkylation buffer (30 mM iodoacetamide). To the samples 5 µl LysC/Trypsin mix (50 mM AmBic; 20 µg protein) was added and an additional 15 µl AmBic buffer (50 mM Ambic pH 8) was added to achieve a final urea concentration of 4 M. Samples were then incubated for 2 h at 37°C. After adding 120 µl Ambic buffer to achieve a urea concentration of 1 M, samples were incubated at 37°C overnight. After that, samples were acidified with 1.65 µl Trifluoroacetic acid (TFA) to a final TFA concentration of 1% (v/v).

For protein desalting Sep-Pak filter columns were placed on top of a vacuum manifold, washed first with 1 ml MeOH, then with 1 ml Solution B (0.1 % TFA (v/v), 80 % acetonitrile (v/v)) and then two times with 1 ml Solution A (0.1 % TFA (v/v)), making sure to apply vacuum until all liquid passed through the column between each wash. Samples were then loaded onto columns and vacuum was applied until all liquid had passed through the column. Columns were again washed two times with 1 ml Solution A, letting all liquid pass through the column, before placing 2 ml tubes under the outlets. Peptides were eluted with 600 µl Solution C (0.1 % TFA (v/v), 60% acetonitrile (v/v)) and the flow through was collected. Peptides were transferred to 1.5 ml tubes and dried in the Speed vacuum concentrator for 3 hours and stored at -20°C until further use.

Proteomics mass spectrometry was performed as described before (Alseekh *et al*., 2022). Isolated peptides were resuspended with MS loading buffer and run on an ACQUITY UPLC M-Class System (Waters, Wilmslow UK) coupled to a Q Exactive HF Orbitrap mass spectrometer (Thermo Fisher Scientific, Waltham MA, USA) Proteomics data were analyzed with the MaxQuant software (Tyanova *et al*., 2016) and the tidyproteomics package (Jones *et al*., 2023). Missing values were imputed by the half minimum per protein and differentially expressed proteins were estimated with two-sided t-tests after normalization with limma (Ritchie *et al*., 2015). Proteins were considered differentially expressed at a log2-fold-change ≥ 1 and an FDR-adjusted p-value ≤ 0.1. Protein groups, that contained multiple proteins were split into all individual proteins. No filtering was performed by the number of unique peptides. Relationship of uniprot IDs to Entrez IDs/KEGG IDs were established from the uniprot proteome UP000004994, downloaded on the 07.09.2022. Proteins that were meanwhile replaced by newer versions were updated (Protein B6ECN9 meanwhile replaced A0A3Q7FS73). For downstream pathway visualization, duplicates were ranked based on p-values of contrasts and the highest ranking entries per duplicate were retained. Enrichment analysis was performed with the packages topGO and clusterProfiler. The raw data as produced by the MaxQuant software as well as table showing raw intensities together with log2-foldchanges and p-values can be found in the Supporting information.

### Metabolomics

Primary metabolites were measured via gas chromatography-mass spectrometry (GC-MS) according to previously used protocols (Lisec *et al*., 2006). For primary metabolites, extracts stored at -80°C were again dried in a speed vacuum concentrator for 10-30 minutes before derivatization treatment. To each sample 40 µl of Methoxyaminhydrochlorid (20 mg/ml Pyridine) was added, samples were shaken for 2 hours at 37°C and 1100 rpm. After briefly spinning down any condensed droplets, 70 µl of MSTFA mix (1 ml MSTFA + 20 µl fatty acid methyl esters (FAMEs)) was added. Samples were again shaken for 30 minutes at 37°C and 1100 rpm and spun down. From the supernatant 100 µl was transferred to a sample vial. Via a Gerstel Multipurpose autosampler (Gerstel GmbH & Co.KG, Mühlheim an der Ruhr, Germany), samples were injected and separated via gas chromatography on an Agilent 7890 B gas chromatograph (Agilent, Santa Clara, CA, U.S.A.). Coupled to this was a Pegasus HT TOFMS mass spectrometer (Leco Corporation; St. Joseph, MI, U.S.A.). Peaks were picked manually in Xcalibur, according to an in-house processing method. Raw peak area was normalized to the internal standard ribitol.

Secondary metabolites were measured via polar liquid chromatography-mass spectrometry (LC-MS) according to literature (Giavalisco *et al*., 2009). Here, dried extracts were resuspended in 100 µl methanol:water (1:1, v/v) and samples were vortexed and then sonicated for 10 minutes. After centrifuging samples for 5 minutes at 10000 g, 80 µl was transferred to glass vials. From here, samples were injected onto an Acquity UPLC system (Waters Corporation, Milford, MA, U.S.A) and analyzed by an Exactive Orbitrap mass detector (ThermoFisher Scientific, Waltham, MA, U.S.A.), using a heated electrospray source (ThermoFisher Scientific, Waltham, MA, U.S.A.). Negative ionization mode was used to obtain mass spectra.

Lipophilic compounds were measured via apolar (LC-MS) according to literature (Hummel *et al*., 2011). To dried lipophilic extracts, 100 µl acetonitrile:isopropanol (7:3, v/v) was added. Samples were vortexed, sonicated for 10 minutes and centrifuged for 5 minutes at 10000g before transferring 80 µl to sample vials. From here, samples were injected to a Orbitrap high-resolution mass spectrometer:Fourier-transform mass spectrometer (FT-MS) attached to a linear ion trap (LTQ) Orbitrap XL (ThermoFisher Scientific, Waltham, MA, U.S.A.). To obtain mass spectra, samples were run in positive ionization mode (Alseekh *et al*., 2018). Peak area for secondary and lipophilic compounds were picked by Genedata Expressionist ® (Genedata, Basel, Switzerland).

Metabolites were annotated according to in-house libraries, the Golm Metabolome Database (GMD, http://gmd.mpimp-golm.mpg.de/) and literature (Larbat *et al*., 2014, Szymański *et al*., 2020, Omidbakhshfard *et al*., 2021). Heatmaps were created with the pheatmap package (Kolde, 2019). Metabolic pathway enrichment was performed with the web-version of MetaboAnalyst 5.0 (https://www.metaboanalyst.ca/MetaboAnalyst/home.xhtml).

Metabolite values from samples extracted from the same source material in 2017 and 2021 were averaged, after normalization to wild type samples, as they constitute technical replicates. All metabolite data are reported following recently updated metabolite reporting standards Supplemental Data S 1 (Fernie et al, 2011; Alseekh et al., 2021).

### Photosynthesis measurements

Photosynthetic parameters were estimated in two independent experiments. The first measurement was done on control plants in the experiment from 2019, using a LI-6400-40 fluorometer (LI-COR Biosciences GmbH, Bad Homburg, Germany). Measurements were done under 400 µE m^-2^ s^-1^. The second measurement was done on plants in the experiment from 2021 including overexpressing plants, using a handheld MultispeQ device V2.0 (Photosynq Inc., East Lansing, MI, U.S.A.)(Kuhlgert *et al*., 2016). Measurements were performed under ambient light, using the Rides 2.0 protocol.

### Omics integration

Different omics datasets were integrated by the help of Pathview (Luo and Brouwer, 2013) and SBGNview (Dong *et al*., 2022). First Vanted (v2.8.3) (Junker *et al*., 2006) and the SBGN-ED plugin (Czauderna *et al*., 2010) were used to download KEGG pathway maps, which were transformed into SBGN process description maps. KEGG REST API was used for gene and compound mapping (https://www.kegg.jp/kegg/rest/keggapi.html). For the glycine cleavage system (EC 1.4.1.27) we manually combined the ECs 1.4.4.2, 1.8.1.4 and 2.1.2.10 and their corresponding KEGG orthologies. Pathway maps were labelled accordingly and their layout optimized with the help of Vanted and newt editor (Junker *et al*., 2006, Sari *et al*., 2015, Balci *et al*., 2021). Finally, omics data was mapped to pathway maps with the SBGNview package (Dong *et al*., 2022) with minor modifications currently forked to https://github.com/micwij/SBGNview.

### Correlation network

Correlation networks were constructed by calculating the Spearman-correlation value for the 15 samples for which all omics-data were available. Different thresholds were used for different networks, but to reduce computational burden correlation values below 0.75 were directly filtered out, when gathering data. Correlation networks were generated with igraph and network graphs were created with the help of ggraph and graphlayouts (Csardi and Nepusz, 2005, Pedersen, 2022, Schoch, 2023).

### Computational analysis, used packages, webtools and software

Computational analysis and visualization was performed with R statistical software v4.2.1 in the Rstudio environment or on our in-house high performance computing platform. In addition to previously mentioned packages we also used the following packages for data handling and visualization: tidyverse, ggtext, ggpubr, gggenes, ggrepel, ggbeeswarm, cowplot, openxlsx, broom, car, deseq2, apeglm, topGO, fgsea, clusterprofiler, viridisLite, lme4, broom.mixed, here, extrafont, magick (Love *et al*., 2014, Fox and Weisberg, 2019, Zhu *et al*., 2019, Müller, 2020, Wilke, 2020, Wilkins, 2020, Alexa and Rahnenfuhrer, 2021, Korotkevich *et al*., 2021, Wu *et al*., 2021, Bolker and Robinson, 2022, Clarke *et al*., 2022, Garnier *et al*., 2022, Larsson, 2022, Wilke and Wiernik, 2022, Kassambara, 2023, Ooms, 2023, Robinson *et al*., 2023, Schauberger and Walker, 2023, Slowikowski, 2023). Benchling [Biology Software] (https://benchling.com) was used for sequence alignments. Serial Cloner v2.6.1 (SerialBasics http://serialbasics.free.fr/Serial_Cloner.html) and Benchling [Biology Software] (https://benchling.com) were used to plan cloning of constructs. Colors were picked from a colorblind-friendly scale as suggested previously (Wong, 2011) with the help of a webtool (https://davidmathlogic.com/colorblind/#%23D81B60-%231E88E5-%23FFC107-%23004D40), which was also used to pick additional colors. The bibtex package was used to export R package citations into .bib files (Francois and Hernangómez, 2023). In the writing and editing of this manuscript Grammarly (©Grammarly Inc., CA, USA) was used to check grammar, interpunctation and improve readability of the text.

## Statistical analysis

As many experiments had an uneven number of samples a pairwise-wilcox test was performed. Equally distributed samples were analyzed by two-sided t-tests. Unless otherwise stated unadjusted p-values are shown.

## Data Statement

All raw data, scripts, workflow descriptions and metadata are publicly available in an annotated research context (ARC) which will be published under a CC-BY 4.0 license in the DataHUB of the DataPLANT consortium for FAIR data management (Weil *et al*., 2023) at the time of publication. The tools SWATE and ARCCommander were used to create and manage the ARC.

## Accession Numbers

Sequence data from this article can be found in the solgenomics database. Accession numbers of genes and proteins are listed in the supporting information Supplemental Data S 5 - Supplemental Data S 9.

## Acknowledgments

We would like to thank Menachem Moshelion for setting up the Lysimeter assay. Further we would like to thank Karin Köhl and the green team for plant transformation and excellent plant care. We would also like to thank José Vallarino for help with LI-COR measurements. We would like to acknowledge the RESIST project for enabling the cooperation between the MPIMP and the University of Cape Town. We would also like acknowledge the support of the DataPLANT consortium and the German Research foundation (DFG) through grant number NFDI 7/1. A.R.F. and S.A. acknowledge the European Union’s Horizon 2020 research and innovation programme, project PlantaSYST (SGA-CSA No. 739582 under FPA No. 664620), and the BG05M2OP001-1.003-001-C01 project, financed by the European Regional Development Fund through the Bulgarian’ Science and Education for Smart Growth’ Operational Programme. S.A. acknowledges the NatGenCrop project: HORIZON-WIDERA-2022-TALENTS-01, No. 101087091.

## Supporting information

Supplemental Table S 1: Enriched terms in GO enrichment analysis of transcriptome with topGO comparing wild type green leaves versus white *canal-1* leaves.

Supplemental Table S 2: Enriched terms in GO enrichment analysis of transcriptome with topGO comparing green *canal-1* leaves versus white *canal-1* leaves.

Supplemental Table S 3: Enriched terms in GO enrichment analysis of transcriptome with topGO comparing wild type green leaves versus green *canal-1* leaves.

Supplemental Table S 4: Enriched terms in GO enrichment analysis of transcriptome with topGO comparing wild type stems versus *canal-1* stems.

Supplemental Table S 5: List of tomato genes orthologous to known photosynthesis genes in *Arabidopsis thaliana*.

Supplemental Table S 6: Enriched terms in gene set enrichment analysis of transcriptome with fgsea comparing wild type green leaves versus white *canal-1* leaves.

Supplemental Table S 7: Enriched terms in gene set enrichment analysis of transcriptome with fgsea comparing green *canal-1* leaves versus white *canal-1* leaves.

Supplemental Table S 8: Enriched terms in gene set enrichment analysis of transcriptome with fgsea comparing wild type green leaves versus green *canal-1* leaves.

Supplemental Table S 9: Enriched terms in gene set enrichment analysis of transcriptome with fgsea comparing wild type stems versus *canal-1* stems.

Supplemental Table S 10: Enriched terms in GO enrichment analysis of proteome with topGO comparing wild type green leaves versus white *canal-1* leaves.

Supplemental Table S 11: Enriched terms in GO enrichment analysis of proteome with topGO comparing *canal-1* green leaves versus white *canal-1* leaves.

Supplemental Table S 12: Enriched terms in KEGG enrichment analysis of proteome with clusterProfiler comparing wild type green leaves versus white *canal-1* leaves.

Supplemental Table S 13: Enriched terms in KEGG enrichment analysis of proteome with clusterProfiler comparing *canal-1* green leaves versus white *canal-1* leaves.

Supplemental Table S 14: Enriched pathways in enrichment analysis of metabolic pathways with Metaboanalyst comparing *canal-1* green leaves versus white *canal-1* leaves.

Supplemental Table S 15: Enriched pathways in enrichment analysis of metabolic pathways with Metaboanalyst comparing wild type green leaves versus white *canal-1* leaves.

Supplemental Table S 16: Results of wilcox-tests on network metrics degree, closeness and betweenness, comparing SCO2 value to the overall filtered network.

Supplemental Figure S 1: Pigments in stems of wild type and *canal-1* plants. Supplemental Figure S 2: Principal component analysis of RNAseq data.

Supplemental Figure S 3: Differentially expressed genes in different contrasts.

Supplemental Figure S 4: Volcano plot of differentially expressed genes comparing wild type and *canal-1* leaves.

Supplemental Figure S 5: Barplot of enriched gene sets according to fgsea comparing wild type and *canal-1* stems.

Supplemental Figure S 6: Phylogenetic tree of glutaredoxin genes orthogroup assembled by ggtree.

Supplemental Figure S 7: Expression of photosystem I reaction center genes. Supplemental Figure S 8: Expression of photosystem II oxygen evolving center genes. Supplemental Figure S 9: Principal component analysis of proteomics data.

Supplemental Figure S 10: Heatmap of raw unfiltered correlation values of photosystem components and assembly factor transcripts and proteins.

Supplemental Figure S 11: Whole-plant traits of different genotypes in experiment including overexpression lines.

Supplemental Figure S 12: Levenès-transformed level of harvest index (HI) in M82 wild type plants and ILs, from the seasons 2001 and 2004.

Supplemental Figure S 13: Alignment of SCO2 coding DNA sequence of *Solanum lycopersicum* (CDS_SL4.0) and *Solanum pennellii* (CDS_Spenn).

Supplemental Figure S 14: Alignment of SCO2 amino acid sequence of *Solanum lycopersicum* (AA_SL4.0) and *Solanum pennellii* (AA_Spenn).

Supplemental Data S 1: List of metabolic features measured across all platforms.

Supplemental Data S 2: Metabolite levels of all individual samples relative to respective wild type samples.

Supplemental Data S 3: QTL mapping using *S.pennellii* ILs for canalization of yield parameters using Levene’s transformed values considering GxE-effects.

Supplemental Data S 4: P-values of QTL mapping using *S.pennellii* ILs for canalization of yield parameters using Levene’s transformed values considering only genotype effects.

Supplemental Data S 5: Expression matrix of raw sequencing reads as produced by LSTrAP

Supplemental Data S 6: Expression matrix of deseq2-normalized reads and log2-fold changes and p-values of respective contrasts.

Supplemental Data S 7: Correspondence table between Solgenomics and Entrez Gene IDs Supplemental Data S 8: Table of protein Groups as produced by MaxQuant software.

Supplemental Data S 9: Table of quantile-normalized LFQ values of protein abundance and log2-fold changes and p-values of respective contrasts.

Supplemental Data S 10: EC number, enzyme name, symbol, KO and corresponding KEGG maps used in SGBVview figures.

Supplemental Data S 11: KEGG Compound ID, compound name and symbol used in SBGNview figures.

## Author Contributions

MWA, SA and ARF planned the work. JF and DZ supplied plant material and seeds from the *canal-1* mutant. MWA performed the experimental work and computational analysis. ES performed proteomics analysis. RL and NI advised the transcriptomics analysis. MWA, SA and ARF wrote and edited the manuscript.

