## supplementary files and figures for "Characterization of tomato *canal-1* mutant using a multi-omics approach"

Supplemental Table S 1: Enriched terms in GO enrichment analysis of transcriptome with topGO comparing wild type green leaves versus white *canal-1* leaves. Columns show ontology (BP: biological process, MF: molecular function, CC: cellular component), ID (Ontology identifier), Term (Ontology term name), (adjusted p-value), count (total number of enriched genes), up (number of upregulated genes), down (number of downregulated genes, Z-score (z-score as calculated in Material and Methods), -log10(p) (negative log10 of p-value) and Contrast (compared tissues and genotypes).

| Ontology | ID | Term | adj_pval | count | up | down | Z-score | -log10(p) | Contrast |
|---|---|---|---|---|---|---|---|---|---|
| BP | GO:0007018 | microtubule-based movement | 0.00034 | 30 | 29 | 1 | 5.1120772 | 3.46852108 | WTglB_vs_MUwl |
| BP | GO:0006629 | lipid metabolic process | 0.0065 | 94 | 25 | 69 | -4.53825348 | 2.18708664 | WTglB_vs_MUwl |
| BP | GO:0006950 | response to stress | 0.01 | 95 | 38 | 57 | -1.94935887 | 2 | WTglB_vs_MUwl |
| BP | GO:0015979 | photosynthesis | 0.023 | 26 | 0 | 26 | -5.09901951 | 1.63827216 | WTglB_vs_MUwl |
| BP | GO:0048544 | recognition of pollen | 0.023 | 24 | 3 | 21 | -3.67423461 | 1.63827216 | WTglB_vs_MUwl |
| MF | GO:0003777 | microtubule motor activity | 0.00012 | 30 | 29 | 1 | 5.1120772 | 3.92081875 | WTglB_vs_MUwl |
| MF | GO:0003700 | DNA-binding transcription factor activit... | 0.00021 | 130 | 33 | 97 | -5.61317132 | 3.67778071 | WTglB_vs_MUwl |
| MF | GO:0015035 | protein disulfide oxidoreductase activit... | 0.00031667 | 30 | 18 | 12 | 1.09544512 | 3.49939765 | WTglB_vs_MUwl |
| MF | GO:0008017 | microtubule binding | 0.000525 | 34 | 33 | 1 | 5.48795472 | 3.2798407 | WTglB_vs_MUwl |
| MF | GO:0043565 | sequence-specific DNA binding | 0.00148 | 82 | 23 | 59 | -3.97553494 | 2.82973828 | WTglB_vs_MUwl |
| MF | GO:0005215 | transporter activity | 0.00283333 | 161 | 44 | 117 | -5.75320597 | 2.54770233 | WTglB_vs_MUwl |
| MF | GO:0005509 | calcium ion binding | 0.00514286 | 67 | 7 | 60 | -6.47498055 | 2.28879554 | WTglB_vs_MUwl |
| MF | GO:0016597 | amino acid binding | 0.005875 | 16 | 9 | 7 | 0.5 | 2.23099213 | WTglB_vs_MUwl |
| MF | GO:0015297 | antiporter activity | 0.01022222 | 29 | 12 | 17 | -0.92847669 | 1.99045468 | WTglB_vs_MUwl |
| MF | GO:0004674 | protein serine/threonine kinase activity | 0.0246 | 27 | 2 | 25 | -4.42635206 | 1.60906489 | WTglB_vs_MUwl |
| MF | GO:0071949 | FAD binding | 0.02541667 | 7 | 2 | 5 | -1.13389342 | 1.59488141 | WTglB_vs_MUwl |
| MF | GO:0005524 | ATP binding | 0.02541667 | 419 | 194 | 225 | -1.5144491 | 1.59488141 | WTglB_vs_MUwl |
| MF | GO:0016762 | xyloglucan:xyloglucosyl transferase acti... | 0.03886667 | 13 | 0 | 13 | -3.60555128 | 1.4104227 | WTglB_vs_MUwl |
| MF | GO:0004197 | cysteine-type endopeptidase activity | 0.03886667 | 7 | 0 | 7 | -2.64575131 | 1.4104227 | WTglB_vs_MUwl |
| MF | GO:0008081 | phosphoric diester hydrolase activity | 0.03886667 | 15 | 4 | 11 | -1.80739223 | 1.4104227 | WTglB_vs_MUwl |
| MF | GO:1901505 | carbohydrate derivative transmembrane tr... | 0.04922222 | 6 | 2 | 4 | -0.81649658 | 1.30783878 | WTglB_vs_MUwl |
| MF | GO:0015932 | nucleobase-containing compound transmemb... | 0.04922222 | 6 | 2 | 4 | -0.81649658 | 1.30783878 | WTglB_vs_MUwl |
| MF | GO:0016758 | transferase activity, transferring hexos... | 0.04922222 | 91 | 17 | 74 | -5.97522357 | 1.30783878 | WTglB_vs_MUwl |
| CC | GO:0009538 | photosystem I reaction center | 0.000053 | 10 | 0 | 10 | -3.16227766 | 4.27572413 | WTglB_vs_MUwl |
| CC | GO:0016020 | membrane | 0.00385 | 368 | 93 | 275 | -9.48740584 | 2.41453927 | WTglB_vs_MUwl |
| CC | GO:0005618 | cell wall | 0.014 | 26 | 4 | 22 | -3.53009043 | 1.85387196 | WTglB_vs_MUwl |

Supplemental Table S 2: Enriched terms in GO enrichment analysis of transcriptome with topGO comparing green *canal-1* leaves versus white *canal-1* leaves. Columns show ontology (BP: biological process, MF: molecular function, CC: cellular component), ID (Ontology identifier), Term (Ontology term name), (adjusted p-value), count (total number of enriched genes), up (number of upregulated genes), down (number of downregulated genes, Z-score (z-score as calculated in Material and Methods), -log10(p) (negative log10 of p-value) and Contrast (compared tissues and genotypes).

| Ontology | ID | Term | adj_pval | count | up | down | Z-score | -log10(p) | Contrast |
|---|---|---|---|---|---|---|---|---|---|
| BP | GO:0015979 | photosynthesis | 3.9E-22 | 43 | 0 | 43 | -6.55743852 | 21.4089354 | MUglB_vs_MUwl |
| BP | GO:0045454 | cell redox homeostasis | 0.0019 | 30 | 10 | 20 | -1.82574186 | 2.7212464 | MUglB_vs_MUwl |
| BP | GO:0006662 | glycerol ether metabolic process | 0.00933333 | 6 | 0 | 6 | -2.44948974 | 2.02996322 | MUglB_vs_MUwl |
| MF | GO:0015035 | protein disulfide oxidoreductase activit... | 0.00013 | 21 | 9 | 12 | -0.65465367 | 3.88605665 | MUglB_vs_MUwl |
| MF | GO:0009055 | electron transfer activity | 0.00013 | 37 | 11 | 26 | -2.46598481 | 3.88605665 | MUglB_vs_MUwl |
| MF | GO:0016655 | oxidoreductase activity, acting on NAD(P... | 0.00366667 | 5 | 0 | 5 | -2.23606798 | 2.43572857 | MUglB_vs_MUwl |
| MF | GO:0003777 | microtubule motor activity | 0.02475 | 15 | 15 | 0 | 3.87298335 | 1.6064248 | MUglB_vs_MUwl |
| MF | GO:0051537 | 2 iron, 2 sulfur cluster binding | 0.0362 | 9 | 1 | 8 | -2.33333333 | 1.44129143 | MUglB_vs_MUwl |
| CC | GO:0009654 | photosystem II oxygen evolving complex | 1.2E-12 | 16 | 0 | 16 | -4 | 11.9208188 | MUglB_vs_MUwl |
| CC | GO:0019898 | extrinsic component of membrane | 3.5E-09 | 14 | 0 | 14 | -3.74165739 | 8.45593196 | MUglB_vs_MUwl |
| CC | GO:0009538 | photosystem I reaction center | 3.3333E-08 | 10 | 0 | 10 | -3.16227766 | 7.47712125 | MUglB_vs_MUwl |

Supplemental Table S 3: Enriched terms in GO enrichment analysis of transcriptome with topGO comparing wild type green leaves versus green *canal-1* leaves. Columns show ontology (BP: biological process, MF: molecular function, CC: cellular component), ID (Ontology identifier), Term (Ontology term name), (adjusted p-value), count (total number of enriched genes), up (number of upregulated genes), down (number of downregulated genes, Z-score (z-score as calculated in Material and Methods), -log10(p) (negative log10 of p-value) and Contrast (compared tissues and genotypes).

| Ontology | ID | Term | adj_pval | count | up | down | Z-score | -log10(p) | Contrast |
|---|---|---|---|---|---|---|---|---|---|
| BP | GO:0006418 | tRNA aminoacylation for protein translat... | 0.000059 | 15 | 14 | 1 | 3.35658557 | 4.22914799 | WTglB_vs_MUgl |
| BP | GO:0006629 | lipid metabolic process | 0.0075 | 41 | 19 | 22 | -0.46852129 | 2.12493874 | WTglB_vs_MUgl |
| BP | GO:0045454 | cell redox homeostasis | 0.008 | 21 | 15 | 6 | 1.96396101 | 2.09691001 | WTglB_vs_MUgl |
| MF | GO:0004812 | aminoacyl-tRNA ligase activity | 0.000018 | 15 | 14 | 1 | 3.35658557 | 4.74472749 | WTglB_vs_MUgl |
| MF | GO:0005509 | calcium ion binding | 0.0085 | 29 | 8 | 21 | -2.4140394 | 2.07058107 | WTglB_vs_MUgl |
| MF | GO:0015035 | protein disulfide oxidoreductase activit... | 0.01066667 | 13 | 8 | 5 | 0.83205029 | 1.97197128 | WTglB_vs_MUgl |
| MF | GO:0004867 | serine-type endopeptidase inhibitor acti... | 0.02875 | 7 | 1 | 6 | -1.88982237 | 1.54136215 | WTglB_vs_MUgl |
| MF | GO:0003700 | DNA-binding transcription factor activit... | 0.042 | 44 | 4 | 40 | -5.4272042 | 1.37675071 | WTglB_vs_MUgl |
| MF | GO:0004866 | endopeptidase inhibitor activity | 0.05 | 13 | 1 | 12 | -3.05085108 | 1.30103 | WTglB_vs_MUgl |
| CC | GO:0005737 | cytoplasm | 0.00098 | 54 | 36 | 18 | 2.44948974 | 3.00877392 | WTglB_vs_MUgl |
| CC | GO:0009654 | photosystem II oxygen evolving complex | 0.00155 | 7 | 7 | 0 | 2.64575131 | 2.8096683 | WTglB_vs_MUgl |
| CC | GO:0019898 | extrinsic component of membrane | 0.01466667 | 6 | 6 | 0 | 2.44948974 | 1.83366858 | WTglB_vs_MUgl |
| CC | GO:0009507 | chloroplast | 0.03375 | 6 | 5 | 1 | 1.63299316 | 1.47172622 | WTglB_vs_MUgl |

Supplemental Table S 4: Enriched terms in GO enrichment analysis of transcriptome with topGO comparing wild type stems versus *canal-1* stems. Columns show ontology (BP: biological process, MF: molecular function, CC: cellular component), ID (Ontology identifier), Term (Ontology term name), (adjusted p-value), count (total number of enriched genes), up (number of upregulated genes), down (number of downregulated genes, Z-score (z-score as calculated in Material and Methods), -log10(p) (negative log10 of p-value) and Contrast (compared tissues and genotypes).

| Ontology | ID | Term | adj_pval | count | up | down | Z-score | -log10(p) | Contrast |
|---|---|---|---|---|---|---|---|---|---|
| BP | GO:0046274 | lignin catabolic process | 0.0048 | 6 | 6 | 0 | 2.44948974 | 2.31875876 | WTsB_vs_MUs |
| BP | GO:0006355 | regulation of transcription, DNA-templat... | 0.01 | 44 | 11 | 33 | -3.31662479 | 2 | WTsB_vs_MUs |
| BP | GO:0042546 | cell wall biogenesis | 0.04333333 | 8 | 3 | 5 | -0.70710678 | 1.3631779 | WTsB_vs_MUs |
| BP | GO:0010411 | xyloglucan metabolic process | 0.045 | 5 | 1 | 4 | -1.34164079 | 1.34678749 | WTsB_vs_MUs |
| MF | GO:0005507 | copper ion binding | 0.000245 | 11 | 8 | 3 | 1.50755672 | 3.61083392 | WTsB_vs_MUs |
| MF | GO:0003700 | DNA-binding transcription factor activit... | 0.000245 | 33 | 6 | 27 | -3.65563078 | 3.61083392 | WTsB_vs_MUs |
| MF | GO:0004857 | enzyme inhibitor activity | 0.0004 | 15 | 6 | 9 | -0.77459667 | 3.39794001 | WTsB_vs_MUs |
| MF | GO:0052716 | hydroquinone:oxygen oxidoreductase activ... | 0.0011 | 6 | 6 | 0 | 2.44948974 | 2.95860731 | WTsB_vs_MUs |
| MF | GO:0016491 | oxidoreductase activity | 0.003 | 77 | 49 | 28 | 2.39317211 | 2.52287875 | WTsB_vs_MUs |
| MF | GO:0016760 | cellulose synthase (UDP-forming) activit... | 0.02333333 | 5 | 4 | 1 | 1.34164079 | 1.63202321 | WTsB_vs_MUs |
| MF | GO:0016758 | transferase activity, transferring hexos... | 0.02442857 | 27 | 16 | 11 | 0.96225045 | 1.61210193 | WTsB_vs_MUs |
| MF | GO:0051082 | unfolded protein binding | 0.03033333 | 8 | 7 | 1 | 2.12132034 | 1.51807986 | WTsB_vs_MUs |
| MF | GO:0016762 | xyloglucan:xyloglucosyl transferase acti... | 0.03033333 | 5 | 1 | 4 | -1.34164079 | 1.51807986 | WTsB_vs_MUs |
| CC | GO:0048046 | apoplast | 0.0000022 | 11 | 7 | 4 | 0.90453403 | 5.65757732 | WTsB_vs_MUs |
| CC | GO:0005618 | cell wall | 0.000365 | 10 | 2 | 8 | -1.8973666 | 3.43770714 | WTsB_vs_MUs |

Supplemental Table S 5: List of tomato genes orthologous to known photosynthesis genes in *Arabidopsis thaliana*. #: Number of orthologues found in *S. lycopersicum*.

| Gene name | A. thaliana locus | Photosystem | Group | Orthogroup | S. lycopersicum gene | # | Unique name |
|---|---|---|---|---|---|---|---|
| cpSRP43 | AT2G47450 | II | assembly | OG0010526 | Solyc02g087400.1 | 1 | cpSRP43 |
| cpSRP54 | AT5G03940 | II | assembly | OG0007531 | Solyc09g009940.3 | 1 | cpSRP54 |
| cpFtsY | AT2G45770 | II | assembly | OG0010619 | Solyc01g091580.3 | 1 | cpFtsY |
| ALB3 | AT2G28800 | II | assembly | OG0008270 | Solyc11g066110.2 | 1 | ALB3 |
| cpSecA1 | AT4G01800 | II | assembly | OG0008510 | Solyc01g080840.3 | 1 | cpSecA1 |
| cpSecA2 | AT1G21650 | II | assembly | OG0011249 | Solyc11g005020.3 | 1 | cpSecA2 |
| cpSecE2 | AT4G38490 | II | assembly | OG0011081 | Solyc05g050720.4 | 1 | cpSecE2 |
| cpSecY1 | AT2G18710 | II | assembly | OG0010623 | Solyc07g006520.3 | 1 | cpSecY1 |
| cpSecY2 | AT2G31530 | II | assembly | OG0010484 | Solyc09g065120.4 | 1 | cpSecY2 |
| Tha4 | AT5G28750 | II | assembly | OG0009524 | Solyc01g097310.3 | 1 | Tha4 |
| HCF106 | AT5G52440 | II | assembly | OG0007686 | Solyc03g025310.3 | 1 | HCF106 |
| cpTatC | AT2G01110 | II | assembly | OG0008157 | Solyc05g008530.3 | 1 | cpTatC |
| CPRabA5e | AT1G05810 | II | assembly | OG0001232 | Solyc09g056340.3 | 4 | CPRabA5e#1 |
| CPRabA5e | AT1G05810 | II | assembly | OG0001232 | Solyc09g097900.3 | 4 | CPRabA5e#2 |
| CPRabA5e | AT1G05810 | II | assembly | OG0001232 | Solyc09g098170.4 | 4 | CPRabA5e#3 |
| CPRabA5e | AT1G05810 | II | assembly | OG0001232 | Solyc11g012460.3 | 4 | CPRabA5e#4 |
| CYO1/SCO2 | AT3G19220 | II | assembly | OG0013960 | Solyc01g108200.3 | 1 | CYO1/SCO2 |
| THF1/PSB29 | AT2G20890 | II | assembly | OG0006507 | Solyc07g054820.4 | 1 | THF1/PSB29 |
| TerC | AT5G12130 | II | assembly | OG0009789 | Solyc10g085980.2 | 1 | TerC |
| VIPP1 | AT1G65260 | II | assembly | OG0006927 | Solyc11g008990.2 | 1 | VIPP1 |
| PPL1 | AT3G55330 | II | assembly | OG0009329 | Solyc03g114930.3 | 1 | PPL1 |
| ELIP1 | AT3G22840 | II | assembly | OG0002986 | Solyc09g082690.3 | 2 | ELIP1#1 |
| ELIP1 | AT3G22840 | II | assembly | OG0002986 | Solyc09g082700.2 | 2 | ELIP1#2 |
| SEP3.1/LIL3:1 | AT4G17600 | II | assembly | OG0004419 | Solyc08g007180.3 | 2 | SEP3.1/LIL3:1#1 |
| SEP3.1/LIL3:1 | AT4G17600 | II | assembly | OG0004419 | Solyc08g077880.3 | 2 | SEP3.1/LIL3:1#2 |
| HCF173 | AT1G16720 | II | assembly | OG0008936 | Solyc08g016080.3 | 1 | HCF173 |
| HCF244 | AT4G35250 | II | assembly | OG0011110 | Solyc01g112060.4 | 1 | HCF244 |
| CtpA1 | AT3G57680 | II | assembly | OG0012163 | Solyc03g059260.4 | 1 | CtpA1 |
| CtpA2 | AT4G17740 | II | assembly | OG0010986 | Solyc12g097030.2 | 1 | CtpA2 |
| LPA1/PratA | AT1G02910 | II | assembly | OG0011435 | Solyc09g074880.4 | 1 | LPA1/PratA |
| MET1 | AT1G55480 | II | assembly | OG0011540 | Solyc01g005520.4 | 1 | MET1 |
| LQY1 | AT1G75690 | II | assembly | OG0011672 | Solyc04g081320.4 | 1 | LQY1 |
| PDI6/PDIL1-2 | AT1G77510 | II | assembly | OG0001320 | Solyc05g056400.3 | 4 | PDI6/PDIL1-2#1 |
| PDI6/PDIL1-2 | AT1G77510 | II | assembly | OG0001320 | Solyc06g005940.3 | 4 | PDI6/PDIL1-2#2 |
| PDI6/PDIL1-2 | AT1G77510 | II | assembly | OG0001320 | Solyc06g060290.4 | 4 | PDI6/PDIL1-2#3 |
| PDI6/PDIL1-2 | AT1G77510 | II | assembly | OG0001320 | Solyc07g021490.3 | 4 | PDI6/PDIL1-2#4 |
| TRX-M1 | AT1G03680 | II | assembly | OG0000839 | Solyc07g063190.3 | 4 | TRX-M1#1 |
| TRX-M1 | AT1G03680 | II | assembly | OG0000839 | Solyc10g006970.3 | 4 | TRX-M1#2 |
| TRX-M1 | AT1G03680 | II | assembly | OG0000839 | Solyc10g008390.3 | 4 | TRX-M1#3 |
| TRX-M1 | AT1G03680 | II | assembly | OG0000839 | Solyc12g013810.2 | 4 | TRX-M1#4 |
| LTO1 | AT4G35760 | II | assembly | OG0008492 | Solyc02g083270.4 | 1 | LTO1 |
| RBD1 | AT1G54500 | II | assembly | OG0004673 | Solyc01g097910.2 | 1 | RBD1 |
| CYP38/TLP40 | AT3G01480 | II | assembly | OG0011891 | Solyc02g086910.4 | 1 | CYP38/TLP40 |
| FKBP20-2 | AT3G60370 | II | assembly | OG0012045 | Solyc10g039270.2 | 1 | FKBP20-2 |
| STN7 | AT1G68830 | II | assembly | OG0011625 | Solyc12g021280.2 | 1 | STN7 |
| STN8 | AT5G01920 | II | assembly | OG0009845 | Solyc10g085680.2 | 1 | STN8 |
| PBCP | AT2G30170 | II | assembly | OG0010406 | Solyc06g007350.4 | 1 | PBCP |
| TLP18.3 | AT1G54780 | II | assembly | OG0011387 | Solyc01g098640.3 | 1 | TLP18.3 |
| PPH1/TAP38 | AT4G27800 | II | assembly | OG0010746 | Solyc03g082960.2 | 1 | PPH1/TAP38 |
| FtsH1 | AT1G50250 | II | assembly | OG0008832 | Solyc04g082250.3 | 1 | FtsH1 |
| FtsH2/VAR2 | AT2G30950 | II | assembly | OG0002445 | Solyc02g081550.3 | 2 | FtsH2/VAR2#1 |
| FtsH2/VAR2 | AT2G30950 | II | assembly | OG0002445 | Solyc07g055320.4 | 2 | FtsH2/VAR2#2 |
| FtsH11 | AT5G53170 | II | assembly | OG0009692 | Solyc03g007330.3 | 1 | FtsH11 |
| Deg1 | AT3G27925 | II | assembly | OG0007223 | Solyc02g086830.3 | 1 | Deg1 |
| Deg2 | AT2G47940 | II | assembly | OG0010709 | Solyc09g074420.4 | 1 | Deg2 |
| Deg5 | AT4G18370 | II | assembly | OG0010973 | Solyc08g048550.3 | 1 | Deg5 |
| Deg7 | AT3G03380 | II | assembly | OG0009303 | Solyc02g091410.3 | 2 | Deg7#1 |
| Deg7 | AT3G03380 | II | assembly | OG0009303 | Solyc03g043660.3 | 2 | Deg7#2 |
| Deg8 | AT5G39830 | II | assembly | OG0012317 | Solyc02g067360.3 | 1 | Deg8 |
| HCF243 | AT3G15095 | II | assembly | OG0005901 | Solyc07g062730.1 | 2 | HCF243#1 |
| HCF243 | AT3G15095 | II | assembly | OG0005901 | Solyc12g014040.1 | 2 | HCF243#2 |
| PSB27-H1 | AT1G03600 | II | assembly | OG0009083 | Solyc07g054290.1 | 1 | PSB27-H1 |
| PSB27-H2/LPA19 | AT1G05385 | II | assembly | OG0013785 | Solyc09g076030.4 | 1 | PSB27-H2/LPA19 |
| HCF136 | AT5G23120 | II | assembly | OG0014076 | Solyc02g014150.4 | 1 | HCF136 |
| PAM68 | AT4G19100 | II | assembly | OG0010885 | Solyc12g096100.2 | 1 | PAM68 |
| PSB28 | AT4G28660 | II | assembly | OG0004386 | Solyc09g064500.3 | 1 | PSB28 |
| LPA2 | AT5G51545 | II | assembly | OG0012506 | Solyc03g083570.3 | 1 | LPA2 |
| LPA3 | AT1G73060 | II | assembly | OG0011396 | Solyc06g068480.3 | 1 | LPA3 |
| PSB33 | AT1G71500 | II | assembly | OG0011429 | Solyc11g042640.2 | 1 | PSB33 |
| HHL1 | AT1G67700 | II | assembly | OG0011291 | Solyc05g014310.3 | 1 | HHL1 |
| MPH1 | AT5G07020 | II | assembly | OG0009484 | Solyc01g096660.3 | 1 | MPH1 |
| Ycf3 | ATCG00360 | I | assembly | OG0003400 | Solyc00g500058.1 | 3 | Ycf3#1 |
| Ycf3 | ATCG00360 | I | assembly | OG0003400 | Solyc00g500145.1 | 3 | Ycf3#2 |

| | | | | | | | |
|---|---|---|---|---|---|---|---|
| Ycf3 | ATCG00360 | I | assembly | OG0003400 | Solyc00g500210.1 | 3 | Ycf3#3 |
| Ycf4 | ATCG00520 | I | assembly | OG0007956 | Solyc00g500019.1 | 2 | Ycf4#1 |
| Ycf4 | ATCG00520 | I | assembly | OG0007956 | Solyc00g500066.1 | 2 | Ycf4#2 |
| Ycf37/Pyg7 | AT1G22700 | I | assembly | OG0007009 | Solyc11g017200.2 | 1 | Ycf37/Pyg7 |
| Y3IP1 | AT5G44650 | I | assembly | OG0009574 | Solyc10g005180.3 | 1 | Y3IP1 |
| PPD1 | AT4G15510 | I | assembly | OG0008344 | Solyc01g106090.3 | 1 | PPD1 |
| Psa2 | AT2G34860 | I | assembly | OG0010679 | Solyc01g006130.4 | 1 | Psa2 |
| Hcf101 | AT3G24430 | I | assembly | OG0012190 | Solyc12g007010.2 | 1 | Hcf101 |
| CnfU | AT4G01940 | I | assembly | OG0011048 | Solyc01g079220.4 | 2 | CnfU#1 |
| CnfU | AT5G49940 | I | assembly | OG0009556 | Solyc01g103710.4 | 2 | CnfU#2 |
| PsaA | ATCG00350 | I | component | OG0000798 | Solyc00g500057.1 | 6 | PsaA#1 |
| PsaA | ATCG00350 | I | component | OG0000798 | Solyc00g500139.1 | 6 | PsaA#2 |
| PsaA | ATCG00350 | I | component | OG0000798 | Solyc00g500209.1 | 6 | PsaA#3 |
| PsaA | ATCG00350 | I | component | OG0000798 | Solyc02g020960.2 | 6 | PsaA#4 |
| PsaA | ATCG00350 | I | component | OG0000798 | Solyc10g017890.1 | 6 | PsaA#5 |
| PsaA | ATCG00350 | I | component | OG0000798 | Solyc12g033040.1 | 6 | PsaA#6 |
| PsaB | ATCG00340 | I | component | OG0006338 | Solyc00g500056.1 | 3 | PsaB#1 |
| PsaB | ATCG00340 | I | component | OG0006338 | Solyc00g500138.1 | 3 | PsaB#2 |
| PsaB | ATCG00340 | I | component | OG0006338 | Solyc00g500208.1 | 3 | PsaB#3 |
| PsaC | ATCG01060 | I | component | OG0010316 | Solyc02g011760.1 | 1 | PsaC |
| PsaD | AT4G02770 | I | component | OG0006784 | Solyc06g054260.1 | 1 | PsaD |
| PsaE | AT4G28750 | I | component | OG0004265 | Solyc06g083680.3 | 2 | PsaE#1 |
| PsaE | AT4G28750 | I | component | OG0004265 | Solyc09g063130.3 | 2 | PsaE#2 |
| PsaF | AT1G31330 | I | component | OG0005761 | Solyc02g069450.3 | 3 | PsaF#1 |
| PsaF | AT1G31330 | I | component | OG0005761 | Solyc02g069460.3 | 3 | PsaF#2 |
| PsaF | AT1G31330 | I | component | OG0005761 | Solyc08g074307.2 | 3 | PsaF#3 |
| PsaG | AT1G55670 | I | component | OG0009104 | Solyc07g066150.1 | 1 | PsaG |
| PsaH | AT3G16140 | I | component | OG0003807 | Solyc03g120640.3 | 3 | PsaH#1 |
| PsaH | AT3G16140 | I | component | OG0003807 | Solyc06g066640.3 | 3 | PsaH#2 |
| PsaH | AT3G16140 | I | component | OG0003807 | Solyc12g044280.2 | 3 | PsaH#3 |
| PsaI | ATCG00510 | I | component | OG0020062 | Solyc07g024040.2 | 1 | PsaI |
| PsaJ | ATCG00630 | I | component | OG0017056 | Solyc09g064400.2 | 1 | PsaJ |
| PsaK | AT1G30380 | I | component | OG0011531 | Solyc08g006930.3 | 1 | PsaK |
| PsaL | AT4G12800 | I | component | OG0005378 | Solyc06g082940.3 | 2 | PsaL#1 |
| PsaL | AT4G12800 | I | component | OG0005378 | Solyc06g082950.4 | 2 | PsaL#2 |
| PsaN | AT5G64040 | I | component | OG0007548 | Solyc08g013670.3 | 1 | PsaN |
| PsaO | AT1G08380 | I | component | OG0011194 | Solyc06g074200.4 | 1 | PsaO |
| PsaP | AT2G46820 | I | component | OG0006416 | Solyc10g005050.3 | 1 | PsaP |
| Lhca1 | AT3G54890 | I | component | OG0009336 | Solyc05g056050.3 | 1 | Lhca1 |
| Lhca2 | AT3G61470 | I | component | OG0009183 | Solyc10g006230.3 | 1 | Lhca2 |
| Lhca3 | AT1G61520 | I | component | OG0006983 | Solyc10g007690.3 | 2 | Lhca3#1 |
| Lhca3 | AT1G61520 | I | component | OG0006983 | Solyc12g011280.2 | 2 | Lhca3#2 |
| Lhca4 | AT3G47470 | I | component | OG0005886 | Solyc06g069730.3 | 1 | Lhca4 |
| Lhca5 | AT1G45474 | I | component | OG0011223 | Solyc07g022900.4 | 1 | Lhca5 |
| Lhca6 | AT1G19150 | I | component | OG0011568 | Solyc12g009200.2 | 1 | Lhca6 |
| psbA | ATCG00020 | II | component | OG0001627 | Solyc00g500130.1 | 4 | psbA#1 |
| psbA | ATCG00020 | II | component | OG0001627 | Solyc00g500200.1 | 4 | psbA#2 |
| psbA | ATCG00020 | II | component | OG0001627 | Solyc00g500296.1 | 4 | psbA#3 |
| psbA | ATCG00020 | II | component | OG0001627 | Solyc00g500329.1 | 4 | psbA#4 |
| psbD | ATCG00270 | II | component | OG0001951 | Solyc00g230080.1 | 5 | psbD#1 |
| psbD | ATCG00270 | II | component | OG0001951 | Solyc05g021190.2 | 5 | psbD#2 |
| psbD | ATCG00270 | II | component | OG0001951 | Solyc07g008990.1 | 5 | psbD#3 |
| psbD | ATCG00270 | II | component | OG0001951 | Solyc08g065370.1 | 5 | psbD#4 |
| psbD | ATCG00270 | II | component | OG0001951 | Solyc09g055950.2 | 5 | psbD#5 |
| psbE | ATCG00580 | II | component | OG0007958 | Solyc00g500021.1 | 2 | psbE#1 |
| psbE | ATCG00580 | II | component | OG0007958 | Solyc00g500068.1 | 2 | psbE#2 |
| psb1 | ATCG00080 | II | component | OG0002839 | Solyc00g500049.1 | 5 | psb1#1 |
| psb1 | ATCG00080 | II | component | OG0002839 | Solyc00g500132.1 | 5 | psb1#2 |
| psb1 | ATCG00080 | II | component | OG0002839 | Solyc00g500202.1 | 5 | psb1#3 |
| psb1 | ATCG00080 | II | component | OG0002839 | Solyc00g500293.1 | 5 | psb1#4 |
| psb1 | ATCG00080 | II | component | OG0002839 | Solyc00g500331.1 | 5 | psb1#5 |
| psbB | ATCG00680 | II | component | OG0003399 | Solyc00g500024.1 | 2 | psbB#1 |
| psbB | ATCG00680 | II | component | OG0003399 | Solyc00g500071.1 | 2 | psbB#2 |

Supplemental Table S 6: Enriched terms in gene set enrichment analysis of transcriptome with fgsea comparing wild type green leaves versus white *canal-1* leaves. Columns show ontology (BP: biological process, MF: molecular function, CC: cellular component), ID (Ontology identifier), Term (Ontology term name), (adjusted p-value), count (total number of enriched genes), up (number of upregulated genes), down (number of downregulated genes, Z-score (z-score as calculated in Material and Methods), -log10(p) (negative log10 of p-value) and Contrast (compared tissues and genotypes).

| pathway | pval | padj | log2err | ES | NES | size | leadingEdge | Contrast |
|---|---|---|---|---|---|---|---|---|
| pseudouridine synthesis | 0.00293541 | 0.0219383 | 0.4317077 | 0.77530245 | 1.75366964 | 22 | | WTglB_vs_MUwl |
| amino acid transmembrane transport | 0.00543486 | 0.02964016 | 0.40701792 | -0.75831453 | -1.65194056 | 20 | | WTglB_vs_MUwl |

| | | | | | | | | |
|---|---|---|---|---|---|---|---|---|
| DNA metabolic process | 0.00349539 | 0.02363552 | 0.4317077 | 0.86530633 | 1.73437831 | 11 | | WTglB_vs_MUwl |
| DNA replication | 0.00026006 | 0.00307742 | 0.49849311 | 0.71357865 | 1.85967126 | 47 | | WTglB_vs_MUwl |
| DNA topological change | 0.0037949 | 0.02437703 | 0.4317077 | 0.88379804 | 1.72400511 | 10 | | WTglB_vs_MUwl |
| DNA repair | 1.424E-06 | 4.5974E-05 | 0.64355184 | 0.71188011 | 2.03658621 | 79 | | WTglB_vs_MUwl |
| rRNA processing | 0.00047794 | 0.00484765 | 0.49849311 | 0.77591999 | 1.83664184 | 30 | | WTglB_vs_MUwl |
| RNA processing | 4.8504E-07 | 3.4438E-05 | 0.6594444 | 0.74711958 | 2.06544004 | 67 | | WTglB_vs_MUwl |
| translation | 4.0309E-35 | 5.7239E-33 | 1.54318309 | 0.77382971 | 2.62525813 | 312 | | WTglB_vs_MUwl |
| translational elongation | 0.0001177 | 0.00167129 | 0.5384341 | 0.81776887 | 1.89805669 | 25 | | WTglB_vs_MUwl |
| tRNA aminoacylation for protein translation | 1.9425E-06 | 4.5974E-05 | 0.62725674 | 0.79575127 | 2.01140002 | 41 | | WTglB_vs_MUwl |
| protein folding | 0.00013793 | 0.00178052 | 0.51884808 | 0.5755282 | 1.77239788 | 126 | | WTglB_vs_MUwl |
| protein phosphorylation | 1.6968E-06 | 4.5974E-05 | 0.64355184 | -0.42702232 | -1.48532943 | 897 | | WTglB_vs_MUwl |
| proteolysis | 0.00533656 | 0.02964016 | 0.40701792 | -0.42533034 | -1.36370835 | 291 | | WTglB_vs_MUwl |
| lipid metabolic process | 0.00117938 | 0.01116482 | 0.45505987 | -0.54369601 | -1.60585702 | 128 | | WTglB_vs_MUwl |
| ion transport | 0.00180435 | 0.01507162 | 0.45505987 | -0.73948249 | -1.74512198 | 31 | | WTglB_vs_MUwl |
| nucleocytoplasmic transport | 0.00046634 | 0.00484765 | 0.49849311 | -0.6933639 | -1.7984414 | 51 | | WTglB_vs_MUwl |
| defense response | 0.00438155 | 0.02592416 | 0.40701792 | -0.68252659 | -1.68719621 | 40 | | WTglB_vs_MUwl |
| microtubule-based movement | 1.1651E-06 | 4.5974E-05 | 0.64355184 | 0.76853114 | 2.10159428 | 60 | | WTglB_vs_MUwl |
| vacuolar transport | 0.00205655 | 0.01622388 | 0.4317077 | -0.8500796 | -1.68178474 | 13 | | WTglB_vs_MUwl |
| signal transduction | 0.00316865 | 0.02249744 | 0.4317077 | -0.48947223 | -1.49187743 | 164 | | WTglB_vs_MUwl |
| metabolic process | 7.7833E-05 | 0.00123855 | 0.5384341 | -0.43055034 | -1.46750185 | 644 | | WTglB_vs_MUwl |
| RNA modification | 0.00645764 | 0.03210635 | 0.40701792 | 0.77046957 | 1.74410133 | 21 | | WTglB_vs_MUwl |
| response to biotic stimulus | 0.00148259 | 0.01315798 | 0.45505987 | -0.78533983 | -1.74470186 | 22 | | WTglB_vs_MUwl |
| photosynthesis | 4.1113E-06 | 8.3402E-05 | 0.61052688 | -0.77913329 | -2.02090933 | 51 | | WTglB_vs_MUwl |
| metal ion transport | 0.00394839 | 0.02437703 | 0.40701792 | -0.59579098 | -1.61871611 | 71 | | WTglB_vs_MUwl |
| methylation | 0.00563581 | 0.02964016 | 0.40701792 | 0.79503961 | 1.69269744 | 16 | | WTglB_vs_MUwl |
| ribosome biogenesis | 7.85E-05 | 0.00123855 | 0.5384341 | 0.8635196 | 1.93617355 | 20 | | WTglB_vs_MUwl |
| recognition of pollen | 0.00655693 | 0.03210635 | 0.40701792 | -0.60921835 | -1.60623769 | 55 | | WTglB_vs_MUwl |

Supplemental Table S 7: Enriched terms in gene set enrichment analysis of transcriptome with fgsea comparing green *canal-1* leaves versus white *canal-1* leaves. Columns show ontology (BP: biological process, MF: molecular function, CC: cellular component), ID (Ontology identifier), Term (Ontology term name), (adjusted p-value), count (total number of enriched genes), up (number of upregulated genes), down (number of downregulated genes, Z-score (z-score as calculated in Material and Methods), -log10(p) (negative log10 of p-value) and Contrast (compared tissues and genotypes).

| pathway | pval | padj | log2err | ES | NES | size | leadingEdge | Contrast |
|---|---|---|---|---|---|---|---|---|
| carbohydrate metabolic process | 0.0004196 | 0.00788585 | 0.49849311 | -0.51375261 | -1.49059619 | 340 | | MUglB_vs_MUwl |
| glucose metabolic process | 0.00091541 | 0.01278664 | 0.47727082 | -0.88861741 | -1.68827034 | 12 | | MUglB_vs_MUwl |
| DNA replication | 0.00208642 | 0.01784757 | 0.4317077 | 0.67025895 | 1.73603649 | 47 | | MUglB_vs_MUwl |
| DNA repair | 1.7605E-06 | 7.8636E-05 | 0.64355184 | 0.73459686 | 2.08979223 | 78 | | MUglB_vs_MUwl |
| DNA recombination | 0.00280407 | 0.02210268 | 0.4317077 | 0.78384767 | 1.74926432 | 20 | | MUglB_vs_MUwl |
| RNA processing | 0.00110848 | 0.01278664 | 0.45505987 | 0.65849053 | 1.82103002 | 64 | | MUglB_vs_MUwl |
| translation | 1.1021E-05 | 0.0003692 | 0.59332548 | 0.47677908 | 1.61867622 | 306 | | MUglB_vs_MUwl |
| proteolysis | 7.5062E-05 | 0.00167639 | 0.5384341 | -0.55017608 | -1.57612523 | 283 | | MUglB_vs_MUwl |
| defense response | 0.0004708 | 0.00788585 | 0.49849311 | -0.78053508 | -1.75161975 | 36 | | MUglB_vs_MUwl |
| microtubule-based movement | 0.00106134 | 0.01278664 | 0.45505987 | 0.675998 | 1.84710892 | 57 | | MUglB_vs_MUwl |
| metabolic process | 3.4232E-09 | 2.2935E-07 | 0.77493903 | -0.54454576 | -1.6223729 | 624 | | MUglB_vs_MUwl |
| response to biotic stimulus | 3.9781E-05 | 0.00106613 | 0.55733224 | -0.8732571 | -1.80958544 | 20 | | MUglB_vs_MUwl |
| response to auxin | 0.00551946 | 0.03892669 | 0.40701792 | -0.62171807 | -1.55075896 | 75 | | MUglB_vs_MUwl |
| photosynthesis | 1.2505E-09 | 1.6757E-07 | 0.78818681 | -0.87006755 | -2.04035123 | 49 | | MUglB_vs_MUwl |
| ATP synthesis coupled proton transport | 0.00501337 | 0.03732172 | 0.40701792 | -0.7804845 | -1.63141143 | 23 | | MUglB_vs_MUwl |
| metal ion transport | 0.00213105 | 0.01784757 | 0.4317077 | -0.65130623 | -1.6021106 | 70 | | MUglB_vs_MUwl |
| ribosome biogenesis | 0.00159847 | 0.01529962 | 0.45505987 | 0.81630902 | 1.79888136 | 19 | | MUglB_vs_MUwl |
| cell redox homeostasis | 0.00124049 | 0.01278664 | 0.45505987 | -0.60640084 | -1.58842087 | 112 | | MUglB_vs_MUwl |
| transmembrane transport | 0.00122639 | 0.01278664 | 0.45505987 | -0.49045049 | -1.42764989 | 379 | | MUglB_vs_MUwl |

Supplemental Table S 8: Enriched terms in gene set enrichment analysis of transcriptome with fgsea comparing wild type green leaves versus green *canal-1* leaves. Columns show ontology (BP: biological process, MF: molecular function, CC: cellular component), ID (Ontology identifier), Term (Ontology term name), (adjusted p-value), count (total number of enriched genes), up (number of upregulated genes), down (number of downregulated genes, Z-score (z-score as calculated in Material and Methods), -log10(p) (negative log10 of p-value) and Contrast (compared tissues and genotypes).

| pathway | pval | padj | log2err | ES | NES | size | leadingEdge | Contrast |
|---|---|---|---|---|---|---|---|---|
| DNA replication | 0.00084664 | 0.01126028 | 0.47727082 | 0.68966147 | 1.82269383 | 47 | | WTglB_vs_MUgl |
| regulation of transcription, DNA-templated | 2.179E-10 | 1.449E-08 | 0.8266573 | -0.45240781 | -1.716752 | 614 | | WTglB_vs_MUgl |
| rRNA processing | 0.00359977 | 0.02992307 | 0.4317077 | 0.71744588 | 1.72631136 | 30 | | WTglB_vs_MUgl |
| translation | 3.7426E-36 | 4.9776E-34 | 1.56318198 | 0.77417915 | 2.53849654 | 300 | | WTglB_vs_MUgl |
| translational elongation | 0.00013499 | 0.0029923 | 0.51884808 | 0.83122733 | 1.93676735 | 25 | | WTglB_vs_MUgl |
| tRNA aminoacylation for protein translation | 2.1655E-08 | 9.6004E-07 | 0.73376199 | 0.8491644 | 2.19862676 | 41 | | WTglB_vs_MUgl |
| protein folding | 4.1254E-07 | 1.3717E-05 | 0.67496286 | 0.65511468 | 1.97530073 | 123 | | WTglB_vs_MUgl |
| protein phosphorylation | 4.7713E-06 | 0.00012692 | 0.61052688 | -0.36870213 | -1.43098834 | 826 | | WTglB_vs_MUgl |
| nucleocytoplasmic transport | 0.00050278 | 0.00742998 | 0.47727082 | -0.65062359 | -1.83122239 | 50 | | WTglB_vs_MUgl |
| response to stress | 0.00464163 | 0.0363139 | 0.40701792 | -0.55981883 | -1.61559498 | 59 | | WTglB_vs_MUgl |
| microtubule-based movement | 0.00139611 | 0.01428324 | 0.45505987 | 0.65382627 | 1.77302688 | 55 | | WTglB_vs_MUgl |
| small GTPase mediated signal transduction | 0.00131155 | 0.01428324 | 0.45505987 | -0.54666352 | -1.68669689 | 89 | | WTglB_vs_MUgl |
| tRNA processing | 0.00314239 | 0.0278625 | 0.4317077 | 0.73822309 | 1.73696258 | 27 | | WTglB_vs_MUgl |
| response to wounding | 0.00047411 | 0.00742998 | 0.49849311 | -0.90294155 | -1.81835453 | 11 | | WTglB_vs_MUgl |
| photosynthesis | 0.00107545 | 0.0130032 | 0.45505987 | 0.68416556 | 1.80816879 | 47 | | WTglB_vs_MUgl |
| protein ubiquitination | 0.00023368 | 0.00443984 | 0.51884808 | -0.6071106 | -1.82899346 | 72 | | WTglB_vs_MUgl |
| ribosome biogenesis | 0.00164827 | 0.01565859 | 0.45505987 | 0.82238939 | 1.79762187 | 19 | | WTglB_vs_MUgl |

Supplemental Table S 9: Enriched terms in gene set enrichment analysis of transcriptome with fgsea comparing wild type stems versus *canal-1* stems. Columns show ontology (BP: biological process, MF: molecular function, CC: cellular component), ID (Ontology identifier), Term (Ontology term name), (adjusted p-value), count (total number of enriched genes), up (number of upregulated genes), down (number of downregulated genes, Z-score (z-score as calculated in Material and Methods), -log10(p) (negative log10 of p-value) and Contrast (compared tissues and genotypes).

| pathway | pval | padj | log2err | ES | NES | size | leadingEdge | Contrast |
|---|---|---|---|---|---|---|---|---|
| translation | 1.7368E-20 | 2.2578E-18 | 1.16906997 | -0.65976213 | -2.39568071 | 298 | | WTsB_vs_MUs |
| protein ubiquitination | 0.00042588 | 0.01384116 | 0.49849311 | -0.62174043 | -1.88173345 | 72 | | WTsB_vs_MUs |
| lignin catabolic process | 0.00018242 | 0.00790501 | 0.51884808 | 0.85331643 | 1.82771042 | 16 | | WTsB_vs_MUs |
| proteolysis involved in cellular protein catabolic process | 0.00015649 | 0.00790501 | 0.51884808 | -0.81186021 | -1.88758127 | 19 | | WTsB_vs_MUs |

Supplemental Table S 10: Enriched terms in GO enrichment analysis of proteome with topGO comparing wild type green leaves versus white *canal-1* leaves. Columns show ontology (BP: biological process, MF: molecular function, CC: cellular component), ID (Ontology identifier), Term (Ontology term name), (adjusted p-value), count (total number of enriched genes), up (number of upregulated genes), down (number of downregulated genes, Z-score (z-score as calculated in Material and Methods), -log10(p) (negative log10 of p-value) and Contrast (compared tissues and genotypes).

| Ontology | ID | Term | adj_pval | count | up | down | Z-score | -log10(p) | Contrast |
|---|---|---|---|---|---|---|---|---|---|
| BP | GO:0015979 | photosynthesis | 5.2E-23 | 124 | 4 | 120 | -10.41710752 | 22.28399666 | WTglB_vs_MUwl |
| BP | GO:0009768 | photosynthesis, light harvesting in phot... | 1.95E-15 | 28 | 0 | 28 | -5.291502622 | 14.70996539 | WTglB_vs_MUwl |
| BP | GO:0009767 | photosynthetic electron transport chain | 1.76667E-08 | 29 | 0 | 29 | -5.385164807 | 7.752845385 | WTglB_vs_MUwl |
| BP | GO:0006412 | translation | 0.000000215 | 179 | 174 | 5 | 12.63165307 | 6.66756154 | WTglB_vs_MUwl |
| BP | GO:0009416 | response to light stimulus | 0.00000056 | 33 | 2 | 31 | -5.048252023 | 6.251811973 | WTglB_vs_MUwl |
| BP | GO:0009772 | photosynthetic electron transport in pho... | 1.51667E-06 | 10 | 0 | 10 | -3.16227766 | 5.819109858 | WTglB_vs_MUwl |
| BP | GO:0006457 | protein folding | 1.85714E-06 | 64 | 63 | 1 | 7.75 | 5.731154688 | WTglB_vs_MUwl |
| BP | GO:0006418 | tRNA aminoacylation for protein translat... | 0.00003125 | 25 | 24 | 1 | 4.6 | 4.505149978 | WTglB_vs_MUwl |
| BP | GO:0006032 | chitin catabolic process | 0.000466667 | 10 | 10 | 0 | 3.16227766 | 3.330993219 | WTglB_vs_MUwl |
| BP | GO:0010498 | proteasomal protein catabolic process | 0.0008 | 43 | 43 | 0 | 6.557438524 | 3.096910013 | WTglB_vs_MUwl |
| BP | GO:0002181 | cytoplasmic translation | 0.000909091 | 36 | 36 | 0 | 6 | 3.041392685 | WTglB_vs_MUwl |
| BP | GO:0042026 | protein refolding | 0.001 | 17 | 16 | 1 | 3.638034376 | 3 | WTglB_vs_MUwl |
| BP | GO:0045036 | protein targeting to chloroplast | 0.001076923 | 11 | 10 | 1 | 2.713602101 | 2.967815317 | WTglB_vs_MUwl |
| BP | GO:0000398 | mRNA splicing, via spliceosome | 0.0015 | 48 | 48 | 0 | 6.92820323 | 2.823908741 | WTglB_vs_MUwl |
| BP | GO:0051085 | chaperone cofactor-dependent protein ref... | 0.0020625 | 14 | 14 | 0 | 3.741657387 | 2.685606043 | WTglB_vs_MUwl |
| BP | GO:0006414 | translational elongation | 0.0020625 | 14 | 14 | 0 | 3.741657387 | 2.685606043 | WTglB_vs_MUwl |
| BP | GO:0001732 | formation of cytoplasmic translation ini... | 0.005176471 | 11 | 11 | 0 | 3.31662479 | 2.285966249 | WTglB_vs_MUwl |
| BP | GO:0019684 | photosynthesis, light reaction | 0.005444444 | 68 | 0 | 68 | -8.246211251 | 2.264046429 | WTglB_vs_MUwl |
| BP | GO:0000463 | maturation of LSU-rRNA from tricistronic... | 0.007210526 | 9 | 9 | 0 | 3 | 2.142033034 | WTglB_vs_MUwl |
| BP | GO:0016998 | cell wall macromolecule catabolic proces... | 0.01195 | 9 | 9 | 0 | 3 | 1.922632095 | WTglB_vs_MUwl |
| BP | GO:0043161 | proteasome-mediated ubiquitin-dependent ... | 0.016380952 | 30 | 30 | 0 | 5.477225575 | 1.785660852 | WTglB_vs_MUwl |
| BP | GO:0015986 | proton motive force-driven ATP synthesis | 0.018090909 | 13 | 1 | 12 | -3.050851079 | 1.742539609 | WTglB_vs_MUwl |
| BP | GO:0006573 | valine metabolic process | 0.019608696 | 6 | 6 | 0 | 2.449489743 | 1.707551294 | WTglB_vs_MUwl |
| BP | GO:0000387 | spliceosomal snRNP assembly | 0.023708333 | 10 | 10 | 0 | 3.16227766 | 1.625098975 | WTglB_vs_MUwl |
| BP | GO:0019852 | L-ascorbic acid metabolic process | 0.02424 | 6 | 6 | 0 | 2.449489743 | 1.615467385 | WTglB_vs_MUwl |
| BP | GO:0034976 | response to endoplasmic reticulum stress | 0.024518519 | 16 | 16 | 0 | 4 | 1.610505775 | WTglB_vs_MUwl |
| BP | GO:0006241 | CTP biosynthetic process | 0.024518519 | 7 | 6 | 1 | 1.889822365 | 1.610505775 | WTglB_vs_MUwl |
| BP | GO:0065002 | intracellular protein transmembrane tran... | 0.027666667 | 16 | 14 | 2 | 3 | 1.558043162 | WTglB_vs_MUwl |
| BP | GO:0007005 | mitochondrion organization | 0.027666667 | 21 | 21 | 0 | 4.582575695 | 1.558043162 | WTglB_vs_MUwl |
| BP | GO:0032543 | mitochondrial translation | 0.027666667 | 10 | 10 | 0 | 3.16227766 | 1.558043162 | WTglB_vs_MUwl |
| BP | GO:0046351 | disaccharide biosynthetic process | 0.031648649 | 8 | 5 | 3 | 0.707106781 | 1.499644829 | WTglB_vs_MUwl |
| BP | GO:0045892 | negative regulation of DNA-templated tra... | 0.031648649 | 7 | 7 | 0 | 2.645751311 | 1.499644829 | WTglB_vs_MUwl |
| BP | GO:0010731 | protein glutathionylation | 0.031648649 | 6 | 6 | 0 | 2.449489743 | 1.499644829 | WTglB_vs_MUwl |
| BP | GO:0031167 | rRNA methylation | 0.031648649 | 6 | 6 | 0 | 2.449489743 | 1.499644829 | WTglB_vs_MUwl |
| BP | GO:0033962 | P-body assembly | 0.031648649 | 6 | 6 | 0 | 2.449489743 | 1.499644829 | WTglB_vs_MUwl |
| BP | GO:0000027 | ribosomal large subunit assembly | 0.031648649 | 12 | 12 | 0 | 3.464101615 | 1.499644829 | WTglB_vs_MUwl |
| BP | GO:0000413 | protein peptidyl-prolyl isomerization | 0.031648649 | 12 | 9 | 3 | 1.732050808 | 1.499644829 | WTglB_vs_MUwl |
| BP | GO:0006541 | glutamine metabolic process | 0.035578947 | 9 | 9 | 0 | 3 | 1.448806905 | WTglB_vs_MUwl |
| BP | GO:0006108 | malate metabolic process | 0.039410256 | 8 | 7 | 1 | 2.121320344 | 1.40439074 | WTglB_vs_MUwl |

| | | | | | | | | | |
|---|---|---|---|---|---|---|---|---|---|
| BP | GO:0001522 | pseudouridine synthesis | 0.046195122 | 6 | 6 | 0 | 2.449489743 | 1.335403882 | WTglB_vs_MUwl |
| BP | GO:0000028 | ribosomal small subunit assembly | 0.046195122 | 9 | 9 | 0 | 3 | 1.335403882 | WTglB_vs_MUwl |
| MF | GO:0140662 | ATP-dependent protein folding chaperone | 9.9E-11 | 33 | 32 | 1 | 5.396407335 | 10.00436481 | WTglB_vs_MUwl |
| MF | GO:0003723 | RNA binding | 3.8115E-09 | 243 | 239 | 4 | 15.07525703 | 8.418904076 | WTglB_vs_MUwl |
| MF | GO:0003735 | structural constituent of ribosome | 1.023E-08 | 117 | 113 | 4 | 10.07705356 | 7.990124366 | WTglB_vs_MUwl |
| MF | GO:0045156 | electron transporter, transferring elect... | 8.415E-07 | 12 | 0 | 12 | -3.464101615 | 6.07494588 | WTglB_vs_MUwl |
| MF | GO:0016887 | ATP hydrolysis activity | 0.00002574 | 54 | 50 | 4 | 6.25980712 | 4.589391457 | WTglB_vs_MUwl |
| MF | GO:0048038 | quinone binding | 3.11143E-05 | 17 | 0 | 17 | -4.123105626 | 4.507040165 | WTglB_vs_MUwl |
| MF | GO:0004812 | aminoacyl-tRNA ligase activity | 3.11143E-05 | 25 | 24 | 1 | 4.6 | 4.507040165 | WTglB_vs_MUwl |
| MF | GO:0051082 | unfolded protein binding | 0.000185625 | 32 | 32 | 0 | 5.656854249 | 3.731363533 | WTglB_vs_MUwl |
| MF | GO:0036402 | proteasome-activating activity | 0.000297 | 11 | 11 | 0 | 3.31662479 | 3.527243551 | WTglB_vs_MUwl |
| MF | GO:0070180 | large ribosomal subunit rRNA binding | 0.0006138 | 11 | 11 | 0 | 3.31662479 | 3.211973116 | WTglB_vs_MUwl |
| MF | GO:0003755 | peptidyl-prolyl cis-trans isomerase acti... | 0.00135 | 22 | 14 | 8 | 1.279204298 | 2.869666232 | WTglB_vs_MUwl |
| MF | GO:0016018 | cyclosporin A binding | 0.00165 | 8 | 8 | 0 | 2.828427125 | 2.782516056 | WTglB_vs_MUwl |
| MF | GO:0003746 | translation elongation factor activity | 0.002284615 | 13 | 13 | 0 | 3.605551275 | 2.641186903 | WTglB_vs_MUwl |
| MF | GO:0008061 | chitin binding | 0.005148 | 7 | 7 | 0 | 2.645751311 | 2.288361462 | WTglB_vs_MUwl |
| MF | GO:0004568 | chitinase activity | 0.005148 | 10 | 10 | 0 | 3.16227766 | 2.288361462 | WTglB_vs_MUwl |
| MF | GO:0003729 | mRNA binding | 0.010395 | 63 | 60 | 3 | 7.181324987 | 1.983175506 | WTglB_vs_MUwl |
| MF | GO:0017069 | snRNA binding | 0.02068 | 8 | 8 | 0 | 2.828427125 | 1.684449466 | WTglB_vs_MUwl |
| MF | GO:0003724 | RNA helicase activity | 0.02068 | 21 | 20 | 1 | 4.146139914 | 1.684449466 | WTglB_vs_MUwl |
| MF | GO:0016655 | oxidoreductase activity, acting on NAD(P... | 0.022040526 | 15 | 1 | 14 | -3.356585567 | 1.656778039 | WTglB_vs_MUwl |
| MF | GO:0005524 | ATP binding | 0.026928 | 202 | 169 | 33 | 9.568926608 | 1.569795901 | WTglB_vs_MUwl |
| MF | GO:0015038 | glutathione disulfide oxidoreductase act... | 0.02937 | 8 | 8 | 0 | 2.828427125 | 1.532096053 | WTglB_vs_MUwl |
| MF | GO:0016615 | malate dehydrogenase activity | 0.042053478 | 8 | 7 | 1 | 2.121320344 | 1.376198078 | WTglB_vs_MUwl |
| MF | GO:0031593 | polyubiquitin modification-dependent pro... | 0.042053478 | 8 | 8 | 0 | 2.828427125 | 1.376198078 | WTglB_vs_MUwl |
| MF | GO:0140492 | metal-dependent deubiquitinase activity | 0.0438372 | 6 | 6 | 0 | 2.449489743 | 1.358157193 | WTglB_vs_MUwl |
| MF | GO:0008137 | NADH dehydrogenase (ubiquinone) activity | 0.0438372 | 10 | 0 | 10 | -3.16227766 | 1.358157193 | WTglB_vs_MUwl |
| MF | GO:0042393 | histone binding | 0.046187308 | 13 | 13 | 0 | 3.605551275 | 1.335477353 | WTglB_vs_MUwl |

Supplemental Table S 11: Enriched terms in GO enrichment analysis of proteome with topGO comparing *canal-1* green leaves versus white *canal-1* leaves. Columns show ontology (BP: biological process, MF: molecular function, CC: cellular component), ID (Ontology identifier), Term (Ontology term name), (adjusted p-value), count (total number of enriched genes), up (number of upregulated genes), down (number of downregulated genes, Z-score (z-score as calculated in Material and Methods), -log10(p) (negative log10 of p-value) and Contrast (compared tissues and genotypes).

| Ontology | ID | Term | adj_pval | count | up | down | Z-score | -log10(p) | Contrast |
|---|---|---|---|---|---|---|---|---|---|
| BP | GO:0015979 | photosynthesis | 3.1E-12 | 90 | 4 | 86 | -8.643558938 | 11.50863831 | MUglB_vs_MUwl |
| BP | GO:0009767 | photosynthetic electron transport chain | 0.00000125 | 25 | 0 | 25 | -5 | 5.903089987 | MUglB_vs_MUwl |
| BP | GO:0006457 | protein folding | 1.56667E-06 | 52 | 52 | 0 | 7.211102551 | 5.805023397 | MUglB_vs_MUwl |
| BP | GO:0009768 | photosynthesis, light harvesting in phot... | 0.00000215 | 18 | 0 | 18 | -4.242640687 | 5.66756154 | MUglB_vs_MUwl |
| BP | GO:0009772 | photosynthetic electron transport in pho... | 0.0000102 | 9 | 0 | 9 | -3 | 4.991399828 | MUglB_vs_MUwl |
| BP | GO:0015986 | proton motive force-driven ATP synthesis | 0.001666667 | 14 | 3 | 11 | -2.138089935 | 2.77815125 | MUglB_vs_MUwl |
| BP | GO:0016998 | cell wall macromolecule catabolic proces... | 0.005222222 | 9 | 9 | 0 | 3 | 2.282144652 | MUglB_vs_MUwl |
| BP | GO:0010731 | protein glutathionylation | 0.005222222 | 7 | 6 | 1 | 1.889822365 | 2.282144652 | MUglB_vs_MUwl |
| BP | GO:0006032 | chitin catabolic process | 0.005222222 | 8 | 8 | 0 | 2.828427125 | 2.282144652 | MUglB_vs_MUwl |
| BP | GO:0051085 | chaperone cofactor-dependent protein ref... | 0.0059 | 12 | 12 | 0 | 3.464101615 | 2.229147988 | MUglB_vs_MUwl |
| BP | GO:0006412 | translation | 0.006545455 | 119 | 101 | 18 | 7.608597525 | 2.184060189 | MUglB_vs_MUwl |
| BP | GO:0000387 | spliceosomal snRNP assembly | 0.007714286 | 10 | 10 | 0 | 3.16227766 | 2.11270428 | MUglB_vs_MUwl |
| BP | GO:0043161 | proteasome-mediated ubiquitin-dependent ... | 0.007714286 | 31 | 31 | 0 | 5.567764363 | 2.11270428 | MUglB_vs_MUwl |
| BP | GO:0009416 | response to light stimulus | 0.007714286 | 20 | 1 | 19 | -4.024922359 | 2.11270428 | MUglB_vs_MUwl |
| BP | GO:0006310 | DNA recombination | 0.0084 | 10 | 10 | 0 | 3.16227766 | 2.075720714 | MUglB_vs_MUwl |
| BP | GO:0034976 | response to endoplasmic reticulum stress | 0.010375 | 18 | 18 | 0 | 4.242640687 | 1.984011895 | MUglB_vs_MUwl |
| BP | GO:0019852 | L-ascorbic acid metabolic process | 0.010705882 | 6 | 6 | 0 | 2.449489743 | 1.970377533 | MUglB_vs_MUwl |
| BP | GO:1990542 | mitochondrial transmembrane transport | 0.015210526 | 14 | 11 | 3 | 2.138089935 | 1.817855758 | MUglB_vs_MUwl |
| BP | GO:0034620 | cellular response to unfolded protein | 0.015210526 | 9 | 9 | 0 | 3 | 1.817855758 | MUglB_vs_MUwl |
| BP | GO:0010605 | negative regulation of macromolecule met... | 0.016857143 | 29 | 28 | 1 | 5.013774131 | 1.773216033 | MUglB_vs_MUwl |
| BP | GO:0045934 | negative regulation of nucleobase-contai... | 0.016857143 | 11 | 11 | 0 | 3.31662479 | 1.773216033 | MUglB_vs_MUwl |
| BP | GO:0031323 | regulation of cellular metabolic process | 0.017608696 | 53 | 45 | 8 | 5.082340866 | 1.754272813 | MUglB_vs_MUwl |
| BP | GO:0006418 | tRNA aminoacylation for protein translat... | 0.017608696 | 16 | 16 | 0 | 4 | 1.754272813 | MUglB_vs_MUwl |
| BP | GO:0051052 | regulation of DNA metabolic process | 0.01936 | 7 | 7 | 0 | 2.645751311 | 1.713094647 | MUglB_vs_MUwl |
| BP | GO:0006378 | mRNA polyadenylation | 0.01936 | 7 | 7 | 0 | 2.645751311 | 1.713094647 | MUglB_vs_MUwl |
| BP | GO:0006334 | nucleosome assembly | 0.02162963 | 11 | 11 | 0 | 3.31662479 | 1.664950917 | MUglB_vs_MUwl |
| BP | GO:0000027 | ribosomal large subunit assembly | 0.02162963 | 11 | 8 | 3 | 1.507556723 | 1.664950917 | MUglB_vs_MUwl |
| BP | GO:0010498 | proteasomal protein catabolic process | 0.025714286 | 39 | 39 | 0 | 6.244997998 | 1.589825535 | MUglB_vs_MUwl |
| BP | GO:0030433 | ubiquitin-dependent ERAD pathway | 0.026413793 | 11 | 11 | 0 | 3.31662479 | 1.578169228 | MUglB_vs_MUwl |
| BP | GO:0006414 | translational elongation | 0.036419355 | 10 | 10 | 0 | 3.16227766 | 1.438667752 | MUglB_vs_MUwl |
| BP | GO:0051276 | chromosome organization | 0.036419355 | 12 | 12 | 0 | 3.464101615 | 1.438667752 | MUglB_vs_MUwl |
| BP | GO:0000154 | rRNA modification | 0.0422 | 9 | 9 | 0 | 3 | 1.374687549 | MUglB_vs_MUwl |
| BP | GO:0016052 | carbohydrate catabolic process | 0.0422 | 26 | 22 | 4 | 3.530090432 | 1.374687549 | MUglB_vs_MUwl |
| BP | GO:0090501 | RNA phosphodiester bond hydrolysis | 0.0422 | 5 | 4 | 1 | 1.341640786 | 1.374687549 | MUglB_vs_MUwl |
| BP | GO:0045892 | negative regulation of DNA-templated tra... | 0.0422 | 6 | 6 | 0 | 2.449489743 | 1.374687549 | MUglB_vs_MUwl |
| BP | GO:0042026 | protein refolding | 0.043888889 | 11 | 11 | 0 | 3.31662479 | 1.357645414 | MUglB_vs_MUwl |
| MF | GO:0016168 | chlorophyll binding | 2.2E-19 | 40 | 0 | 40 | -6.32455532 | 18.65757732 | MUglB_vs_MUwl |
| MF | GO:0140662 | ATP-dependent protein folding chaperone | 0.00000009 | 26 | 26 | 0 | 5.099019514 | 7.045757491 | MUglB_vs_MUwl |
| MF | GO:0045156 | electron transporter, transferring elect... | 2.26667E-06 | 11 | 0 | 11 | -3.31662479 | 5.644612342 | MUglB_vs_MUwl |
| MF | GO:0016887 | ATP hydrolysis activity | 0.00000475 | 48 | 45 | 3 | 6.062177826 | 5.32330639 | MUglB_vs_MUwl |

| | | | | | | | | | |
|---|---|---|---|---|---|---|---|---|---|
| MF | GO:0003723 | RNA binding | 0.0000048 | 190 | 172 | 18 | 11.17233425 | 5.318758763 | MUglB_vs_MUwl |
| MF | GO:0036402 | proteasome-activating activity | 0.00004 | 11 | 11 | 0 | 3.31662479 | 4.397940009 | MUglB_vs_MUwl |
| MF | GO:0030527 | structural constituent of chromatin | 9.57143E-05 | 19 | 19 | 0 | 4.358898944 | 4.019023237 | MUglB_vs_MUwl |
| MF | GO:0051082 | unfolded protein binding | 0.00011625 | 28 | 28 | 0 | 5.291502622 | 3.934607038 | MUglB_vs_MUwl |
| MF | GO:0003735 | structural constituent of ribosome | 0.000477778 | 82 | 65 | 17 | 5.300713252 | 3.320774054 | MUglB_vs_MUwl |
| MF | GO:0008422 | beta-glucosidase activity | 0.001818182 | 15 | 15 | 0 | 3.872983346 | 2.740362689 | MUglB_vs_MUwl |
| MF | GO:0015038 | glutathione disulfide oxidoreductase act... | 0.001818182 | 9 | 9 | 0 | 3 | 2.740362689 | MUglB_vs_MUwl |
| MF | GO:0046982 | protein heterodimerization activity | 0.00425 | 18 | 18 | 0 | 4.242640687 | 2.37161107 | MUglB_vs_MUwl |
| MF | GO:0051787 | misfolded protein binding | 0.004923077 | 9 | 9 | 0 | 3 | 2.307763378 | MUglB_vs_MUwl |
| MF | GO:0003756 | protein disulfide isomerase activity | 0.005357143 | 8 | 7 | 1 | 2.121320344 | 2.271066772 | MUglB_vs_MUwl |
| MF | GO:0046912 | acyltransferase activity, acyl groups co... | 0.014133333 | 8 | 5 | 3 | 0.707106781 | 1.849755398 | MUglB_vs_MUwl |
| MF | GO:0003729 | mRNA binding | 0.017421053 | 51 | 45 | 6 | 5.461092328 | 1.758925607 | MUglB_vs_MUwl |
| MF | GO:0016018 | cyclosporin A binding | 0.017421053 | 6 | 6 | 0 | 2.449489743 | 1.758925607 | MUglB_vs_MUwl |
| MF | GO:0004812 | aminoacyl-tRNA ligase activity | 0.017421053 | 16 | 16 | 0 | 4 | 1.758925607 | MUglB_vs_MUwl |
| MF | GO:0004568 | chitinase activity | 0.017421053 | 8 | 8 | 0 | 2.828427125 | 1.758925607 | MUglB_vs_MUwl |
| MF | GO:0017069 | snRNA binding | 0.0215 | 7 | 7 | 0 | 2.645751311 | 1.66756154 | MUglB_vs_MUwl |
| MF | GO:0003677 | DNA binding | 0.035428571 | 70 | 69 | 1 | 8.127554543 | 1.450646359 | MUglB_vs_MUwl |

Supplemental Table S 12: Enriched terms in KEGG enrichment analysis of proteome with clusterProfiler comparing wild type green leaves versus white *canal-1* leaves. Columns show ontology (BP: biological process, MF: molecular function, CC: cellular component), ID (Ontology identifier), Term (Ontology term name), (adjusted p-value), count (total number of enriched genes), up (number of upregulated genes), down (number of downregulated genes, Z-score (z-score as calculated in Material and Methods), -log10(p) (negative log10 of p-value) and Contrast (compared tissues and genotypes).

| ID | Description | interm | all | BgRatio | Count | pvalue | p.adjust | qvalue | Gene Ratio | logp | Contrast |
|---|---|---|---|---|---|---|---|---|---|---|---|
| sly03430 | Mismatch repair | 7 | 430 | 18/3490 | 7 | 0.003862295 | 0.042485245 | 0.038848816 | 0.01627907 | 1.371761869 | WTglB_vs_MUwl |
| sly00970 | Aminoacyl-tRNA biosynthesis | 13 | 430 | 30/3490 | 13 | 2.08125E-05 | 0.000294349 | 0.000269155 | 0.030232558 | 3.531137692 | WTglB_vs_MUwl |
| sly00196 | Photosynthesis - antenna proteins | 15 | 430 | 19/3490 | 15 | 4.44281E-11 | 1.09959E-09 | 1.00548E-09 | 0.034883721 | 8.958767466 | WTglB_vs_MUwl |
| sly03018 | RNA degradation | 16 | 430 | 60/3490 | 16 | 0.001812458 | 0.022429164 | 0.020509389 | 0.037209302 | 1.649186922 | WTglB_vs_MUwl |
| sly03050 | Proteasome | 17 | 430 | 33/3490 | 17 | 4.55899E-08 | 9.0268E-07 | 8.25417E-07 | 0.039534884 | 6.044466028 | WTglB_vs_MUwl |
| sly03013 | Nucleocytoplasmic transport | 21 | 430 | 64/3490 | 21 | 1.25943E-05 | 0.000207807 | 0.00019002 | 0.048837209 | 3.682340442 | WTglB_vs_MUwl |
| sly03040 | Spliceosome | 38 | 430 | 96/3490 | 38 | 5.45418E-12 | 1.79988E-10 | 1.64582E-10 | 0.088372093 | 9.744756533 | WTglB_vs_MUwl |
| sly00195 | Photosynthesis | 42 | 430 | 101/3490 | 42 | 4.98408E-14 | 2.46712E-12 | 2.25595E-12 | 0.097674419 | 11.60780959 | WTglB_vs_MUwl |
| sly03010 | Ribosome | 74 | 430 | 246/3490 | 74 | 8.38634E-15 | 8.30248E-13 | 7.59185E-13 | 0.172093023 | 12.08079225 | WTglB_vs_MUwl |
| ID | Description | interm | all | BgRatio | Count | pvalue | p.adjust | qvalue | Gene Ratio | logp | Contrast |
| sly03430 | Mismatch repair | 7 | 430 | 18/3490 | 7 | 0.003862295 | 0.042485245 | 0.038848816 | 0.01627907 | 1.371761869 | WTglB_vs_MUwl |
| sly00970 | Aminoacyl-tRNA biosynthesis | 13 | 430 | 30/3490 | 13 | 2.08125E-05 | 0.000294349 | 0.000269155 | 0.030232558 | 3.531137692 | WTglB_vs_MUwl |
| sly00196 | Photosynthesis - antenna proteins | 15 | 430 | 19/3490 | 15 | 4.44281E-11 | 1.09959E-09 | 1.00548E-09 | 0.034883721 | 8.958767466 | WTglB_vs_MUwl |
| sly03018 | RNA degradation | 16 | 430 | 60/3490 | 16 | 0.001812458 | 0.022429164 | 0.020509389 | 0.037209302 | 1.649186922 | WTglB_vs_MUwl |

Supplemental Table S 13: Enriched terms in KEGG enrichment analysis of proteome with clusterProfiler comparing *canal-1* green leaves versus white *canal-1* leaves. Columns show ontology (BP: biological process, MF: molecular function, CC: cellular component), ID (Ontology identifier), Term (Ontology term name), (adjusted p-value), count (total number of enriched genes), up (number of upregulated genes), down (number of downregulated genes, Z-score (z-score as calculated in Material and Methods), -log10(p) (negative log10 of p-value) and Contrast (compared tissues and genotypes).

| ID | Description | interm | all | BgRatio | Count | pvalue | p.adjust | qvalue | Gene Ratio | logp | Contrast |
|---|---|---|---|---|---|---|---|---|---|---|---|
| sly00970 | Aminoacyl-tRNA biosynthesis | 9 | 365 | 30/3490 | 9 | 0.002627709 | 0.035661758 | 0.030821243 | 0.024657534 | 1.447797248 | MUglB_vs_MUwl |
| sly00196 | Photosynthesis - antenna proteins | 10 | 365 | 19/3490 | 10 | 5.40038E-06 | 0.000102607 | 8.86799E-05 | 0.02739726 | 3.988822238 | MUglB_vs_MUwl |
| sly03050 | Proteasome | 14 | 365 | 33/3490 | 14 | 1.87247E-06 | 4.44712E-05 | 3.84349E-05 | 0.038356164 | 4.351921528 | MUglB_vs_MUwl |
| sly03013 | Nucleocytoplasmic transport | 18 | 365 | 64/3490 | 18 | 5.7167E-05 | 0.000905144 | 0.000782285 | 0.049315068 | 3.043282167 | MUglB_vs_MUwl |
| sly03040 | Spliceosome | 32 | 365 | 96/3490 | 32 | 6.44858E-10 | 3.06307E-08 | 2.64731E-08 | 0.087671233 | 7.513842511 | MUglB_vs_MUwl |
| sly00195 | Photosynthesis | 33 | 365 | 101/3490 | 33 | 6.22321E-10 | 3.06307E-08 | 2.64731E-08 | 0.090410959 | 7.513842511 | MUglB_vs_MUwl |
| sly03010 | Ribosome | 53 | 365 | 246/3490 | 53 | 8.41499E-08 | 2.66475E-06 | 2.30305E-06 | 0.145205479 | 5.574343921 | MUglB_vs_MUwl |
| ID | Description | interm | all | BgRatio | Count | pvalue | p.adjust | qvalue | Gene Ratio | logp | Contrast |
| sly00970 | Aminoacyl-tRNA biosynthesis | 9 | 365 | 30/3490 | 9 | 0.002627709 | 0.035661758 | 0.030821243 | 0.024657534 | 1.447797248 | MUglB_vs_MUwl |
| sly00196 | Photosynthesis - antenna proteins | 10 | 365 | 19/3490 | 10 | 5.40038E-06 | 0.000102607 | 8.86799E-05 | 0.02739726 | 3.988822238 | MUglB_vs_MUwl |
| sly03050 | Proteasome | 14 | 365 | 33/3490 | 14 | 1.87247E-06 | 4.44712E-05 | 3.84349E-05 | 0.038356164 | 4.351921528 | MUglB_vs_MUwl |
| sly03013 | Nucleocytoplasmic transport | 18 | 365 | 64/3490 | 18 | 5.7167E-05 | 0.000905144 | 0.000782285 | 0.049315068 | 3.043282167 | MUglB_vs_MUwl |
| sly03040 | Spliceosome | 32 | 365 | 96/3490 | 32 | 6.44858E-10 | 3.06307E-08 | 2.64731E-08 | 0.087671233 | 7.513842511 | MUglB_vs_MUwl |
| sly00195 | Photosynthesis | 33 | 365 | 101/3490 | 33 | 6.22321E-10 | 3.06307E-08 | 2.64731E-08 | 0.090410959 | 7.513842511 | MUglB_vs_MUwl |
| sly03010 | Ribosome | 53 | 365 | 246/3490 | 53 | 8.41499E-08 | 2.66475E-06 | 2.30305E-06 | 0.145205479 | 5.574343921 | MUglB_vs_MUwl |
| ID | Description | interm | all | BgRatio | Count | pvalue | p.adjust | qvalue | Gene Ratio | logp | Contrast |

Supplemental Table S 14: Enriched pathways in enrichment analysis of metabolic pathways with Metaboanalyst comparing *canal-1* green leaves versus white *canal-1* leaves. Columns show pathway, ratio (ratio of mapped metabolites to total metabolites in pathway), total compounds (total compounds in pathway), hits (number of metabolites mapped to pathway), p-values, -log10(p) (negative log10 of p-value), holm (Holm-adjusted p-value), FDR (FDR-adjusted p-value), impact (pathway impact) and Contrast (compared tissues and genotypes).

| pathway | Full pathway name | ratio | total compounds | hits | p-values | -log10(p) | holm | FDR | impact | Contrast |
|---|---|---|---|---|---|---|---|---|---|---|
| Pant/Coa bio | Pantothenate and CoA biosynthesis | 0.17391304 | 23 | 4 | 7.69E-08 | 5.7721133 | 0.00000438 | 0.00000169 | 0.08423 | WTglB_vs_MUwl |
| Val/Leu/Ile bio | Valine, leucine and isoleucine biosynthesis | 0.27272727 | 22 | 6 | 1.18E-07 | 5.7721133 | 0.0000066 | 0.00000169 | 0.10727 | WTglB_vs_MUwl |
| GSH met | Glutathione metabolism | 0.26923077 | 26 | 7 | 5.86E-07 | 5.4225082 | 0.0000293 | 0.00000378 | 0.13362 | WTglB_vs_MUwl |
| Ino-phos met | Inositol phosphate metabolism | 0.10714286 | 28 | 3 | 0.00000203 | 4.97061622 | 0.0000972 | 0.0000107 | 0.12552 | WTglB_vs_MUwl |
| S met | Sulfur metabolism | 0.2 | 15 | 3 | 0.0000108 | 4.28399666 | 0.00050581 | 0.000052 | 0.10774 | WTglB_vs_MUwl |
| Arg/Pro met | Arginine and proline metabolism | 0.23529412 | 34 | 8 | 0.000036 | 3.82605656 | 0.0016212 | 0.00014926 | 0.5247 | WTglB_vs_MUwl |
| Glyox met | Glyoxylate and dicarboxylate metabolism | 0.37931034 | 29 | 11 | 0.0000467 | 3.77152265 | 0.0020144 | 0.00016923 | 0.50324 | WTglB_vs_MUwl |
| Tyr met | Tyrosine metabolism | 0.25 | 16 | 4 | 0.000094 | 3.56965077 | 0.0037586 | 0.00026937 | 0.27703 | WTglB_vs_MUwl |
| Phe/Tyr/Trp bio | Phenylalanine, tyrosine and tryptophan biosynthesis | 0.18181818 | 22 | 4 | 0.0001319 | 3.45875784 | 0.0048803 | 0.00034773 | 0.1016 | WTglB_vs_MUwl |
| Starch met | Starch and sucrose metabolism | 0.31818182 | 22 | 7 | 0.00033327 | 3.07550362 | 0.011998 | 0.00084042 | 0.72431 | WTglB_vs_MUwl |
| Glyc/Gluc | Glycolysis / Gluconeogenesis | 0.11538462 | 26 | 3 | 0.00047536 | 2.95655912 | 0.016638 | 0.0011052 | 0.12036 | WTglB_vs_MUwl |
| TCA cyc | Citrate cycle (TCA cycle) | 0.35 | 20 | 7 | 0.00047637 | 2.95655912 | 0.016638 | 0.0011052 | 0.32413 | WTglB_vs_MUwl |
| Glyc-lipid met | Glycerolipid metabolism | 0.19047619 | 21 | 4 | 0.0018387 | 2.43447094 | 0.05516 | 0.0036773 | 0.02235 | WTglB_vs_MUwl |
| Arg bio | Arginine biosynthesis | 0.44444444 | 18 | 8 | 0.003661 | 2.17812635 | 0.098847 | 0.0066355 | 0.34079 | WTglB_vs_MUwl |
| Aa-tRNA bio | Aminoacyl-tRNA biosynthesis | 0.41304348 | 46 | 19 | 0.0047869 | 2.07502846 | 0.12446 | 0.0084134 | 0.11111 | WTglB_vs_MUwl |
| Galact met | Galactose metabolism | 0.25925926 | 27 | 7 | 0.0091016 | 1.80894042 | 0.22754 | 0.015526 | 0.29394 | WTglB_vs_MUwl |
| But met | Butanoate metabolism | 0.29411765 | 17 | 5 | 0.0098481 | 1.78727985 | 0.23635 | 0.01632 | 0.13636 | WTglB_vs_MUwl |
| Ala/Asp/Glu met | Alanine, aspartate and glutamate metabolism | 0.45454545 | 22 | 10 | 0.017776 | 1.55494095 | 0.39107 | 0.027865 | 0.84892 | WTglB_vs_MUwl |
| CyanAA met | Cyanoamino acid metabolism | 0.20689655 | 29 | 6 | 0.023478 | 1.44569259 | 0.49304 | 0.035835 | 0.35593 | WTglB_vs_MUwl |
| Lys deg | Lysine degradation | 0.16666667 | 18 | 3 | 0.026983 | 1.40072843 | 0.53967 | 0.039744 | 0 | WTglB_vs_MUwl |

Supplemental Table S 15: Enriched pathways in enrichment analysis of metabolic pathways with Metaboanalyst comparing wild type green leaves versus white *canal-1* leaves. Columns show pathway, ratio (ratio of mapped metabolites to total metabolites in pathway), total compounds (total compounds in pathway), hits (number of metabolites mapped to pathway), p-values, -log10(p) (negative log10 of p-value), holm (Holm-adjusted p-value), FDR (FDR-adjusted p-value), impact (pathway impact) and Contrast (compared tissues and genotypes).

| pathway | Full pathway name | ratio | total compounds | hits | p-values | -log10(p) | holm | FDR | impact | Contrast |
|---|---|---|---|---|---|---|---|---|---|---|
| Ala/Asp/Glu met | Alanine, aspartate and glutamate metabolism | 0.45454545 | 22 | 10 | 0.0010156 | 1.62624231 | 0.05789 | 0.023646 | 0.84892 | WTglB_vs_MUgl |
| GSH met | Glutathione metabolism | 0.26923077 | 26 | 7 | 0.0014774 | 1.62624231 | 0.082733 | 0.023646 | 0.13362 | WTglB_vs_MUgl |
| CyanAA met | Cyanoamino acid metabolism | 0.20689655 | 29 | 6 | 0.0016307 | 1.62624231 | 0.089691 | 0.023646 | 0.35593 | WTglB_vs_MUgl |

Supplemental Table S 16: Results of wilcox-tests on network metrics degree, closeness and betweenness, comparing SCO2 value to the overall filtered network.

| Metric | SCO2 value | Confidence interval | P-value |
|---|---|---|---|
| degree | 29 | [16.9999415681835, 18.5000608797383] | 1.16E-102 |
| betweenness | 62977.03 | [19085.0180087324, 20299.3192423883] | 0 |
| closeness | 1.56E-05 | [4.64600052347922e-06, 1.78657776347781e-05] | 0 |

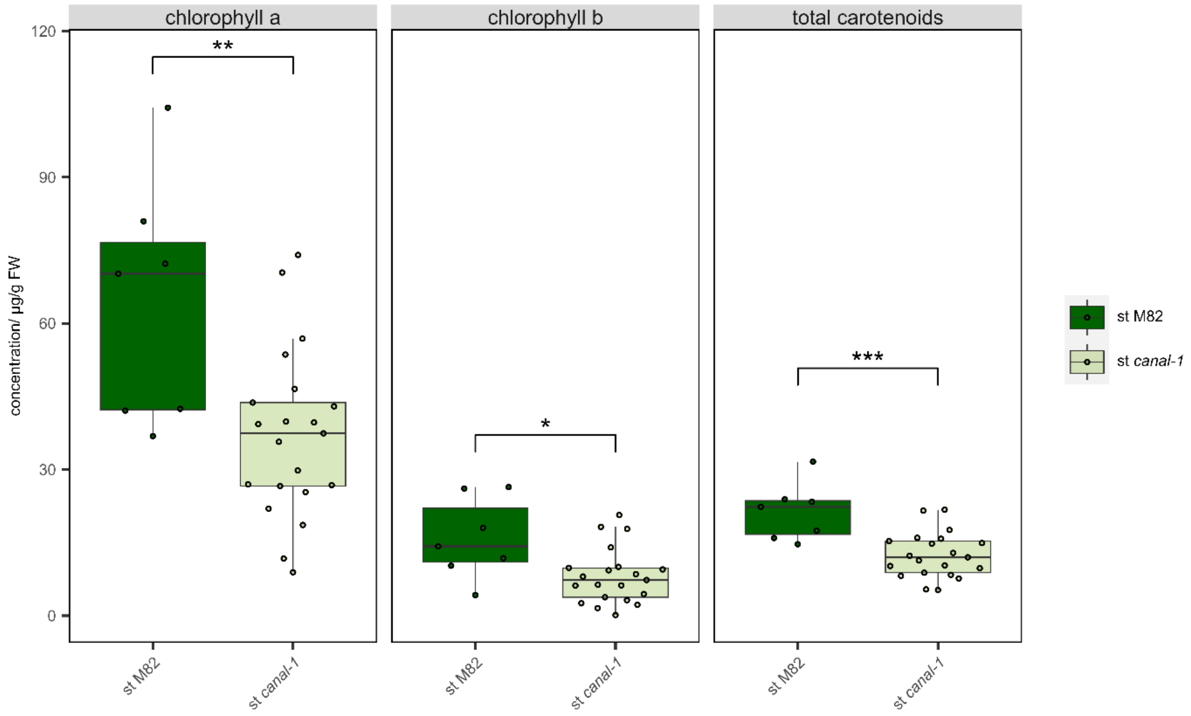

Supplemental Figure S 1: Pigments in stems of wild type and *canal-1* plants. Subpanels show different pigments. All boxplots show the interquartile range (IQR) between the first and the third quartile as the box, with the median indicated by a black center line. Whiskers extend from quartiles to most extreme points with a maximum of 1.5 x IQR. Points beyond that range are considered outliers but overlaid by individual data points, shown as quasirandom. Boxes and points are filled according to genotype (M82: green; *canal-1*: beige). Statistical significance was estimated by pairwise-wilcox and significant differences are indicated by a bracket and asterisks (*: $p \leq 0.05$; **: $p \leq 0.005$; ***: $p \leq 0.0005$).

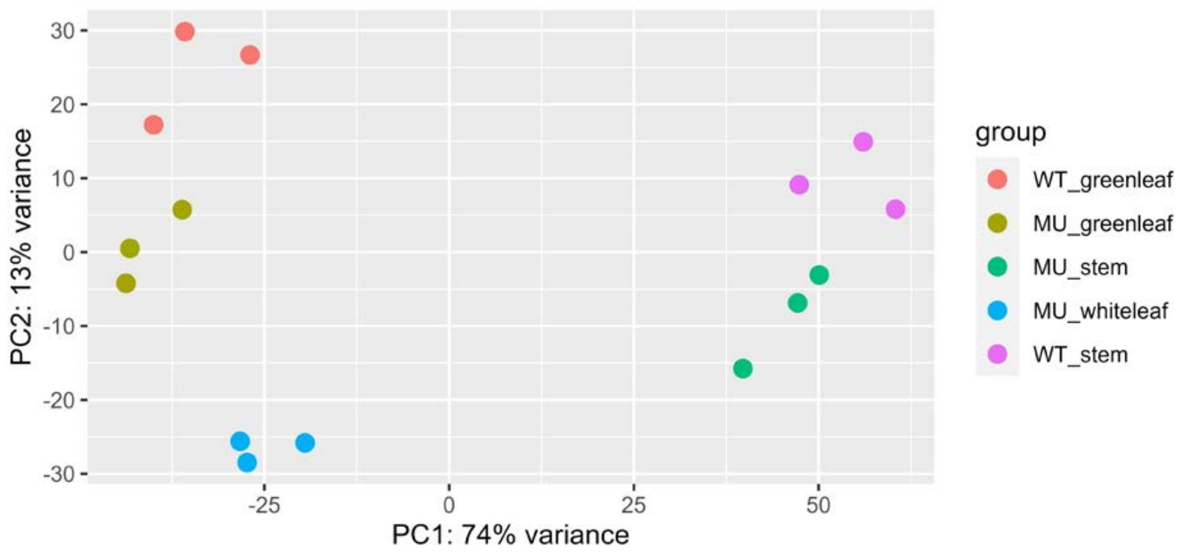

Supplemental Figure S 2: Principal component analysis of RNAseq data. Plot shows the first two principal components. Each dot comprises one sample and is colored according to tissue and genotype. WT: wild type; MU: mutant

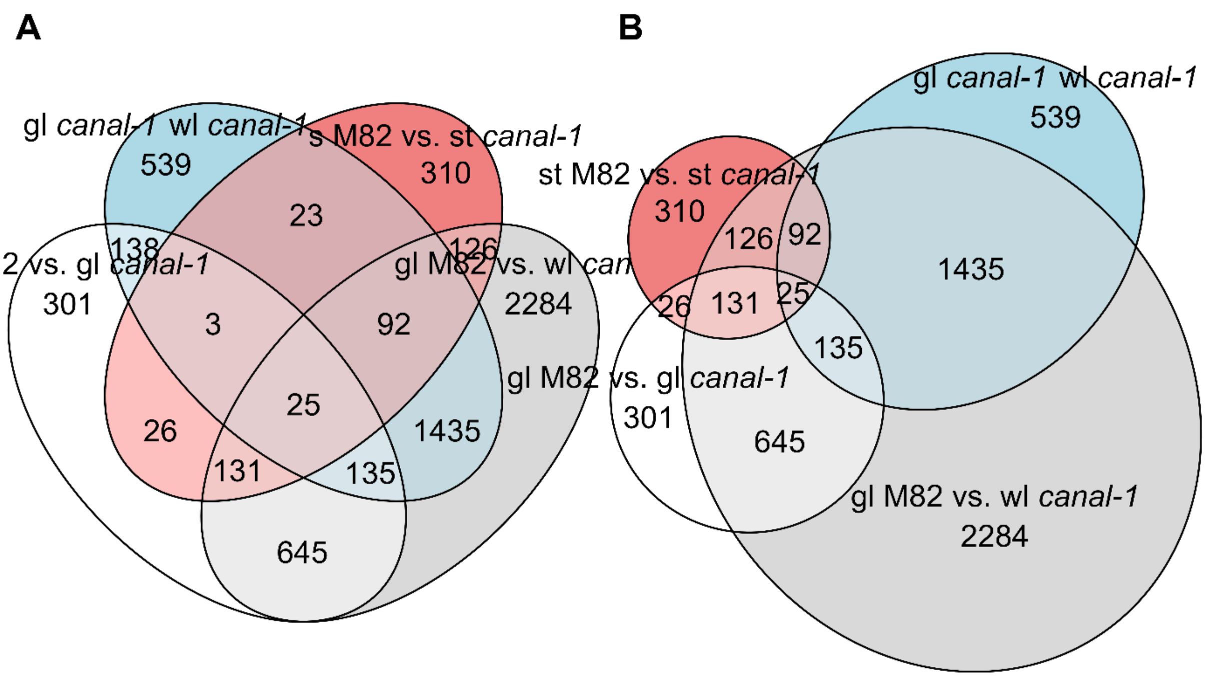

Supplemental Figure S 3: Differentially expressed genes in different contrasts
A, Venn and B, Euler plots showing number of DEGs in different contrasts. WTglB_vs_MU_wl: wild type green leaves versus white canal-1 leaves, WTglB_vs_MU_gl: wild type green leaves versus green canal-1 leaves, MUglB_vs_MU_gl: green canal-1 leaves versus white canal-1 leaves, WTsB_vs_MUs: wild type stems versus canal-1 stems

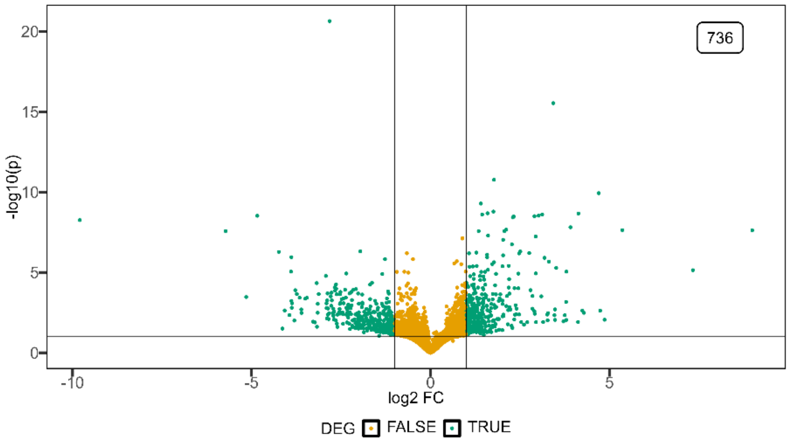

Supplemental Figure S 4: Volcano plot of differentially expressed genes comparing wild type and *canal-1* leaves. Differentially expressed genes are shown in green and non-DE genes are shown in yellow. Label in top right-hand corner shows the number of DE genes of the contrast. gl: green leaves; wl: white leaves (DEG: p ≤ 0.05 & |log2FC| ≥ 2).

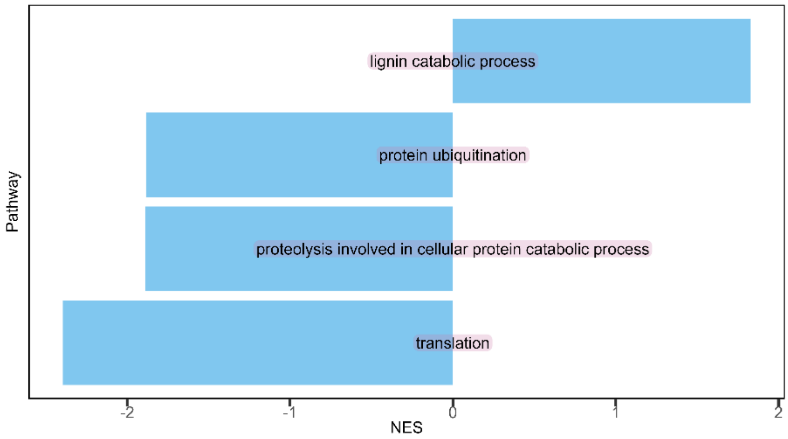

Supplemental Figure S 5: Barplot of enriched gene sets according to fgsea comparing wild type and *canal-1* stems. NES: normalized enrichment score.

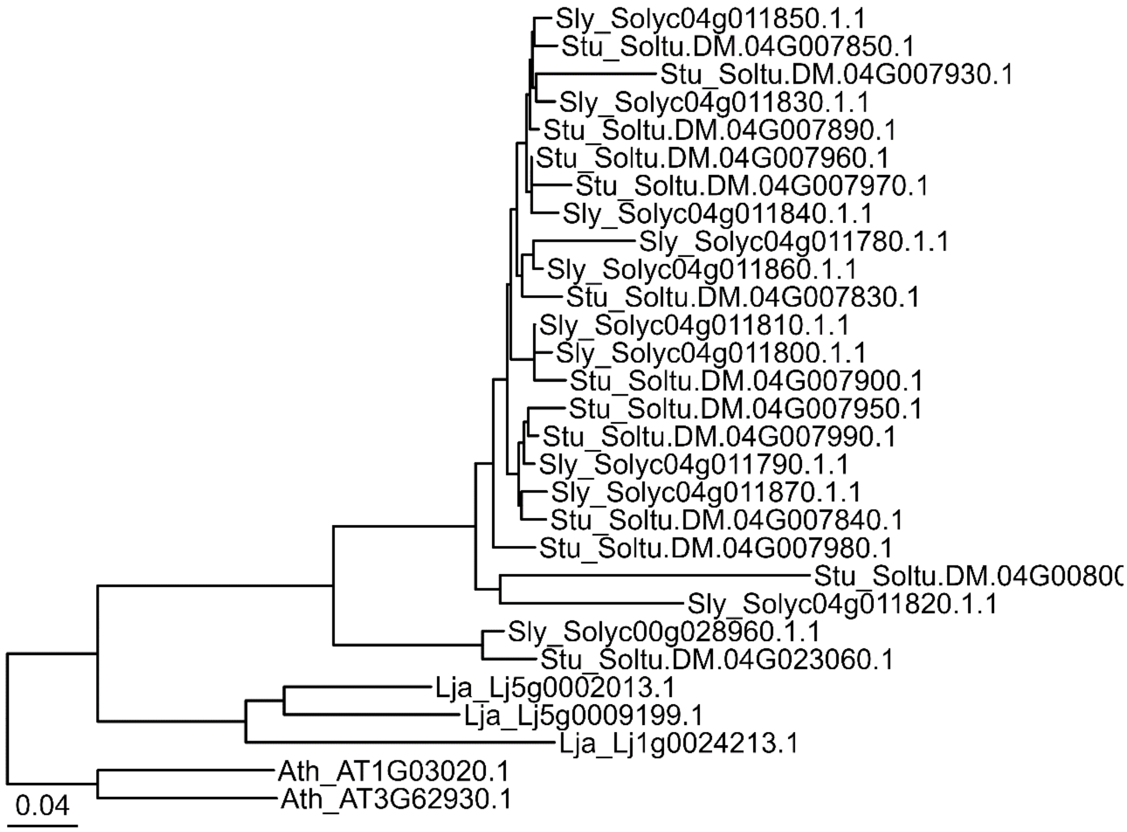

Supplemental Figure S 6: Phylogenetic tree of glutaredoxin genes orthogroup assembled by ggtree.

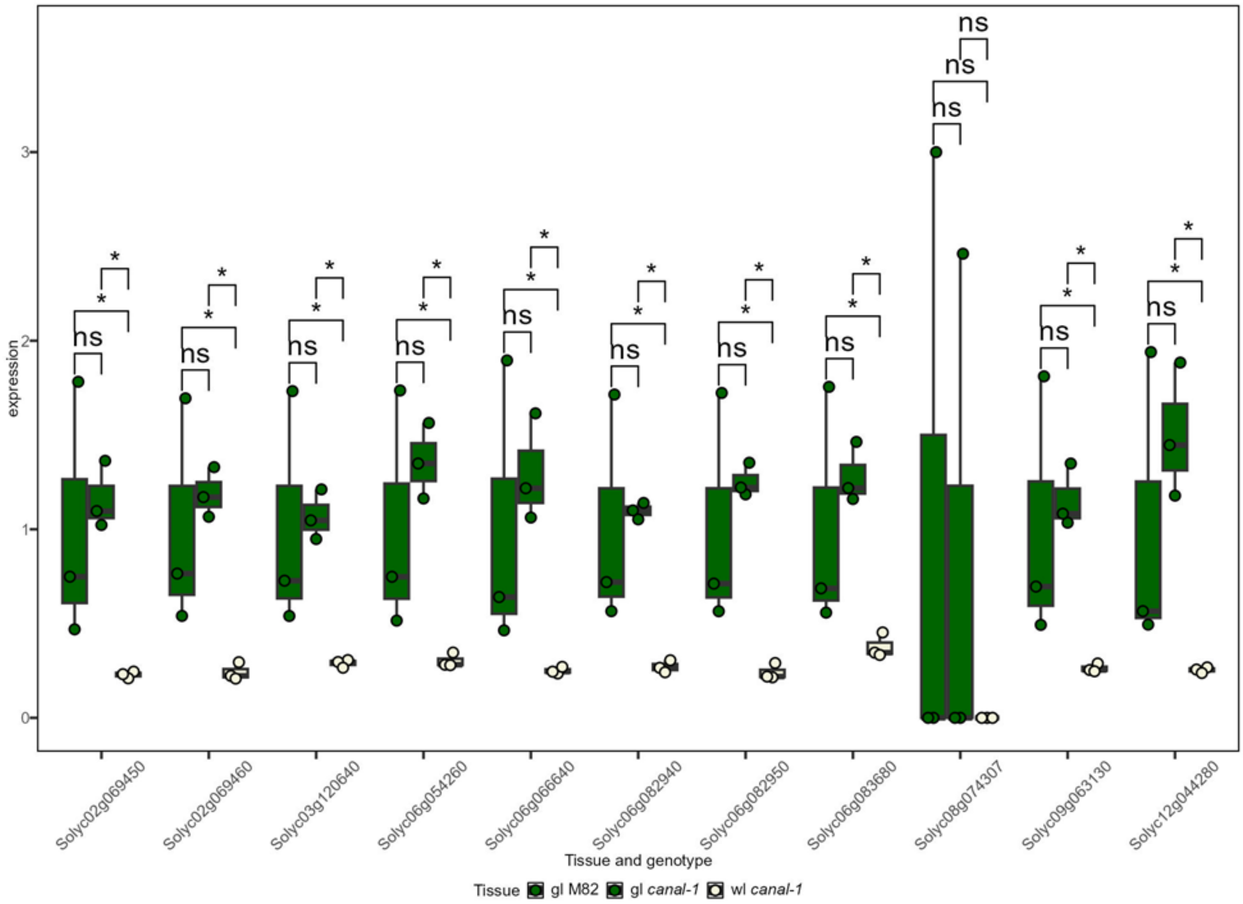

Supplemental Figure S 7: Expression of photosystem I reaction center genes. Boxes and points are filled according to tissue (green leaves: green, white leaves: beige). Statistical significance was extracted from differential gene expression analysis. Significant differences are indicated by a bracket and asterisks (*: $p \leq 0.05$ & $|\log2FC| \geq 2$).

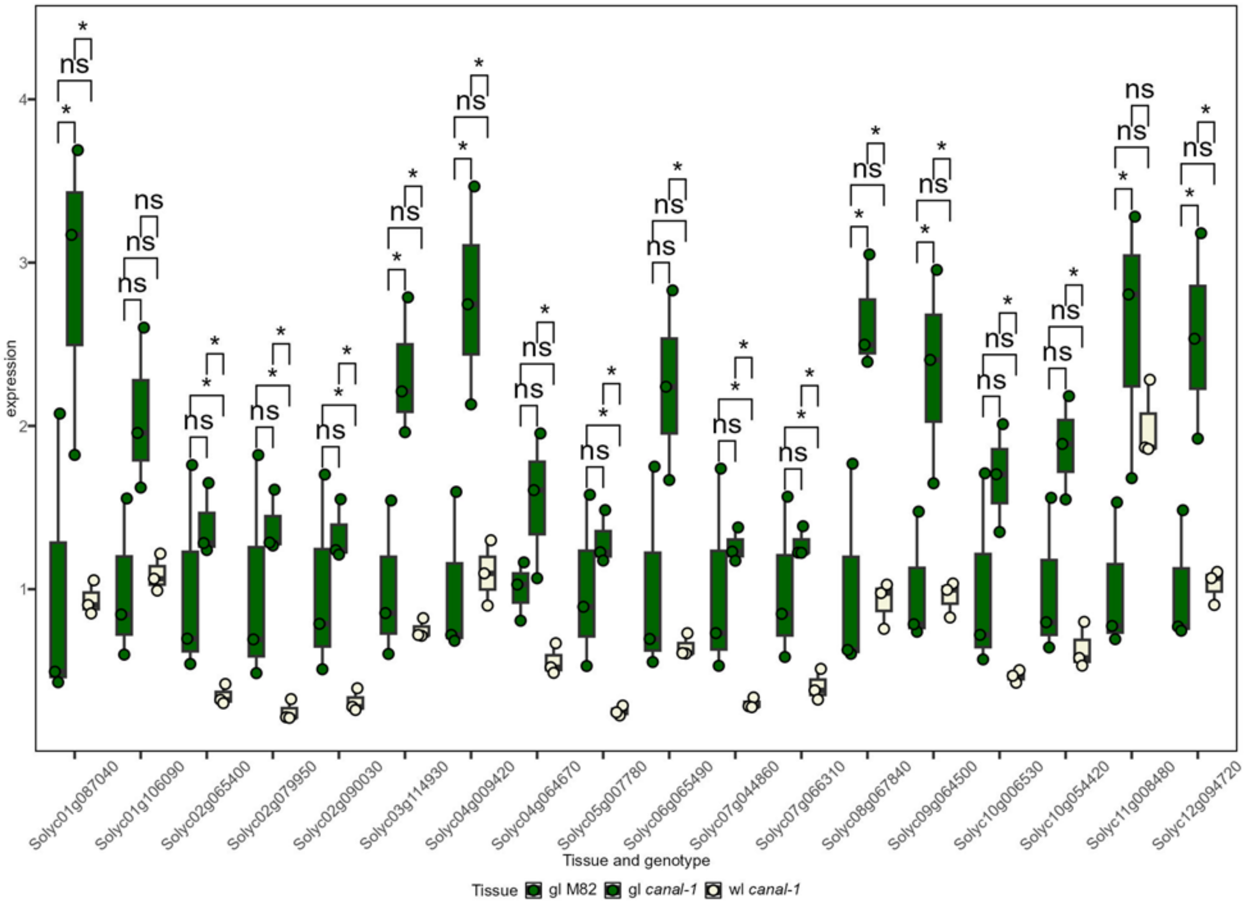

Supplemental Figure S 8: Expression of photosystem II oxygen evolving center genes. Boxes and points are filled according to tissue (green leaves: green, white leaves: beige). Statistical significance was extracted from differential gene expression analysis. Significant differences are indicated by a bracket and asterisks (*: p ≤ 0.05 & |log2FC| ≥ 2).

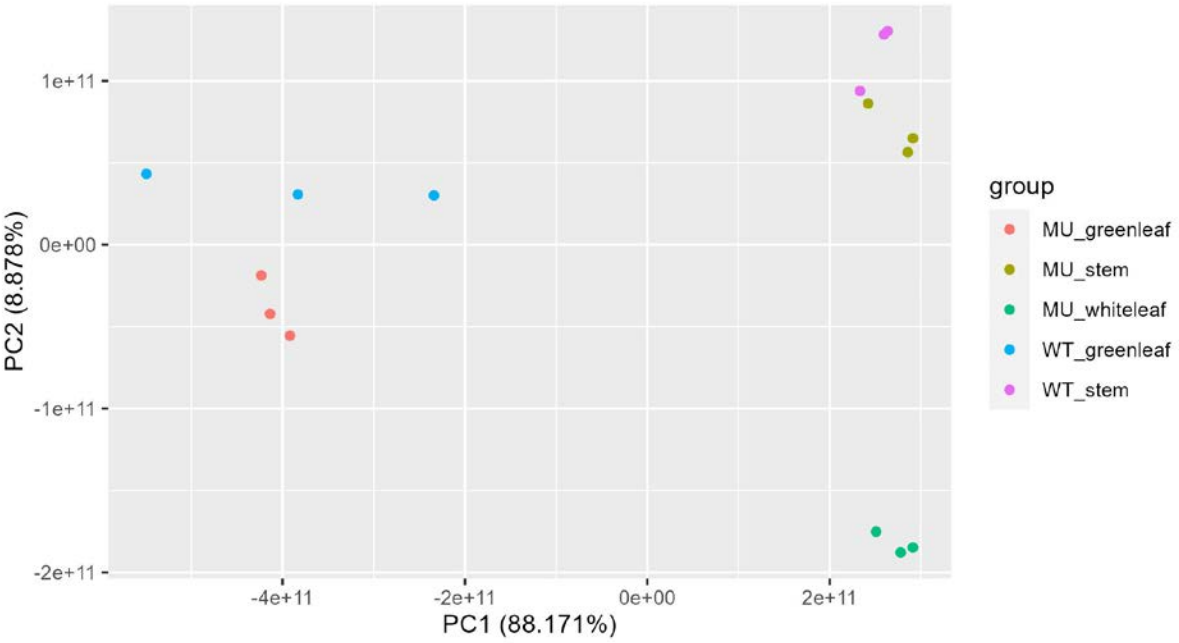

Supplemental Figure S 9: Principal component analysis of proteomics data.

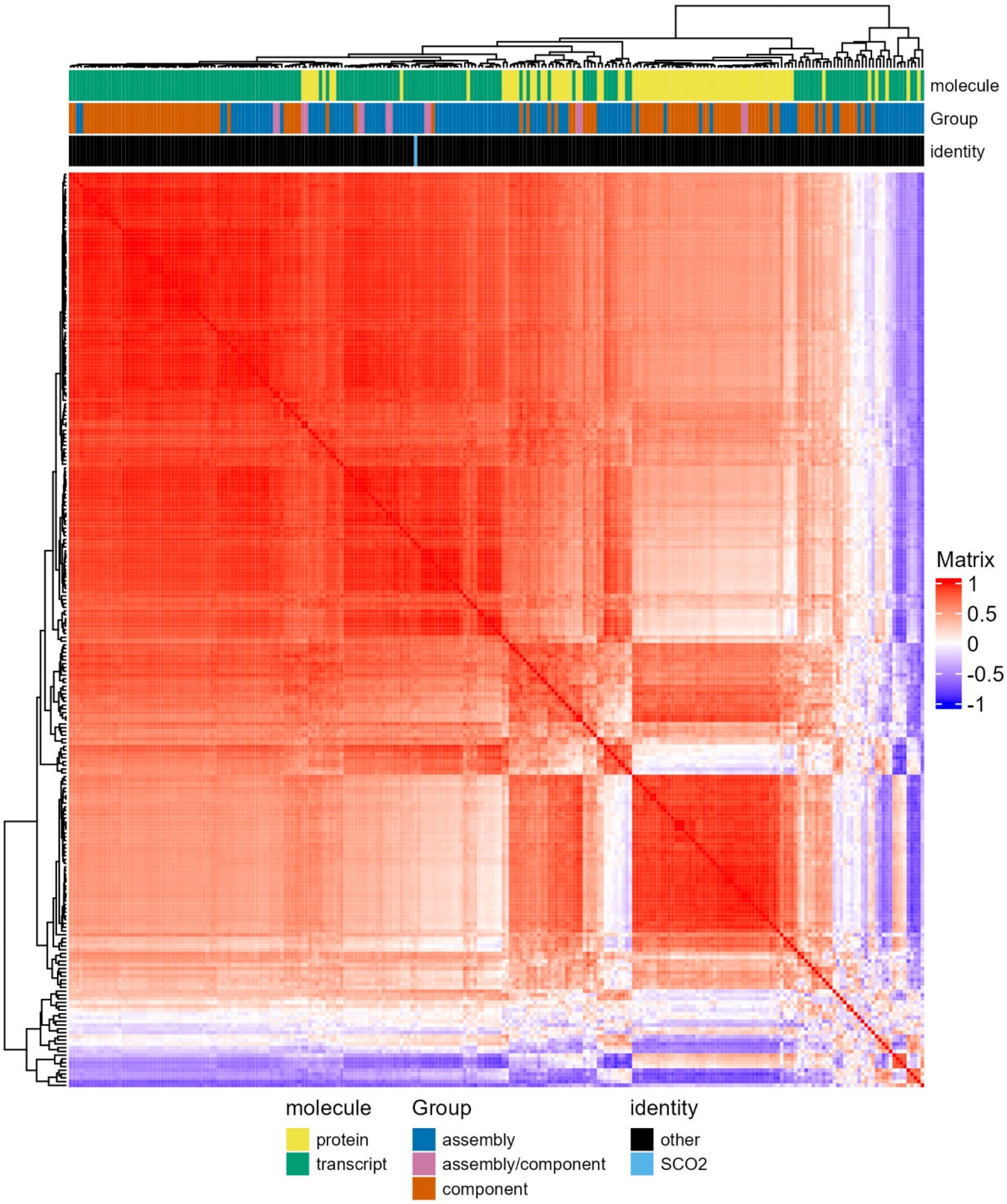

Supplemental Figure S 10: Heatmap of raw unfiltered correlation values of photosystem components and assembly factor transcripts and proteins. Annotation rows are colored according to molecule type, functional group and to highlight the identity of the SCO2 transcript.

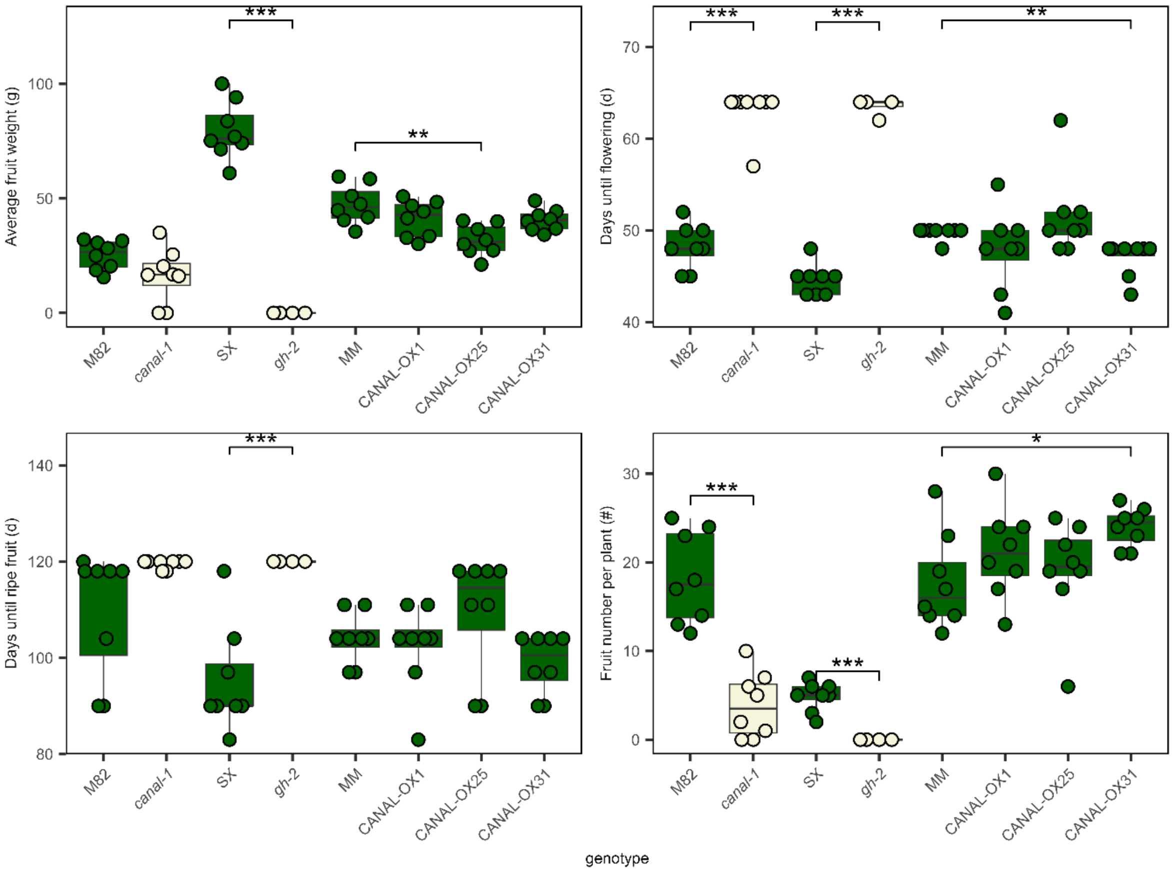

Supplemental Figure S 11: Phenotype Whole-plant traits of different genotypes in experiment including overexpression lines. Different panels show different traits. Statistical significance was estimated by pairwise-t-test. Significant differences are indicated by a bracket and asterisks (*: $p \leq 0.05$; **: $p \leq 0.005$; ***: $p \leq 0.0005$). g: gram, d: days, #: number.

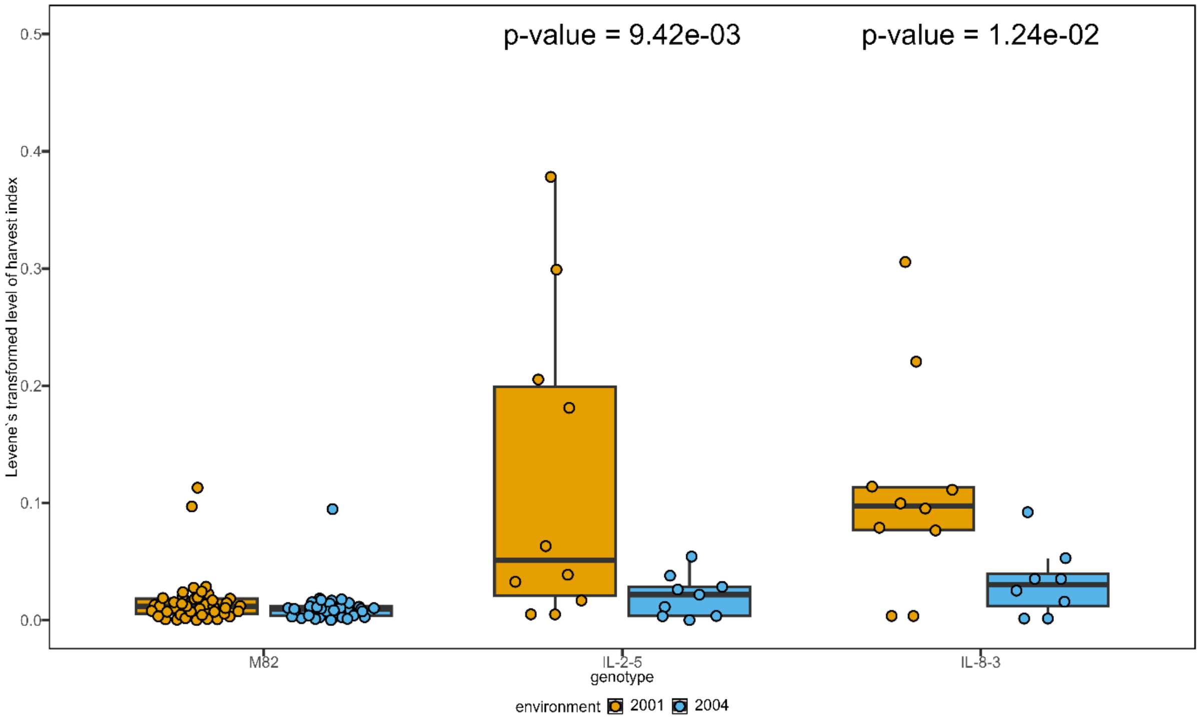

Supplemental Figure S 12: Levene`s-transformed level of harvest index (HI) in M82 wild type plants and ILs, from the seasons 2001 and 2004. P-value for GxE-effect was estimated via a mixed linear model as described previously (Alseekh et al., 2017).

```
                1                                                                                 82
CDS_SL4.0...   ATGTTGCCTCTGATAAATTCTTGTTCAATCTTCACTCCTCTAAGTTCTTCATTACTTCCTCCTCGTAGATCCACCGTCATTA
CDS_Spenn...   ATGTTGCCTCTGATAAATTCTTGTTCAATCTTCACTCCTCCAAGTTCTTCATTACTTCCTCCTCGTAGATCCACCGTCATTC

                83                                                                               164
CDS_SL4.0...   GTTGCCAAGCCGCTGCTGACCCATCAGCTGCCGGCGGACCCTCGTCCTTTCGCTGGTGGTTCAACTTCGGCGCCGCCGCCTC
CDS_Spenn...   GTTGCCAAGCCGCTGTTGACCCATCAGCTGCCGGCGGACCGTCGTCCTCTCGCTGGTGGTTCAACTTCGGCGCCGCCGCCTC

                165                                                                              246
CDS_SL4.0...   TGCTGCTGCTGCACCTGGGTTTGGGAGAAATAGCCAAAACTCCGACGAAGTTGCTGATAATGGAATCGGGAATAACATTAAG
CDS_Spenn...   TGCTGCTGCTGCACCTGGGTTTGGGAGAAATAGCCAAAACTCCGACGAAGTTGCTGATAATGGAATCGGGAATAACATTAAG

                247                                                                              328
CDS_SL4.0...   AAGAAGAAGACAAAAGTGAATGCGAAGGAAAGGCGTTGGTCGCGTAATAGAGAGAGTTATTTGGCTGATGATGGTGATGCTC
CDS_Spenn...   AAGAAGAAGACAAAAGTGAATGCGAAGGAAAGGCGTTGGTCGCGTAATAGAGAGAGTTATTTGGCTGATGATGGTGATGCTC

                329                                                                              410
CDS_SL4.0...   TTCCTCTTCCTATGACTTACCCTGACACTTCCCCTTGTTCCCCGGAGGAAATCGACCGCCGGCTGCAGTGTGATCCTATAAT
CDS_Spenn...   TTCCTCTTCCTATGACTTACCCTGACACTTCCCCTTGTTCCCCGGAGGAAATCGACCGCCGGCTGCAGTGTGATCCTATAAT

                411                                                                              492
CDS_SL4.0...   TGAGGATTGCAAGCAAGTTGTCTATGAGTGGACTGGAAAATGTCGGAGTTGCCAAGGGACAGGACTTGTCAGCTATTATAAC
CDS_Spenn...   TGAGGATTGCAAGCAAGTTGTCTATGAGTGGACTGGAAAATGTCGGAGTTGCCAAGGGACAGGACTTGTCAGCTATTATAAC

                493                                                                              574
CDS_SL4.0...   AAAAAGGGGAAAGAGACCATCTGTAAATGTATACCTTGTGCTGGAATCGGTTATGTGCAGAAAATAACACTACGCACGGATA
CDS_Spenn...   AAAAAGGGGAAAGAGACCATCTGTAAATGTATACCTTGCGCTGGAATCGGTTATGTGCAGAAAATAACACTACGCACGGATA

                575                                   612
CDS_SL4.0...   TTGATGTAATGGTGGACTTGGATGATAAACCACCTTAG
CDS_Spenn...   TTGATGTAATGGTGGACTTGGATGATAAACCACCTTAG
```

Supplemental Figure S 13: Alignment of SCO2 coding DNA sequence of Solanum lycopersicum (CDS_SL4.0) and Solanum pennellii (CDS_Spenn). Altered nucleobases are marked in red. Sequences were obtained from https://www.solgenomics.net/.

```
                1                                                                                82
AA_SL4.0 ...    MLPLINSCSIFTPLSSSLLPPRRSTVISCQAAADPSAAGGPSSFRWWFNFGAAASAAAAPGFGRNSQNSDEVADNGIGNNIK
AA_Spenn ...    MLPLINSCSIFTPPSSSLLPPRRSTVIRCQAAVDPSAAGGPSSSRWWFNFGAAASAAAAPGFGRNSQNSDEVADNGIGNNIK

.......................................................................................................

                83                                                                              164
AA_SL4.0 ...    KKKTKVNAKERRWSRNRESYLADDGDALPLPMTYPDTSPCSPEEIDRRLQCDPIIEDCKQVVYEWTGKCRSCQGTGLVSYYN
AA_Spenn ...    KKKTKVNAKERRWSRNRESYLADDGDALPLPMTYPDTSPCSPEEIDRRLQCDPIIEDCKQVVYEWTGKCRSCQGTGLVSYYN

.......................................................................................................

                165                                  203
AA_SL4.0 ...    KKGKETICKCIPCAGIGYVQKITLRTDIDVMVDLDDKPP
AA_Spenn ...    KKGKETICKCIPCAGIGYVQKITLRTDIDVMVDLDDKPP

.......................................................................................................
```

Supplemental Figure S 14: Alignment of SCO2 AA amino acid sequence of Solanum lycopersicum (AA_SL4.0) and Solanum pennellii (AA_Spenn). Altered amino acid bases are marked in red.
